# Interaction of Carbamoyl-Phosphate Synthase 1 with Agmatinase in the Liver of Torpid Bats

**DOI:** 10.1101/2025.09.26.678923

**Authors:** Yangyang Li, Chen-Chung Liao, Libiao Zhang, Junyan Lv, Shan Zheng, Ji Ma, Yiwen Wang, Bing Ni, Tianxiao Yang, Guimei He, Haipeng Li, Yi-Hsuan Pan

## Abstract

Mammalian torpor imposes unique metabolic constraints, yet the mechanisms of nitrogen metabolism during this state remain unclear. In this study, we show that the urea cycle is selectively regulated rather than broadly suppressed in torpid bats. A significantly increased abundance of carbamoyl-phosphate synthase 1 (CPS1) maintained its functional capacity during torpor and arousal in *Myotis ricketti*. Proteomic analyses and confocal microscopy identified a specific association and co-localization between CPS1 and agmatinase (AGMAT), an ATP-independent enzyme involved in nitrogen metabolism. Co-localization of CPS1 and AGMAT was also observed in torpid *Rhinolophus ferrumequinum*, a phylogenetically distant species. Fluorescence resonance energy transfer (FRET) further supported an indirect CPS1-AGMAT interaction. Most urea cycle enzymes exhibited stable or only moderately reduced expression during bat torpor, and metabolic profiling demonstrated sustained nitrogen flux. Together, these findings reveal a conserved adaptive mechanism that maintains urea cycle function, potentially enhancing osmotic stability and energy efficiency during prolonged fasting and water scarcity.

## Introduction

Hibernation is a physiological adaptation that allows mammals to survive extended periods of environmental stress, particularly cold and food scarcity. Before entering hibernation, many species accumulate white adipose tissue to serve as a primary energy reserve^1^. During winter, hibernators undergo repeated bouts of torpor and arousal. In torpor, small mammals such as bats can reduce their core body temperature to near-freezing levels while remaining viable^2,3^. This state is characterized by cessation of water intake and profound suppression of metabolic and cardiac activity. These changes are reversed quickly during spontaneous arousal, highlighting the reversible and regulated nature of the hibernation phenotype.

While lipid metabolism has been extensively studied and shown to support energy demands during torpor^4,5^, how nitrogen metabolism is modulated remains poorly defined. Prolonged fasting and dehydration during torpor pose significant challenges for nitrogen waste management. In a prior study, we reported that carbamoyl-phosphate synthase 1 (CPS1), the rate-limiting enzyme in the urea cycle, is upregulated during torpor in *Myotis ricketti* ^5,6^. This suggests that ammonia detoxification via the urea cycle may remain active during hypometabolic states.

The urea cycle is a highly conserved pathway responsible for converting toxic ammonia, generated by amino acid catabolism and gut microbial activity into urea, a process carried out by five core enzymes: carbamoyl-phosphate synthase 1 (CPS1), ornithine transcarbamylase (OTC), argininosuccinate synthase 1 (ASS1), argininosuccinate lyase (ASL), and arginase 1 (ARG1), along with two essential mitochondrial transporters, ornithine transporter 1 (ORNT1) and citrin^7,8^. CPS1 activity is allosterically activated by N-acetylglutamate (NAG), which is synthesized from glutamate and acetyl-CoA by N-acetylglutamate synthase (NAGS)^9^. However, how this pathway is regulated in the context of torpor and restricted water availability remains to be investigated, particularly under conditions of torpor and restricted water availability.

In this study, we examined CPS1 abundance, enzymatic activity, and protein interactions in two phylogenetically distant bat species, *Myotis ricketti* and *Rhinolophus ferrumequinum*. Through co-immunoprecipitation and mass spectrometry, we identified CPS1-interacting proteins, notably agmatinase (AGMAT), an ATP-independent enzyme involved in nitrogen metabolism. We validated their spatial association using confocal microscopy and confirmed their interaction proximity by fluorescence resonance energy transfer (FRET). Our results uncover a flexible, species-specific nitrogen metabolic program that maintains urea cycle functionality during torpor, likely contributing to osmotic and metabolic stability under conditions of dehydration and energy limitation.

## Materials and Methods

### Ethics Statement

Collection of animals and all experimental procedures were approved by the Animal Ethics Committee of East China Normal University (No. AE2012/03001). The bat species *Rhinolophus ferrumequinum* and *Myotis ricketti* investigated in this study are not listed as endangered. All animals were sacrificed by cervical dislocation, and ethical guidelines to minimize pain and suffering were strictly followed in accordance with the regulations of the Animal Ethics Committee of East China Normal University.

### Animal and Tissue Acquisition

Torpid *M. ricketti* bats and *R. ferrumequinum* bats were captured using hand nets from Fangshan Cave (39°70′N, 115°71′E), Beijing, China. The temperature was 17°C inside and 6.5°C outside the cave. As female bats are often pregnant during hibernation, only adult male bats were sampled in this study. Thirty male bats of each species were captured (Table S1), and ten bats of each species were sacrificed immediately upon capture to obtain liver tissues. Each of the remaining bats was placed separately in a cloth bag and transported to the lab, where the temperature was maintained at 28°C. Bats were spontaneously aroused during transportation. Ten bats from each species were sacrificed 2 h after arousal, and the remaining bats were sacrificed 24 h after arousal. Ten active bats of each species captured from the cave in summer were also sacrificed immediately upon capture.

Three 7-week-old C57BL/6J male mice, obtained from Shanghai Jihui Laboratory Animal Care Co., Ltd (Shanghai, China), were maintained in 12 h dark-light cycles at 22 ± 2°C with food and water provided ad libitum. The mice were sacrificed a week later to obtain liver tissues, which were flash-frozen in liquid nitrogen and stored at −80°C until used.

### Preparation of Liver Protein for Western Blotting

Each liver tissue (0.1 g) was homogenized in 1 ml lysis buffer (10% glycerol, 2% SDS, 1.25% β-mercaptoethanol, 12.5 mM EDTA, 25 mM Tris-HCl, pH 6.8, and 1/50 tablet of cOmplete™ EDTA-free protease inhibitor cocktail) with a Precellys^®^ 24 grinder (Bertin technologies, France). The homogenate was heated at 100°C for 10 min and centrifuged at 12,000 rpm, 4°C for 10 min. Proteins in the supernatant were separated by 10% SDS-PAGE and then transferred onto 0.2-μm PVDF membranes (Millipore, USA) with an electro-blotting apparatus (GHE4201, GenePure, Taiwan). The PVDF membranes were blocked in a solution containing 5% skim milk and 1% BSA at 4°C overnight and reacted with a series of primary antibodies (Supplementary Excel file 1) that were selected based on their ability to recognize conserved epitopes of target proteins of many mammalian species. After washing with TBST buffer (20 mM Tris-base, pH 7.6, 137 mM NaCl, and 0.1% Tween-20), blots were reacted with appropriate secondary antibodies and Immobilon^TM^ Western Chemiluminescent HRP Substrate WBKLSO500 (Millipore, USA). Images were captured using ImageQuant^TM^ LAS-4000 (Amersham Biosciences, USA), and the intensity of each detected band was quantified by ImageQuant^TM^ TL (v 7.0, Amersham Biosciences, USA). Equal loading of samples was monitored by Ponceau staining (Supplementary Information 1). The relative quantity of a protein was calculated by dividing the density value of the protein band on Western blot by the total density value of all protein bands in a lane of a Ponceau-stained blot^10,11^. Results are presented as mean ± SD from biological replicates (3 ≥ *n* ≤ 6), each with more than four technical replicates, and were analyzed using one-way ANOVA with the Holm-Sidak test or Student’s *t*-test, as appropriate for each analysis.

### Enzyme activity assay

The activity of carbamoyl-phosphate synthase 1 (CPS1) was determined as described previously^12^. For determination of CPS1 activity, each liver tissue (0.1 g) was homogenized in 1 ml lysis buffer (100 mM Tris-HCl, pH 7.5, 50 mM KCl, 0.5% Triton X-100, 1 mM EDTA, 1 mM DTT, and 1/50 tablet of cOmplete™ EDTA-free protease inhibitor cocktail) with a Precellys^®^ 24 grinder (Bertin technologies, France). The homogenate was centrifuged at 10,000 xg, 4°C for 10 min. A 0.2-ml aliquot of the supernatant was incubated with 0.8 ml reaction buffer (50 mM potassium phosphate, pH 7.5, 50 mM ammonium chloride, 50 mM sodium bicarbonate, 15 mM MgSO_4_, 10 mM L-ornithine, 10 mM ATP, 5 mM N-acetyl-L-glutamate, and 150U OTC) in a DNase- and RNase-free 1.5-ml Eppendorf tube and shaken on a shaker at 160 rpm, 37°C for 1 h. For negative controls, the reaction buffer without N-acetyl-L-glutamate and L-ornithine was used.

The reaction mixture (1.5 ml) containing citrulline generated by the enzyme was then centrifuged at 13,500 xg, 4°C for 15 min, and 0.1 ml of the supernatant was mixed with 0.6 ml of chromogenic reagent in a glass vial and heated at 100°C for 15 min in darkness. The chromogenic reagent was freshly prepared by mixing 100 ml solution A containing 0.4 g diacetyl monoxime (DAMO) and 7.5 g NaCl in ddH_2_O and 200 ml solution B containing 0.74 g antipyrine, 2.5 mg NH_4_Fe (SO_4_)_2_, 50 ml, 85% H_3_PO_4_, and 50 ml H_2_SO_4_ in ddH_2_O and kept at 4°C in darkness before being used. In the chromogenic reagent, DAMO is the chromophore for detection of citrulline. After cooling, 0.25 ml of the yellow solution was transferred to a well of a 96-well microtiter plate and measured for absorbance at a wavelength of 464 nm with a Synergy^TM^ HT spectrophotometer (Biotek, Winooski, VT) to obtain an OD_464_ value, which is positively correlated with the enzyme activity. The concentration of citrulline was determined against the citrulline standards (0.2-, 0.4-, 0.6-, 0.8-, 1-, 1.2-mM). *M. ricketti* and *R. ferrumequinum* bats of each state (3 ≥ *n* ≤ 7) were used in this assay, and at least three technical repeats were performed for each bat.

### Interactomics

#### Co-immunoprecipitation

Liver proteins were obtained from torpid, 24 h aroused, and active bats. Co-IP was performed using the Immunoprecipitation Kit from Thermo Fisher Scientific (catalog number 10006D) as previously described^5^. Briefly, Dynabeads^®^ were incubated with binding and washing buffer (0.01% Tween-20 in PBS, pH 7.4) containing a specific antibody (Ab) at room temperature for 20 min. Binding and washing buffer instead of a specific antibody was used as the negative control. The bead-Ab complexes were washed with the same buffer and then incubated with a protein sample containing the antigen (Ag) for 40 min at room temperature. The bead-Ab-Ag complexes were washed three times with washing buffer and then heated at 70°C for 10 min in denaturing solution (2% SDS, 1.25% β-mercaptoethanol, and 10% glycerol in 50 mM Tris-HCl, pH 6.8). The eluted proteins were fractionated by 10% SDS-PAGE, and the gel was stained for 20 min with Coomassie G-250^13^.

#### Recovery and identification of proteins

Each lane of the gel was cut into ten equal pieces after G-250 staining, and each gel piece was destained in a buffer containing 50% acetonitrile and 25 mM NH_4_HCO_3_ (1:1, v/v). After vacuum drying, each gel piece was rehydrated with 25 mM NH_4_HCO_3_ containing 1% β-mercaptoethanol and alkylated in the buffer containing 5% 4-vinylpyridine in 25 mM NH_4_HCO_3_ and 50% acetonitrile (1:1, v/v) for 20 min. Proteins in each gel piece were digested with trypsin (Promega, Mannheim, Germany) in 25 mM NH_4_HCO_3_ (1%, w/v) at 37°C overnight. The tryptic peptides were extracted from the gel piece with 25 mM NH_4_HCO_3_ for 10 min, dried, and stored at −20°C until used.

Tryptic peptides of a sample were dissolved in 10 μl formic acid (0.1%, v/v) and then loaded onto an online NanoAcquity UP LC system (Waters, Manchester, UK) connected to an LTQ-Orbitrap XL (Thermo Scientific, San Jose, CA). Solution A (0.1% formic acid in water) and solution B (0.1% formic acid in acetonitrile) were used as the mobile phases. Peptides loaded were first desalted in a trap column (ACQUITY UPLC Symmetry C18 100Å, 5 µm, 180 μm x 20 mm, Waters Corp.) and then chromatographed in a tip column (Acquity BEH C18 130Å, 1.7 µm, 100 μm × 100 mm, Waters Corp.) with 5 - 35% linear solution B gradient for 90 min and then with 35 - 95% solution B gradient for 10 min at a flow rate of 0.5 μl/min.

Eluted peptides were ionized with a spray voltage of 2 kV and then introduced into the mass spectrometer. Mass spectrometry data were recorded with the data dependent acquisition mode with an isolation width of 1.5 Da and then searched for six most intensely charged ions (2^+^ and 3^+^) with a full MS survey scan (m/z: 200-1500) at a resolution > 30,000 full width at half maximum (FWHM). Fragmented ions of each selected precursor were generated by collision-induced dissociation with helium gas at a relative collision energy of 35%.

The RAW data were first processed by the Xcalibur^TM^ software package (version 2.0.7 SR1, Thermo-Finnigan Inc., San Jose, CA), analyzed by PEAKS software (version 8.5, Bioinformatics Solutions Inc., Waterloo, Ontario, Canada), and then searched for best-matched peptides in the bat protein database^14^. The parameters used to identify proteins were as previously described^5^. All identified proteins were annotated with UniPort ID (Supplementary Excel file 2). Information on identified peptides and their charge states has been submitted to ProteomeXchange (https://doi.org/10.6019/PXD066236 containing 60 Raw, 60 Mgf, 1 Txt and 1 Xml files).

### Immunofluorescence

Each 2 mm^3^ cube of a liver tissue was embedded in Tissue Freezing Medium^®^ (Leica Biosystems, UK), frozen in liquid nitrogen, and sectioned with a cryostat. Each 7-μm-thick section was placed on a glass slide, fixed with 4% paraformaldehyde, and incubated with blocking solution (3% BSA and 0.3% Triton X-100 in PBS, pH 7.4) at room temperature for 2 h. Sections were then incubated with a primary antibody 24 h at 4°C, washed with PBS three times, followed by incubating with a secondary antibody labeled with Cy3 or Cy5 for 2 h in darkness at room temperature. The sections were washed again with PBS three times (Supplementary Excel file 1), mounted on glass slides with anti-fade mounting medium containing 4′, 6-diamidino-2-phenylindole (Beyotime Biotechnology, Shanghai, China) to stain nuclear DNA, and examined with a laser scanning Olympus FluoView FV10i confocal microscope (Tokyo, Japan). Double fluorescence from green (Cy3) and red (Cy5) channels was imaged with an excitation wavelength of 559 nm for Cy3 and 635 nm for Cy5. For determination of overlapping, images were analyzed using the JACoP Plugin of ImageJ. The intensity of each pixel in corresponding channels, Pearson’s correlation coefficient, and Manders’ correlation coefficient were determined for each image. Pearson’s correlation coefficient values of 0 and 1 represent no co-localization (random correlation) and complete co-localization (perfect correlation) of CPS1 and AGMAT, respectively. For Manders’ correlation coefficient, an M1 value indicates the proportion of overlapped AGMAT-CPS1 in AGMAT, and an M2 value indicates the proportion of AGMAT-CPS1 in CPS1. In Pearson’s or Manders’ correlation coefficient measurements^15^, a *P* value < 0.05 determined by one-way ANOVA with Holm-Sidak test was considered significant.

### Measurement of Fluorescence Resonance Energy Transfer (FRET)

The antibodies used for FRET experiments are listed in Supplementary Excel file 1. A modified FRET acceptor bleaching method^16^ with a conventional confocal laser scanning microscope (CLSM) TCS SP5 (Leica, Germany) was used. Before and after bleaching, images in Cy3 (donor) and Cy5 (acceptor) channels were acquired in line-by-line sequential mode with two automatic noise reduction steps. Cy5 was bleached 60 to 80 times with 120 x zoom and 100% power of a 633-nm HeNe laser line. The freehand region of interest (ROI) was outlined to measure FRET by determining the fluorescence intensity in Cy3 (donor) channel in bleached area before and after bleaching. The excitation wavelengths for Cy3 and Cy5 were 30% power of 559 nm and 5% power of 635 nm laser, respectively. The detection wavelength for Cy3 was 553 to 628 nm, and that for Cy5 was 643 to 731 nm. Six measurements were performed for each experimental condition.

### Targeted metabolomics

Concentrations of metabolites in urea cycle or polyamine metabolism (*i.e.*, L-citrulline, L-arginine, L-ornithine, agmatine, cadaverine, putrescine, spermidine, spermine, and S-adenosyl-L-methionine) of bats were determined by ultra-performance liquid chromatography-tandem mass spectrometry (UPLC-MS/MS, Shanghai Biotree biotech Co., LTD., China) with a multiple reaction monitoring mode (MRM) (Supplementary Excel file 3).

#### Preparation of standard solution

Standard compounds of various metabolites were purchased from CNW technologies (Dusseldorf, Germany). To establish a standard curve for each metabolite, standard solutions were prepared from the 10 mM stock solution of the compound by serial dilution (Supplementary Excel file 4). Limitations of detection and quantification of each metabolite were identified using standard compounds before analysis.

#### Sample preparation for metabolomics analysis

Each bat liver tissue listed in Supplementary Excel file 4 was weighted, placed in a 1.5-ml Eppendorf tube containing 3-mm stainless steel beads, and incubated in 200 μl of 80% acetonitrile in water. Each sample was vortexed for 15 s, homogenized with a grinder (JXFSTPRP-24, Jingxin Industrial Development Co., Ltd, Shanghai, China) at 4°C for 4 min, and then sonicated with a YM-080S ultra-sonicator at 40 kHz, 4°C for 5 min. After repeating the homogenization process three times, samples were stood at −40°C for 1 h and centrifuged at 13,800 xg, 4°C for 15 min; 100 μl of the clarified supernatant was then placed in a 1.5-ml Eppendorf tube for dansylation. Detection of dansylated polyamines was performed as previously described^17^. Briefly, the supernatant (or standard solution) was mixed with 50 μl of 20 mg/ml dansyl chloride in acetonitrile and 50 μl of 0.1 M sodium bicarbonate. After 1 h incubation in darkness at 40°C, the sample was mixed with 50 μl of 0.1% formic acid in water and vortexed for 15 s, followed by centrifugation at 13,800 xg, 4°C for 10 min. An 80-μl aliquot of the supernatant was transferred to a vial in the auto-sampler tray and maintained at 4°C.

#### UPLC-MRM-MS/MS Analysis

A 2-μl aliquot of a sample was introduced into an online Agilent 1290 Infinity II series UPLC System (Agilent Technologies) connected to an Agilent 6460 triple quadrupole mass spectrometer (Agilent Technologies). Solution A (10 mM ammonium formate and 0.1% formic acid in water) and solution B (100% acetonitrile) were used as mobile phases. Loaded sample was chromatographed at 35°C in a Waters Acquity UPLC HSS T3 analytical column (1.8 μm, 100 μm x 2.1 mm) with solution B gradient of 25% for 30 s, 25 - 98% for 5.4 min, 98% for 3.5 min, 98 - 25% for 6 s, and 25% for 2.5 min at a flow rate of 0.4 ml/min.

Eluted analytes that were ionized by an Agilent Jet Stream thermal gradient focusing electrospray ionization (AJS-ESI) source, introduced into an Agilent 6460 triple quadrupole mass spectrometer, and examined with the following parameters: capillary voltage, +4000/-3500 V; nozzle voltage, +500/-500 V; gas, 300°C nitrogen at a flow rate of 5 L/min; sheath gas, 300°C nitrogen at a flow rate of 11 L/min; nebulizer, 45 psi. Mass spectrometry data were acquired and processed with the MRM scan mode using the MassHunter workstation software (B.08.00, Agilent Technologies). Flow injection analysis was used to optimize MRM parameters for each analyte. Several most sensitive transition Q1/Q3 pairs were scanned by MS1/MS2 and used to optimize the collision energy for each ion pair. Among the optimized MRM transitions per analyte, the Q1/Q3 pair (Supplementary Excel file 4) with the highest sensitivity and selectivity was selected as the quantifier for quantitation, and the other transition pairs were used as qualifiers to verify the identity of target analytes.

## Results

### Expression, activity, and interacting partners of CPS1 in torpid Myotis ricketti bats

To confirm the elevated hepatic expression of CPS1 in torpid *Myotis ricketti* bats compared with summer-active individuals^5,6^, we quantified CPS1 protein levels across physiological states by Western blotting (Fig. 1A). CPS1 abundance was significantly higher in torpid bats (*P* < 0.0001) and remained elevated at 2 hours after arousal, with a slight decline by 24 hours post-arousal (Fig. 1B). Overall, CPS1 expression in torpid and aroused bats was significantly higher than in summer-active bats (Fig. 1B, *P* < 0.0001). However, the total amount of citrulline, primarily produced through the urea cycle, remained comparable among 2-hour, 24-hour arousal, and summer-active states, despite reduced CPS1 activity per enzyme unit during torpor and arousal (Fig. S1A), suggesting dynamic modulation independent of protein abundance. These findings indicate that CPS1 is differentially regulated at both the protein and functional levels, potentially supporting urea cycle activity throughout torpor and rewarming.

**Figure 1.**
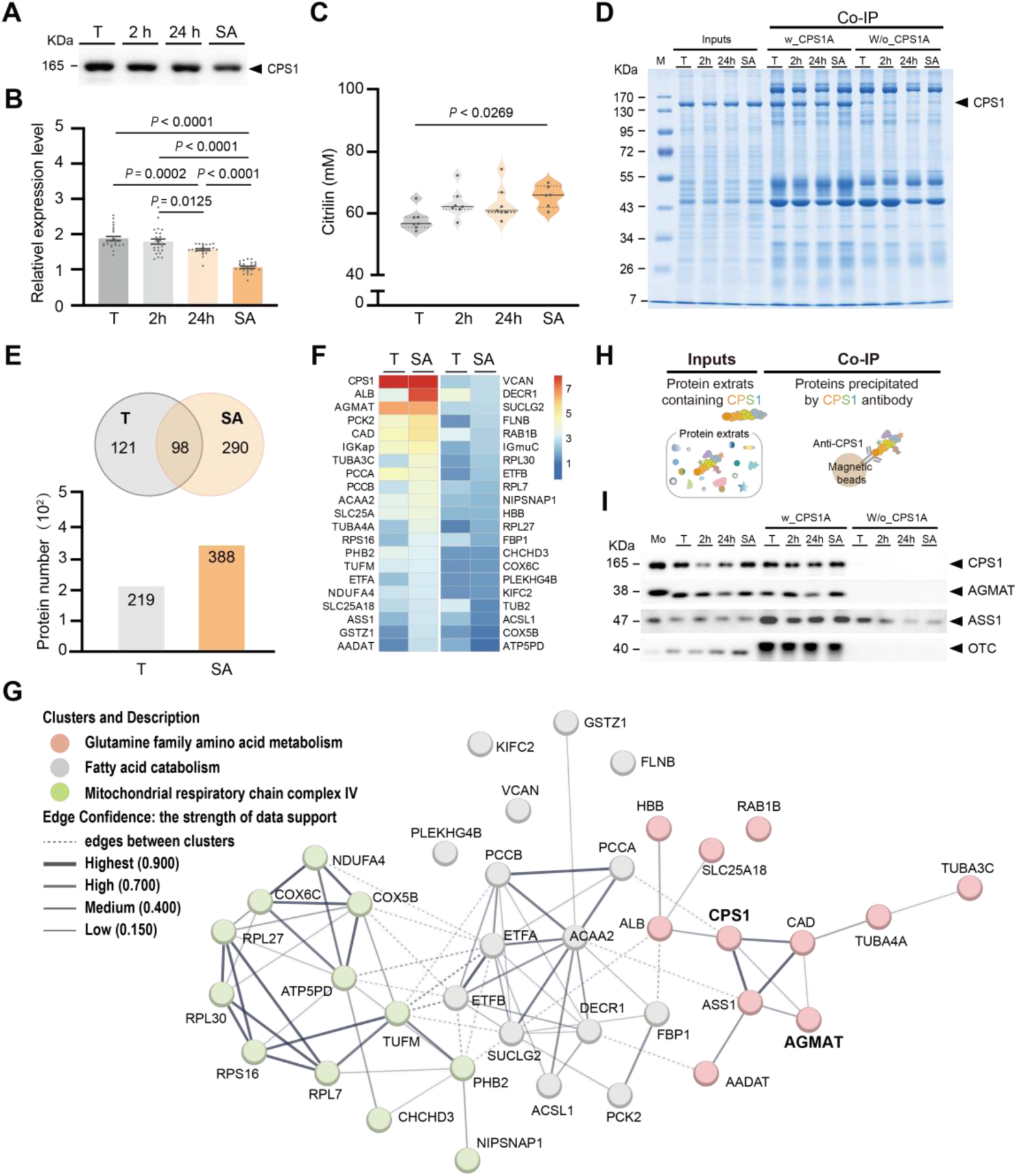
Abundance, activity, and interacting partners of CPS1. **(A)** Western blotting and **(B)** relative hepatic abundance of CPS1 in *M. ricketti.* **(C)** Citrulline (mM) produced by CPS1 across different physiological states. Data are presented as mean ± SE and analyzed using one-way ANOVA with Tukey’s multiple comparisons test. A *P* < 0.05 was considered significant. **(D)** SDS-PAGE of CPS1 co-immunoprecipitation (Co-IP). M, marker; Inputs, total protein; CPS1 antibody with or without CPS1 binding. **(E)** Venn diagram and histogram showing the number of CPS1-interacting proteins detected in torpid and summer-active states. **(F)** Heatmap of nonredundant CPS1-interacting proteins identified in both torpid and summer-active bats. **(G)** Protein-protein interaction network of the proteins shown in panel (F), analyzed by STRING and constructed using the k-means algorithm. **(H)** Schematic and **(I)** Western blot validation of AGMAT, ASS1, and OTC co-precipitated with CPS1. Mo, mouse liver sample; T, torpor; 2h, 2 hours after arousal; 24h, 24 hours after arousal; SA, summer-active state.

To identify CPS1-interacting proteins, we performed co-immunoprecipitation (Co-IP). Protein complexes precipitated with or without CPS1 antibody were resolved by SDS-PAGE (Fig. 1D) and subsequently analyzed by mass spectrometry. In total, 219 CPS1-associated proteins were identified specifically in torpid bats, 388 in summer-active bats, and 98 were shared between both physiological states (Fig. 1E and Supplementary Excel File 2). Associations between CPS1 and its interacting proteins were analyzed and visualized using a heatmap (Fig. 1F) and STRING (Fig 1G). The interactions between CPS1 and AGMAT, ASS1 or OTC^18,19^ (Supplementary Excel File 5) were further validated using Co-IP protein samples (Fig. 1H and 1I). These results suggest that CPS1 regulates the urea cycle through its incorporation into nitrogen-metabolism protein complexes.

### Co-localization and interaction analysis of CPS1 and AGMAT in hibernating bats

To investigate whether CPS1 and AGMAT are spatially associated, we examined their localization in hepatic tissues of *M. ricketti* and *R. ferrumequinum* using confocal microscopy. Liver tissues from mice maintained under standard laboratory conditions were used as a control. Co-localization of CPS1 and AGMAT was observed in both torpid bat species but not in mice (Fig. 2A and Supplementary Information 2), suggesting that this spatial association is a conserved feature in hibernating bats.

**Figure 2.**
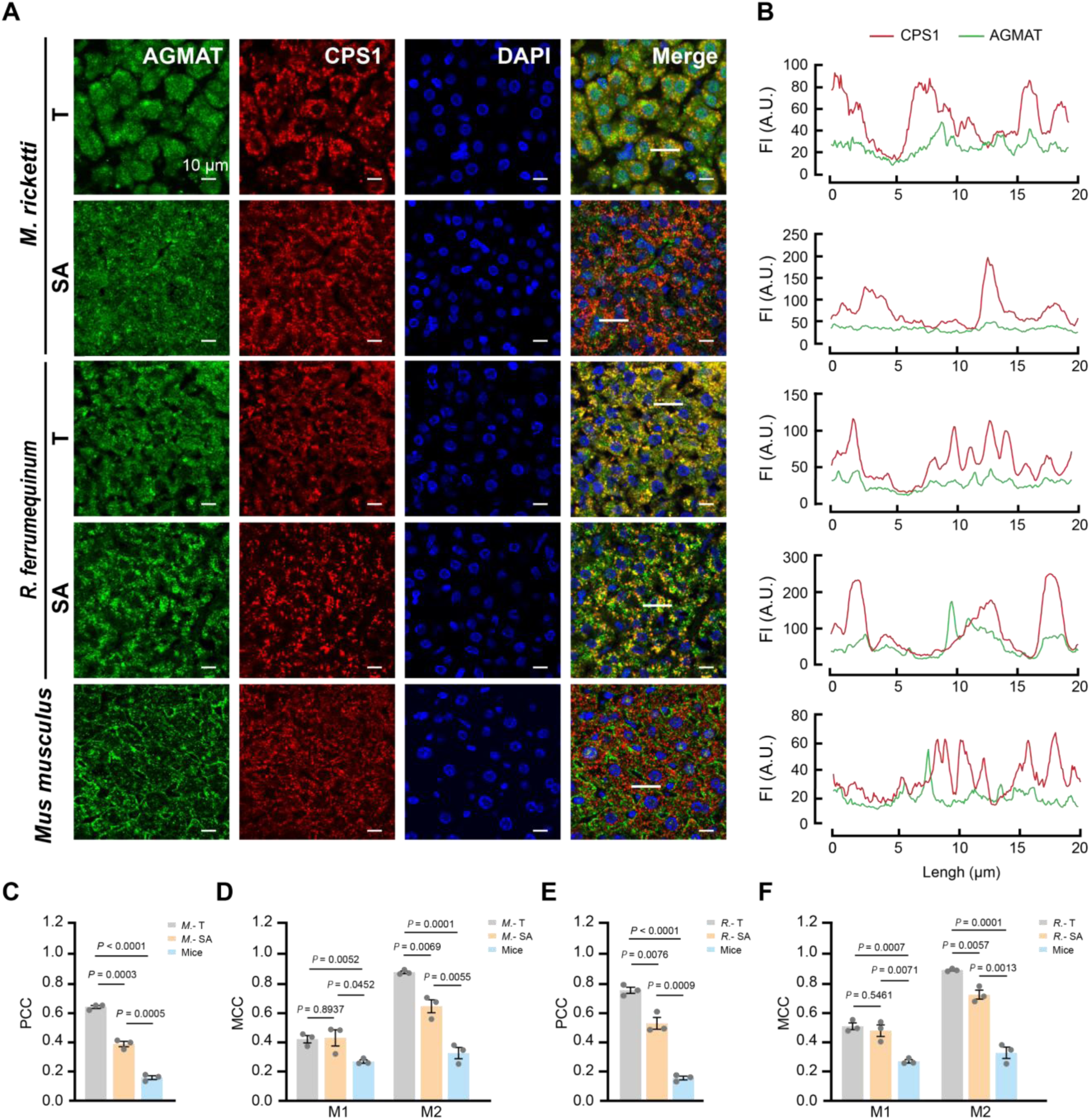
Co-localization and interaction analysis of CPS1 and AGMAT. **(A)** Immunofluorescence images of AGMAT (green), CPS1 (red), nuclei (blue), and overlap (yellow). Scale bar, 10 μm. **(B)** Fluorescence intensity (A.U.) of CPS1 and AGMAT along the white lines indicated in the merged panels of (A), plotted as red and green traces, respectively. **(C)** Quantification of co-localization using Pearson’s correlation coefficient (PCC) and Manders’ correlation coefficient (MCC) from the merged images in (A). M1, fraction of AGMAT overlapping CPS1; M2, fraction of CPS1 overlapping AGMAT. Data are obtained from three bats and three mice. T, torpor; SA, summer-active state. Results are presented as mean ± SEM and analyzed using a two-tailed Student’s *t*-test. A *P* value < 0.05 is considered statistically significant.

In *M. ricketti*, fluorescence intensity (FI) and the co-localization coefficient were quantified (Fig. 2B)^15^. Pearson’s correlation coefficient was significantly higher in torpid bats than in summer-active individuals (0.62 vs 0.43; *P* = 0.0003), with the lowest value observed in mice (Fig. 2C). Manders’ coefficient analysis yielded consistent results. More than 70% of CPS1 fluorescence overlapped with AGMAT in torpid bats, and the extent of co-localization was significantly greater than that in summer-active bats (Fig. 2D). These data indicate enhanced co-localization of CPS1 and AGMAT in torpid *M. ricketti*.

A similar pattern was observed in *R. ferrumequinum*. Pearson’s correlation coefficient was significantly higher in torpid bats compared to summer-active individuals (0.73 vs 0.52; *P* = 0.0076) (Fig. 2E), and the overlap fraction between CPS1 and AGMAT signals was also greater in torpid bats than in both summer-active bats and mice (0.90 vs 0.77; *P* = 0.0057) (Fig. 2F). These results confirm that CPS1 and AGMAT are co-localized in both bat species during torpor, supporting a potential functional association related to nitrogen metabolism.

To assess whether the observed interaction is direct, we performed fluorescence resonance energy transfer (FRET) analysis by measuring the increase in AGMAT fluorescence (ΔIF). FRET occurs only when the distance between two proteins is less than 10 nm^20^, typically corresponding to a FRET efficiency greater than 0.08. Although FRET efficiencies in both torpid and summer-active bats were significantly higher than those in mice, the mean values below 0.08, implying that the interaction between CPS1 and AGMAT is likely indirect (Fig. S2), possibly mediated through a nitrogen metabolism-associated protein complex.

### Selective regulation of nitrogen metabolism supports urea cycle activity during hibernation

To investigate nitrogen metabolism during torpor, we examined hepatic abundance of urea cycle and other nitrogen-related enzymes in *Myotis ricketti* (Fig. 3A). Western blotting showed significantly upregulation of NAGS, SIRT5, and CPS1 (*P* < 0.0001) during bat torpor (Fig. 3B and 3C), while ASL and ARG1 remained unchanged (Fig. 3D and 3E). Meanwhile, the abundance of OTC, ASS1, AGMAT, and ODC showed a moderate reduction relative to summer-active bats (Fig. 3D-G). These results indicate that nitrogen metabolism is selectively regulated rather than globally suppressed, preserving urea cycle function under hypometabolic conditions.

**Figure 3.**
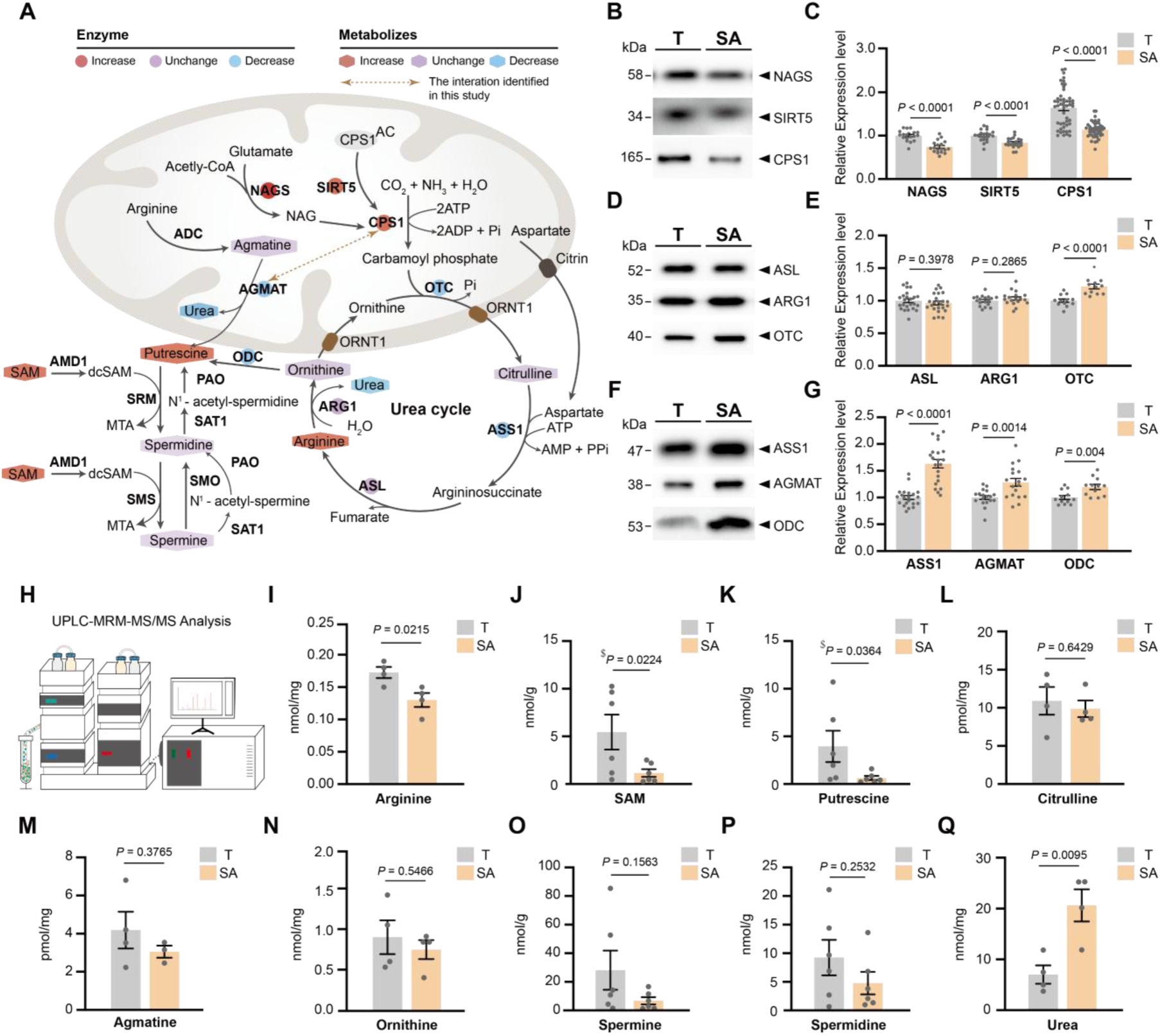
Abundance of enzymes and metabolites involved in urea and polyamine metabolism in *M. ricketti*. **(A)** Schematic of the urea cycle and polyamine biosynthesis. NAGS, N-acetylglutamate synthase; ADC, arginine decarboxylase; AGMAT, agmatinase; CPS1, carbamoyl phosphate synthetase 1; OTC, ornithine transcarbamylase; ASS1, argininosuccinate synthase 1; ASL, argininosuccinate lyase; ARG1, arginase 1; ODC, ornithine decarboxylase; SAM, S-adenosyl-L-methionine; AMD1, SAM decarboxylase; dcSAM, decarboxylated SAM; MTA, 5’-methylthioadenosine; SMS, spermine synthase; SRM, spermidine synthase; SMO, spermine oxidase; SAT1, spermidine/spermine N1-acetyltransferase 1; PAO, polyamine oxidase (also known as APAO, acetylpolyamine oxidase). **(B, D, F)** Western blotting and **(C, E, G)** relative protein expression of NAGS, SIRT5, CPS1 (B, C); ASL, ARG1, OTC (D, E); and ASS1, AGMAT, ODC (F, G). **(H)** Schematic of UPLC-MRM-MS/MS analysis. Concentrations of **(I)** arginine, **(J)** SAM, **(K)** putrescine, **(L)** citrulline, **(M)** agmatine, **(N)** ornithine, **(O)** spermine, **(P)** spermidine, and **(Q)** urea. T, torpor; SA, summer-active state. Results are presented as mean ± SEM (n = 3-6) and analyzed with a two-tailed Student’s *t*-test. A *P* value < 0.05 is considered statistically significant.

To evaluate physiological outcomes, we quantified hepatic metabolites (Fig. 3H). Arginine, S-adenosyl-L-methionine (SAM), and putrescine were elevated in torpid bats (Fig. 3I-K), whereas citrulline, agmatine, ornithine, spermine, and spermidine remained stable across states (Fig. 3L-P). Urea levels declined during torpor (Fig. 3Q). Together, these results suggest that nitrogen metabolism is modulated in a state-dependent manner, preserving balance rather than being broadly suppressed during hibernation.

To test whether this regulatory pattern is conserved, we analyzed *Rhinolophus ferrumequinum*. Immunoblotting showed that most enzymes exhibited similar abundance between torpid and summer-active states (Fig. 4A and 4B), with ASS1, AGMAT, and ODC displaying modest decreases during torpor. Notably, the specific activity of CPS1 (Fig. 4E) and metabolite levels remained comparable across states (Fig. 4L-M). These data suggest that nitrogen metabolism in torpor is governed by slightly different strategies across species but ultimately supports homeostasis during prolonged fasting and limited water availability.

**Figure 4.**
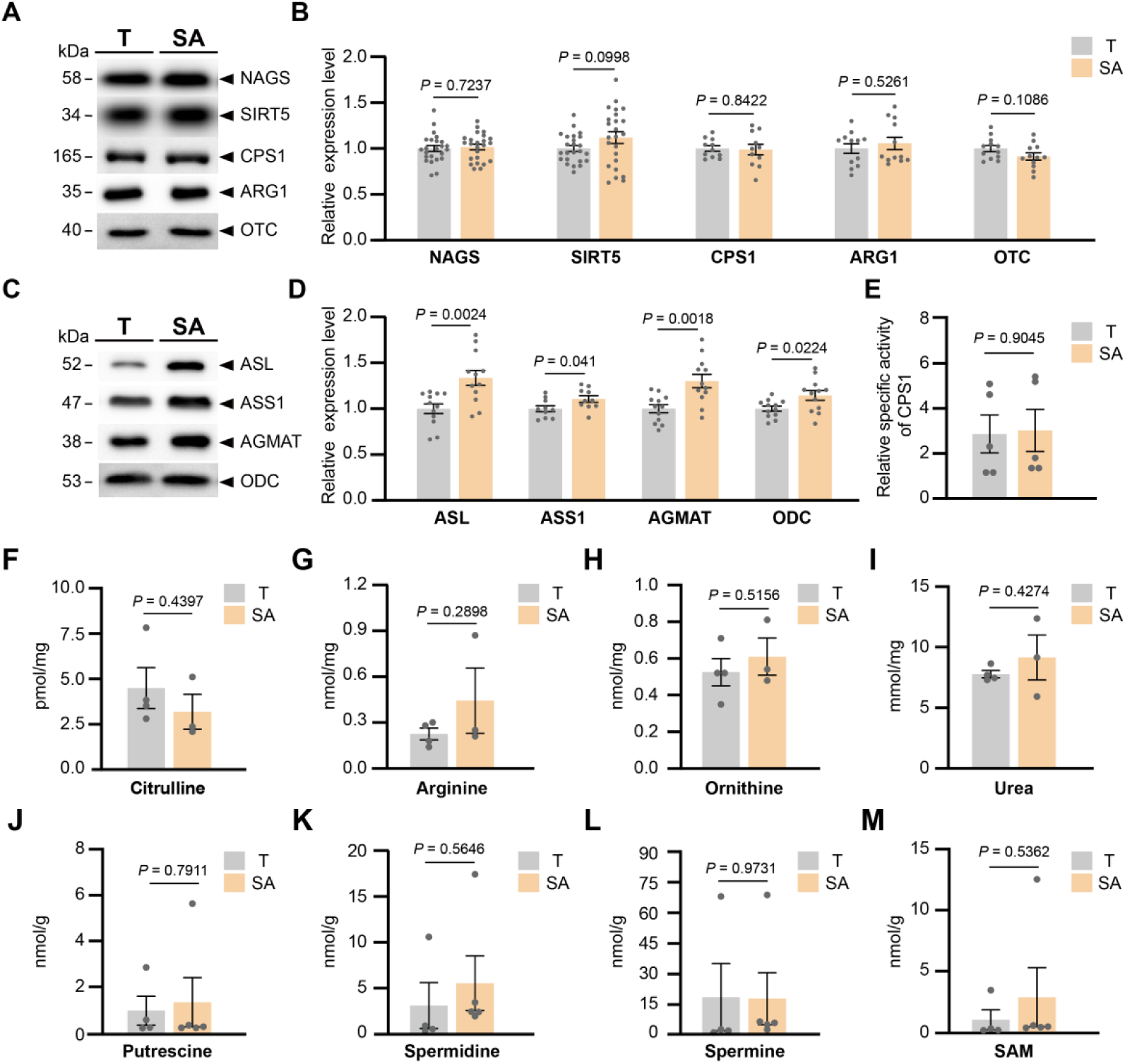
Abundance of enzymes and metabolites involved in urea and polyamine metabolism in *R. ferrumequinum*. **(A, C)** Western blotting and **(B, D)** relative protein expression levels of NAGS, SIRT5, CPS1, ARG1, OTC (A, B), and ASL, ASS1, AGMAT, ODC (C, D). **(E)** Relative CPS1 activity per unit of enzyme. Concentrations of **(F)** citrulline, **(G)** arginine, **(H)** ornithine, **(I)** urea, **(J)** putrescine, **(K)** spermidine, **(L)** spermine, **(M)** S-adenosyl-L-methionine, SAM. T, torpor; SA, summer-active state. Results are presented as mean ± SEM (n = 3-6) and analyzed with a two-tailed Student’s *t*-test. A *P* value < 0.05 is considered statistically significant.

## Discussion

Hibernation imposes prolonged fasting, dehydration, and hypothermia, yet nitrogen homeostasis is maintained with minimal toxicity. In ground squirrels, torpor is accompanied by nitrogen buffering and recycling, with amino acids and the γ-glutamyl system serving as temporary nitrogen sinks and substantial flux occurring during interbout arousals.^21^ Transcriptomic studies in American black bears have likewise revealed coordinated downregulation of urea-cycle genes in winter, suggesting a seasonal throttling of canonical ureagenesis.^22^ Our data in two phylogenetically distant bats, *M. ricketti* and *R. ferrumequinum*, refine this view: the urea cycle is not broadly suppressed but selectively tuned through differential enzyme abundance, post-translational regulation, and protein-protein association, thereby preserving ammonia detoxification capacity during torpor and rewarming.

A central observation is the dissociation between CPS1 abundance and activity in *M. ricketti*: protein levels rose sharply in torpor, while catalytic activity decreased modestly and rebounded on arousal. This pattern is compatible with known post-translational control of CPS1. SIRT5 deacetylates and activates CPS1 in liver mitochondria in response to fasting-induced changes in the NAD/NADH ratio, and SIRT5 deficiency blunts CPS1 activation with resultant hyperammonemia.^23^ Elevated NAGS in torpor is also notable because N-acetylglutamate is the obligatory allosteric activator of CPS1; variation in NAGS level or activity alters CPS1 effective capacity at a given protein abundance.^24^ Together with the increase of SIRT5, these findings support a model in which bats maintain a “poised” CPS1 pool during torpor, high protein capacity with tunable activity, allowing safe ammonia handling at low temperature and rapid upshift during arousal. Species differences between *M. ricketti* and *R. ferrumequinum* in NAGS and SIRT5 changes suggest multiple regulatory solutions converging on preserved urea-cycle function under energy and water limitation.

A second advance is the CPS1-AGMAT association we identified in torpid liver. Agmatinase hydrolyzes agmatine to putrescine while releasing urea, linking polyamine turnover to nitrogen disposal.^25^ Confocal co-localization in both bat species, together with Co-IP and FRET distances that argue for indirect rather than tight binary binding, point to a temperature-tolerant protein assembly involved in nitrogen metabolism rather than a fixed heterodimer. Such mesoscale organization could reduce substrate diffusion distances or coordinate local microenvironments for ureagenesis and polyamine interconversion when reaction rates are depressed by hypothermia. The functional plausibility of this adaptation is supported by broader literature highlighting polyamine metabolism as a key regulator of stress responses, proteostasis, and longevity.^26,27^

Enzyme and metabolite profiles support selective rather than global regulation. In *M. ricketti*, NAGS, SIRT5, and CPS1 increased, ASL and ARG1 were stable, and OTC, ASS1, AGMAT, and ODC decreased moderately; metabolites such as arginine and putrescine rose, whereas citrulline, ornithine, spermine, and spermidine were maintained. In *R. ferrumequinum*, most enzymes and metabolites changed little. This plasticity fits a broader hibernation paradigm in which nitrogen is conserved and redistributed rather than excreted wholesale, and where flux may be shifted between ureagenesis and alternative fates depending on species ecology and torpor depth.^21,22^ The maintenance of key intermediates despite reduced urea output is consistent with regulated throttling of the urea cycle to balance detoxification with osmotic economy.

Water balance is integral to nitrogen management in hibernators. Renal filtration and plasma flow decline during torpor, concentrating urine and limiting water loss, while the kidney undergoes structural and functional adjustments across torpor-arousal cycles.^28^ Our observation of markedly distended, fluid-filled bladders (grape-shaped bladder) in torpid *M. ricketti* but not in *R. ferrumequinum* (Fig. S3, Supplementary Information 3) hints at species-specific differences in bladder storage or urea handling that may complement hepatic regulation. Although we did not directly assess urea recycling, nitrogen salvage and urea retention are increasingly recognized as strategies that buffer ammonia toxicity while conserving water in hibernators.^29,30^ Future studies should extend these findings to the cellular level, where controlled low-temperature systems could be used to dissect how CPS1-AGMAT associations influence mitochondrial nitrogen flux and polyamine balance under hypometabolic conditions. Such approaches may help clarify whether these protein assemblies represent a generalizable strategy of cold-adapted nitrogen regulation across small hibernating mammals.

In summary, bats appear to preserve ammonia-detoxifying capacity during torpor through a triad of mechanisms: maintaining a large but post-translationally tunable CPS1 pool, assembling nitrogen-metabolism proteins that include AGMAT, and coordinating hepatic processes with renal and bladder water-conservation strategies. These results integrate with and refine existing models of hibernation nitrogen economy by emphasizing selective control rather than global shutdown of ureagenesis. A deeper understanding of these controls may inform medical approaches to hypothermia, organ preservation, and acute hyperammonemia management.

## Abbreviations

CPS1: Carbamoyl-phosphate synthase 1
AGMAT: Agmatinase
FRET: Fluorescence resonance energy transfer
UPLC-MS/MS: Ultra-performance liquid-chromatography tandem mass spectrometry

## Acknowledgments

We thank Dr. Chao-Hung Lee for editing the manuscript and providing valuable advice and the ECNU Multifunctional Platform for Innovation (010, 011 and 004) for their mouse breeding service. This work was supported by the National Natural Science Foundation of China (31100273, to YHP) and the Eastern Talent Plan Leading Project (to HL).

## Author Contributions

Y.L., C.L., J.L., Y.P., S.Z., and T.Y. performed experiments. Y.L., C.L., Y.W., B.N., and G.H. collected data. Y.L. and Y.P. conducted statistical and bioinformatics analyses, L.Z. assisted in field work. Y.P., G.H., and H.L. provided experimental materials. Y.P. designed the study. Y.L., G.H., H.L. and Y.P. wrote the manuscript.

## Supplementary Figures

**Figure S1.**
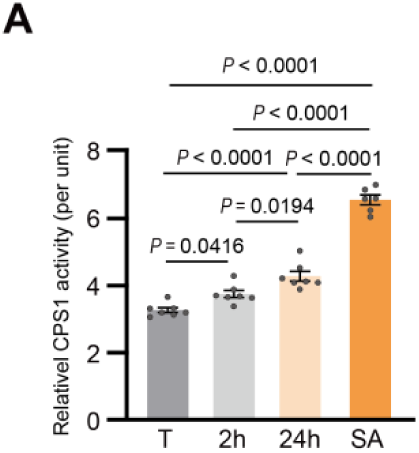
CPS1 activity of *M. ricketti.* **(A)** Relative enzyme activity per unit of CPS1. Data are presented as mean ± SE and analyzed using one-way ANOVA with Tukey’s multiple comparisons test. A *P* < 0.05 was considered significant. T, torpor; 2h, 2 hours after arousal; 24h, 24 hours after arousal; SA, summer-active state.

**Figure S2.**
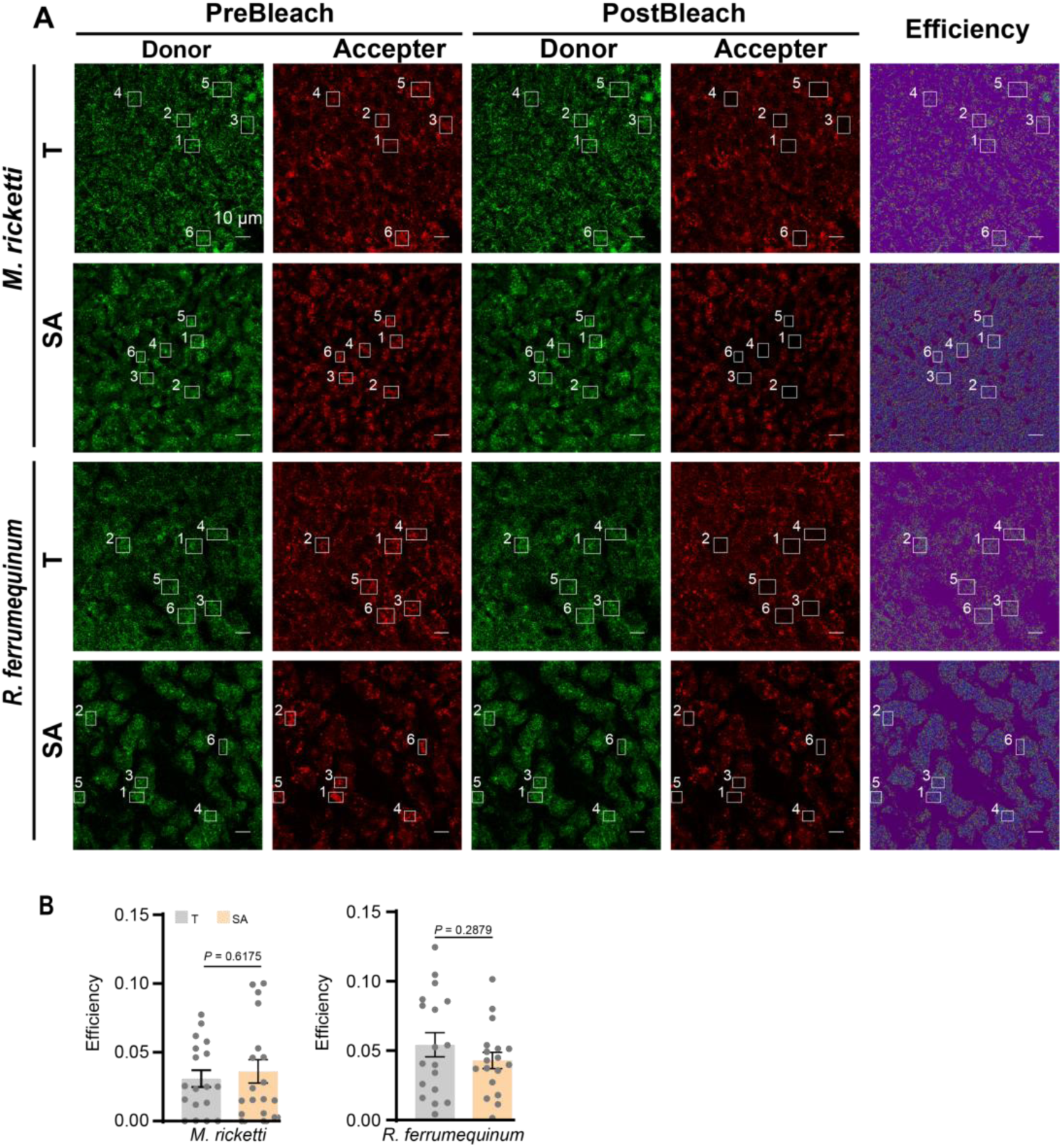
Fluorescence images of liver sections used in FRET analysis. **(A)** Donor and acceptor are labeled with Cy3 (green) and Cy5 (red)-conjugated secondary antibodies, respectively. The acceptor (Cy5) is bleached in the region of interest (ROI). **(B)** ROI efficiency in *M. ricketti* and *R. ferrumequinum* is calculated and presented as mean ± SEM (n = 3) and analyzed using a two-tailed Student’s *t*-test. A *P* value < 0.05 is considered statistically significant. Scale bar, 10 μm.

**Figure S3.**
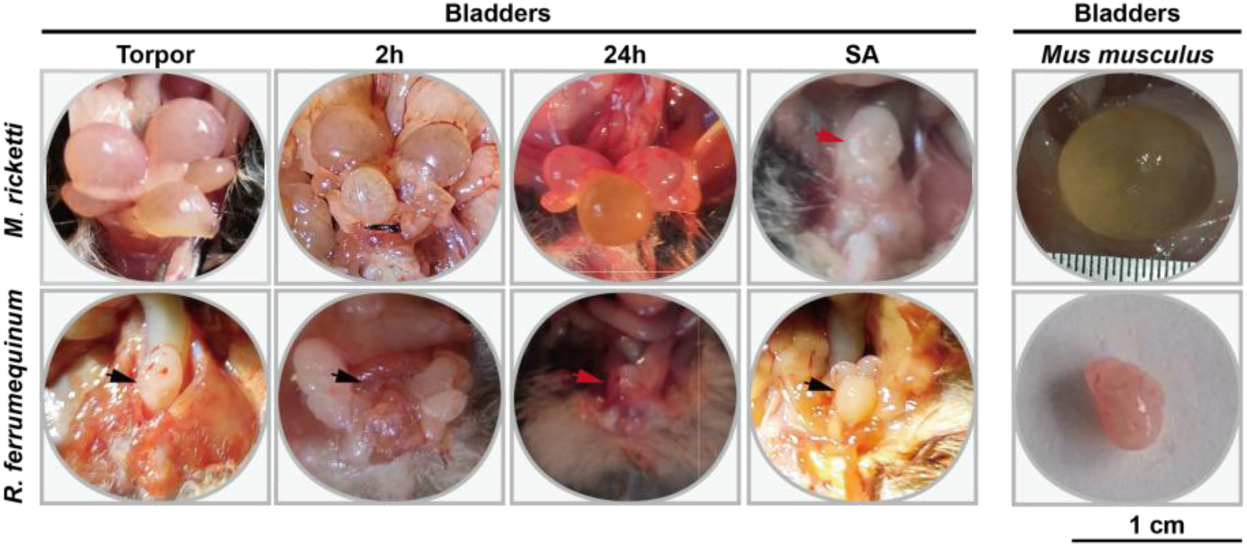
Representative images of grape-shaped bladders observed in torpid and aroused *M. ricketti*, but not in *R. ferrumequinum* or mice.

