## Supplementary Information 1 for "Interaction of Carbamoyl-Phosphate Synthase 1 with Agmatinase in the Liver of Torpid Bats"

### CPS1

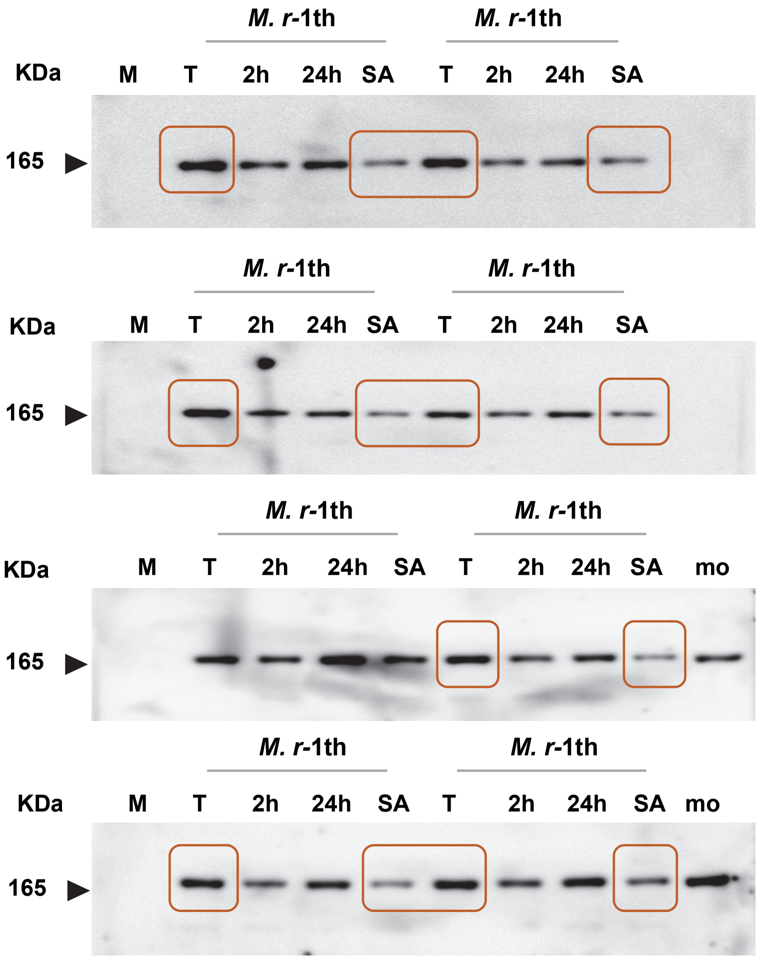
M. r-1th
M. r-1th
KDa
M
T
2h
24h
SA
T
2h
24h
SA
165
M. r-1th
M. r-1th
KDa
M
T
2h
24h
SA
T
2h
24h
SA
165
M. r-1th
M. r-1th
KDa
M
T
2h
24h
SA
T
2h
24h
SA
mo
165
M. r-1th
M. r-1th
KDa
M
T
2h
24h
SA
T
2h
24h
SA
mo
165

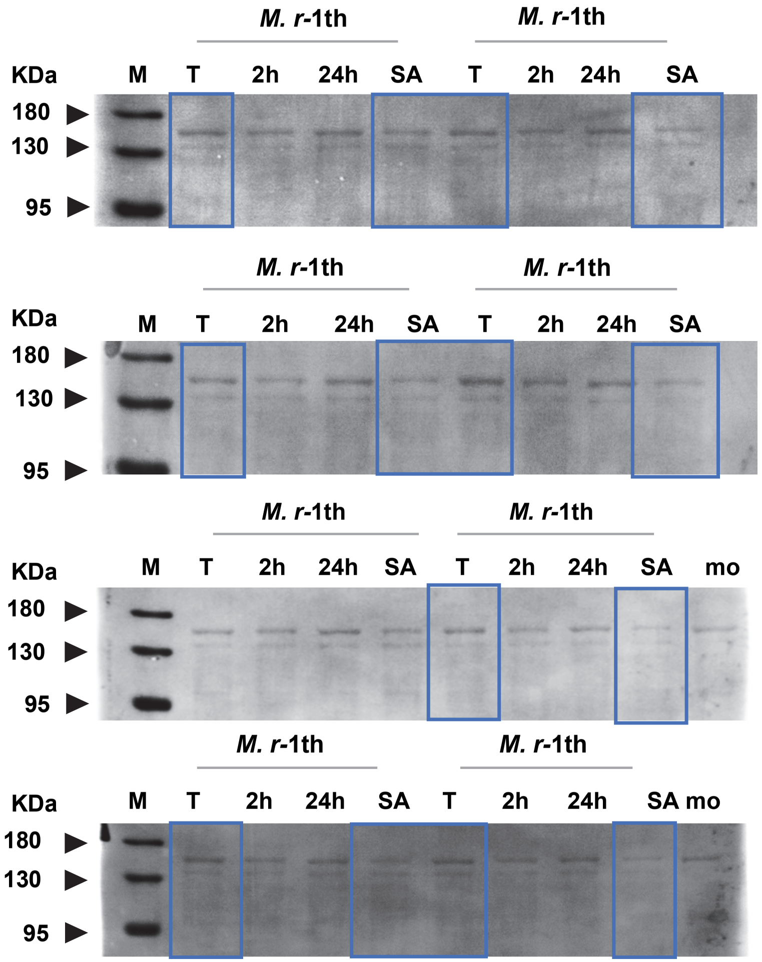
M. r-1th
M. r-1th
KDa
M
T
2h
24h
SA
T
2h
24h
SA
180
130
95
M. r-1th
M. r-1th
KDa
M
T
2h
24h
SA
T
2h
24h
SA
180
130
95
M. r-1th
M. r-1th
KDa
M
T
2h
24h
SA
T
2h
24h
SA
mo
180
130
95
M. r-1th
M. r-1th
KDa
M
T
2h
24h
SA
T
2h
24h
SA mo
180
130
95

### CPS1

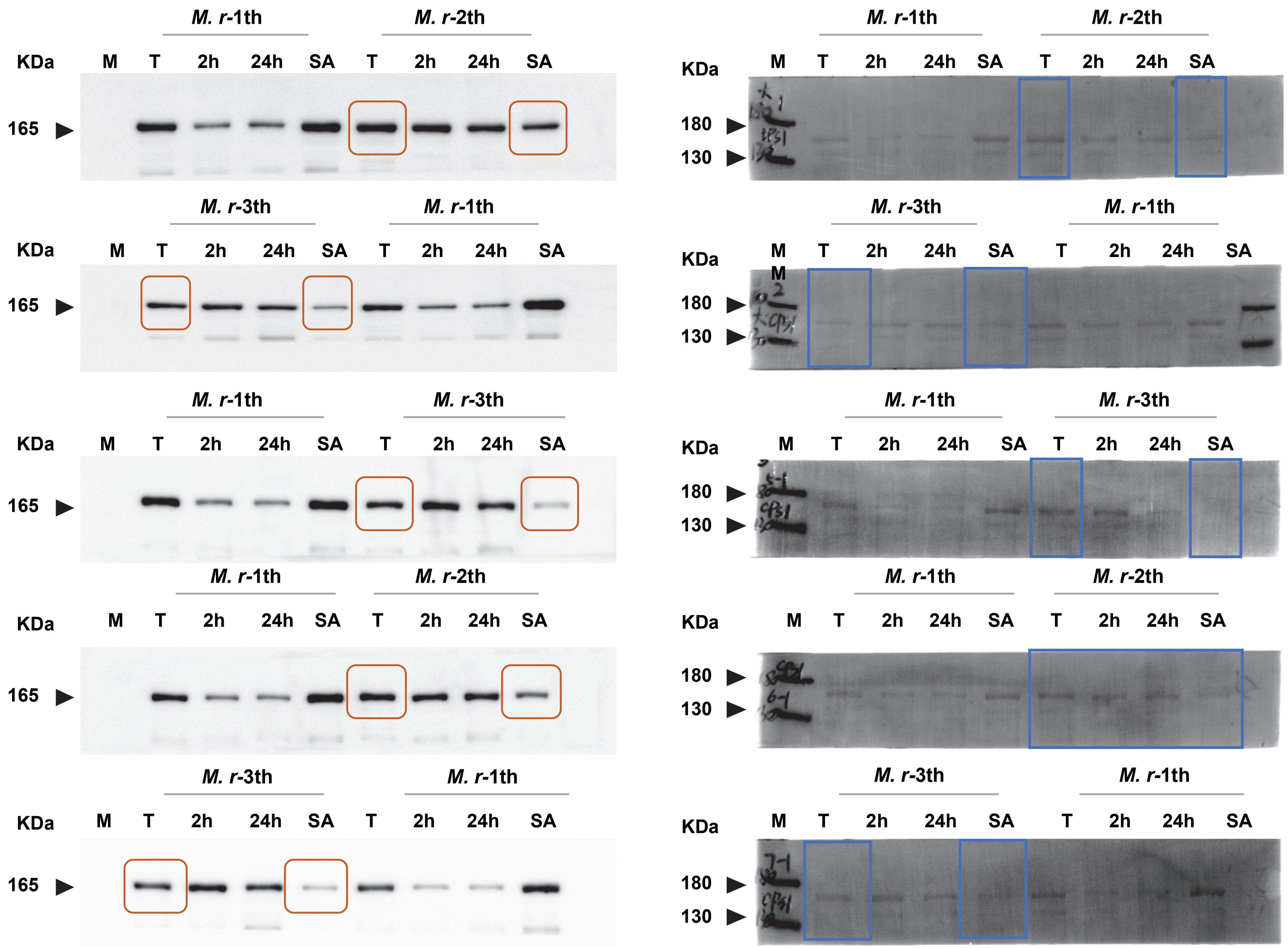
M. r-1th
M. r-2th
KDa
M T 2h 24h SA T 2h 24h SA
165
M. r-3th
M. r-1th
KDa
M T 2h 24h SA T 2h 24h SA
165
M. r-1th
M. r-3th
KDa
M T 2h 24h SA T 2h 24h SA
165
M. r-1th
M. r-2th
KDa
M T 2h 24h SA T 2h 24h SA
165
M. r-3th
M. r-1th
KDa
M T 2h 24h SA T 2h 24h SA
165
M. r-1th
M. r-2th
KDa
M T 2h 24h SA T 2h 24h SA
180
130
M. r-3th
M. r-1th
KDa
M T 2h 24h SA T 2h 24h SA
180
130
M. r-1th
M. r-3th
KDa
M T 2h 24h SA T 2h 24h SA
180
130
M. r-1th
M. r-2th
KDa
M T 2h 24h SA T 2h 24h SA
180
130
M. r-3th
M. r-1th
KDa
M T 2h 24h SA T 2h 24h SA
180
130

#### CPS1

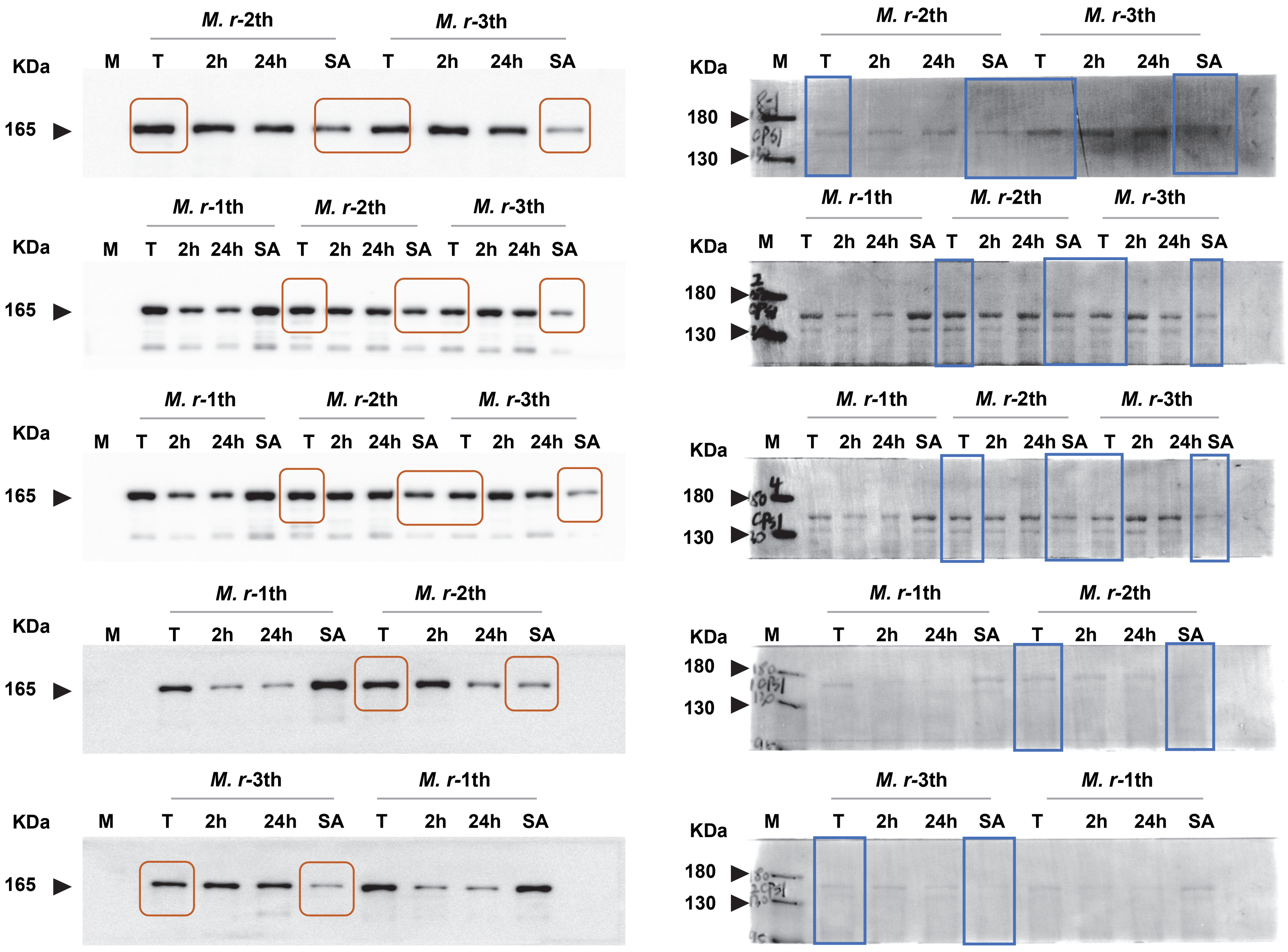

CPS1

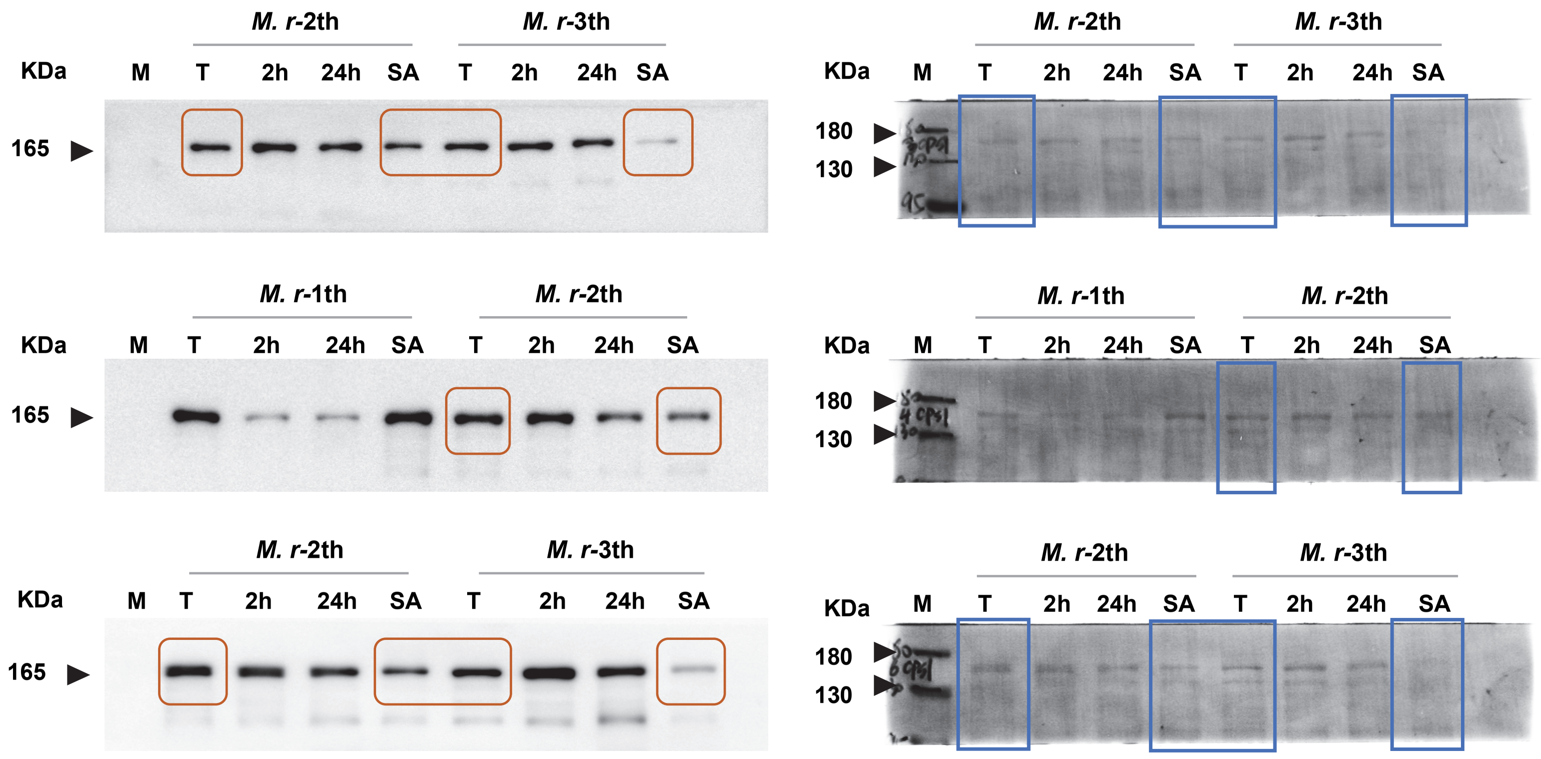

### OTC

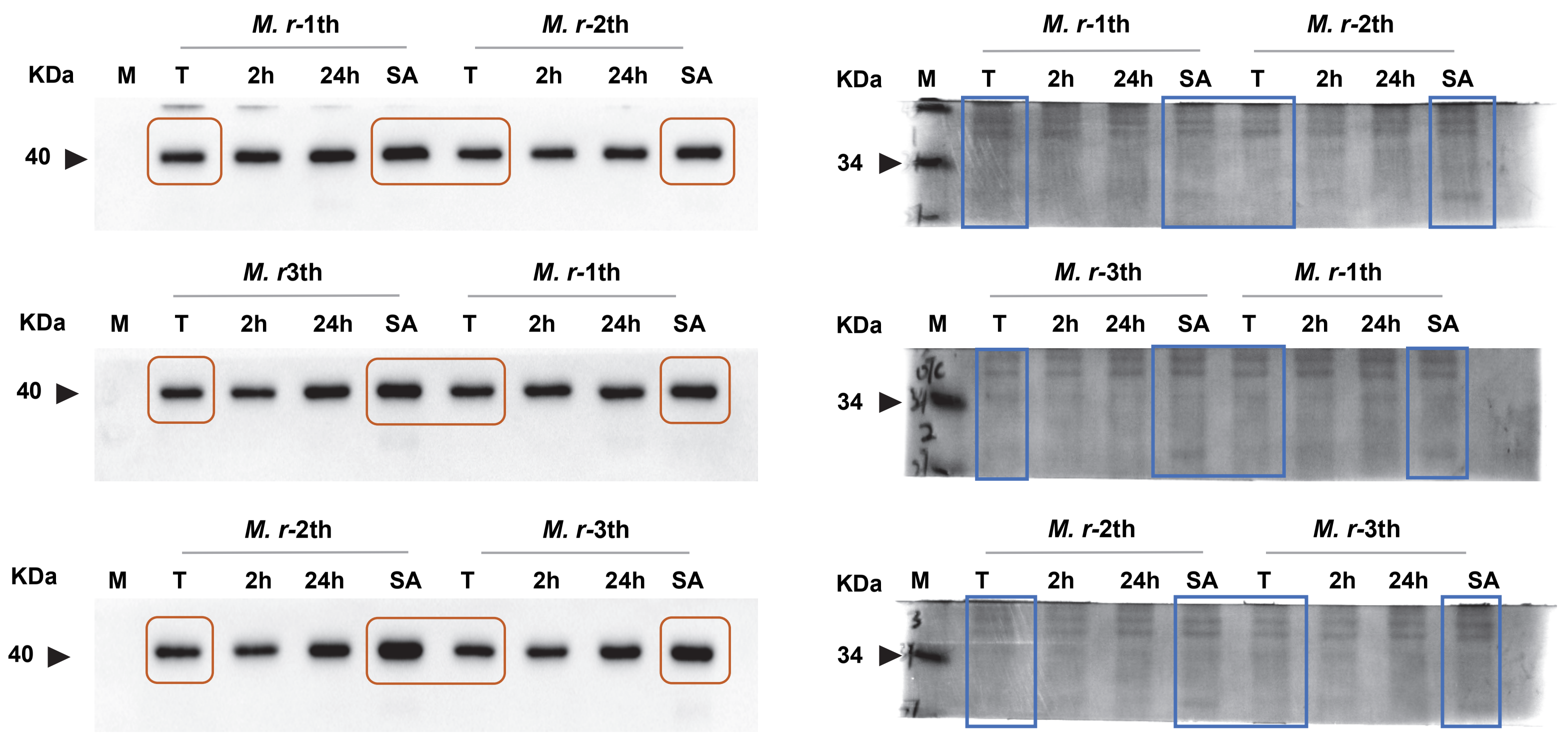
M. r-1th
M. r-2th
KDa
M
T
2h
24h
SA
T
2h
24h
SA
40
M. r3th
M. r-1th
KDa
M
T
2h
24h
SA
T
2h
24h
SA
40
M. r-2th
M. r-3th
KDa
M
T
2h
24h
SA
T
2h
24h
SA
40
M. r-1th
M. r-2th
KDa
M
T
2h
24h
SA
T
2h
24h
SA
34
M. r-3th
M. r-1th
KDa
M
T
2h
24h
SA
T
2h
24h
SA
34
M. r-2th
M. r-3th
KDa
M
T
2h
24h
SA
T
2h
24h
SA
34

OTC

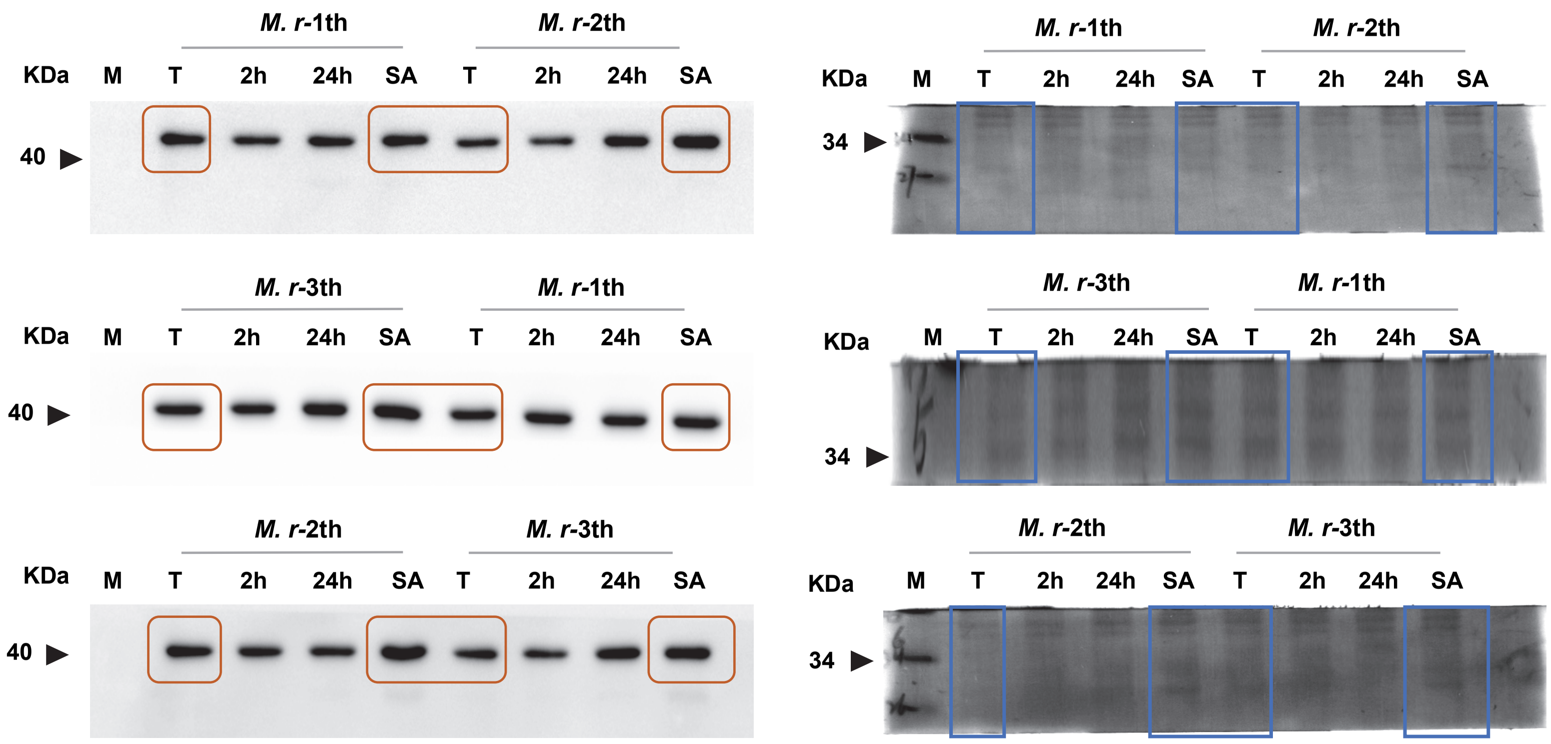

ASS1

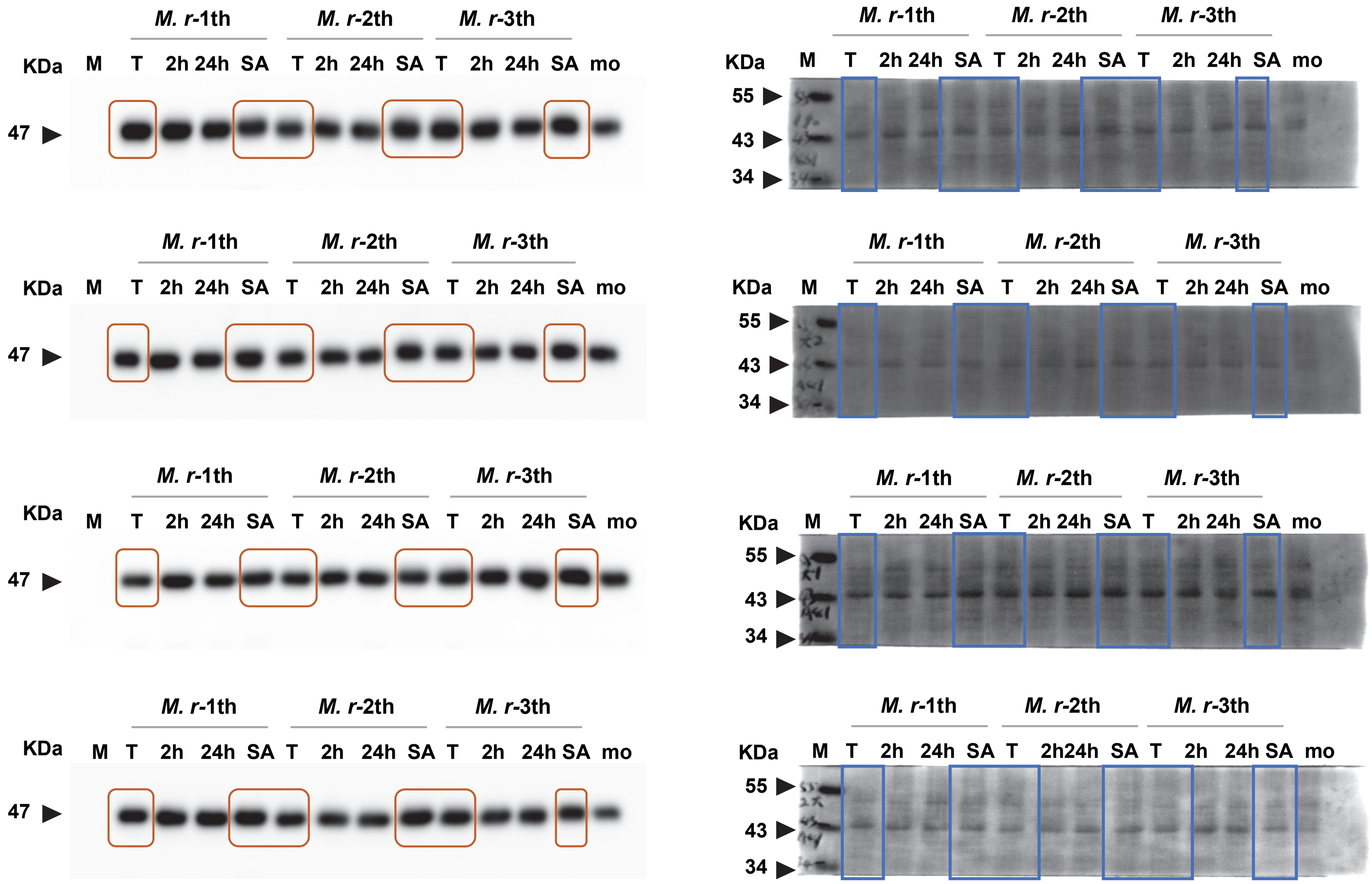

### ASS1

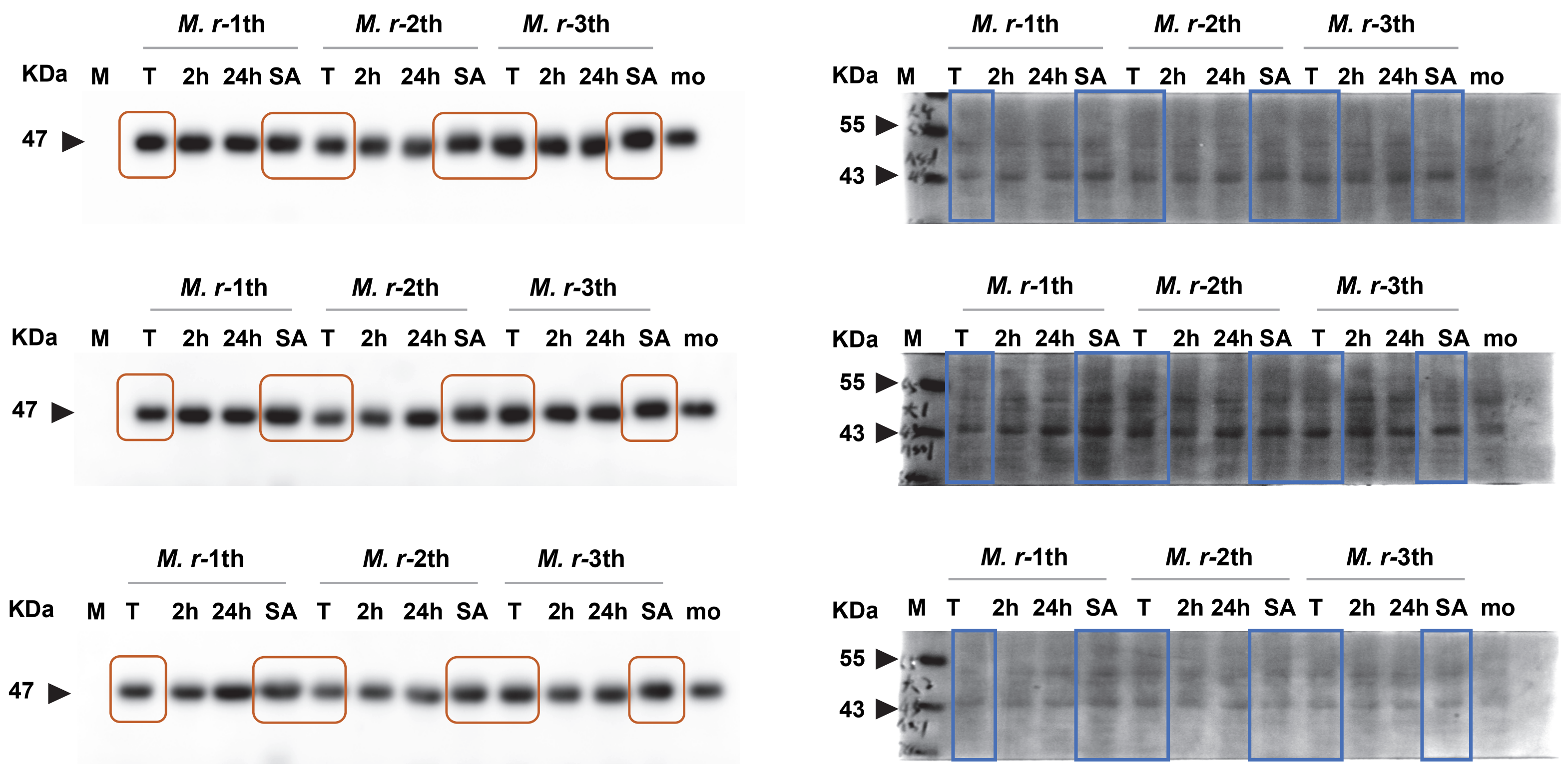
M. r-1th
M. r-2th
M. r-3th
KDa M T 2h 24h SA T 2h 24h SA T 2h 24h SA mo
47
55
43

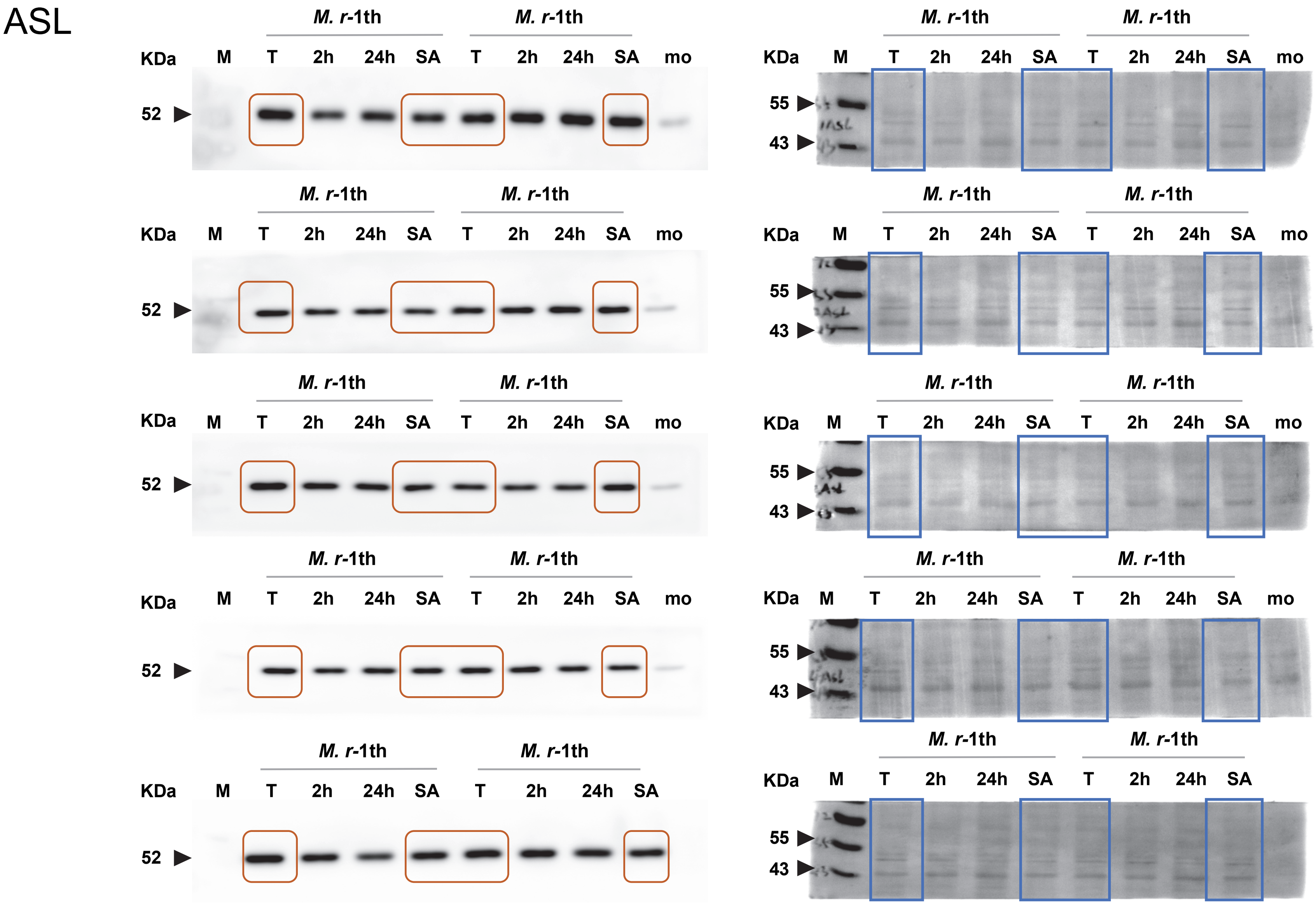
ASL
M. r-1th
KDa
M
T
2h
24h
SA
mo
52
55
43

ASL

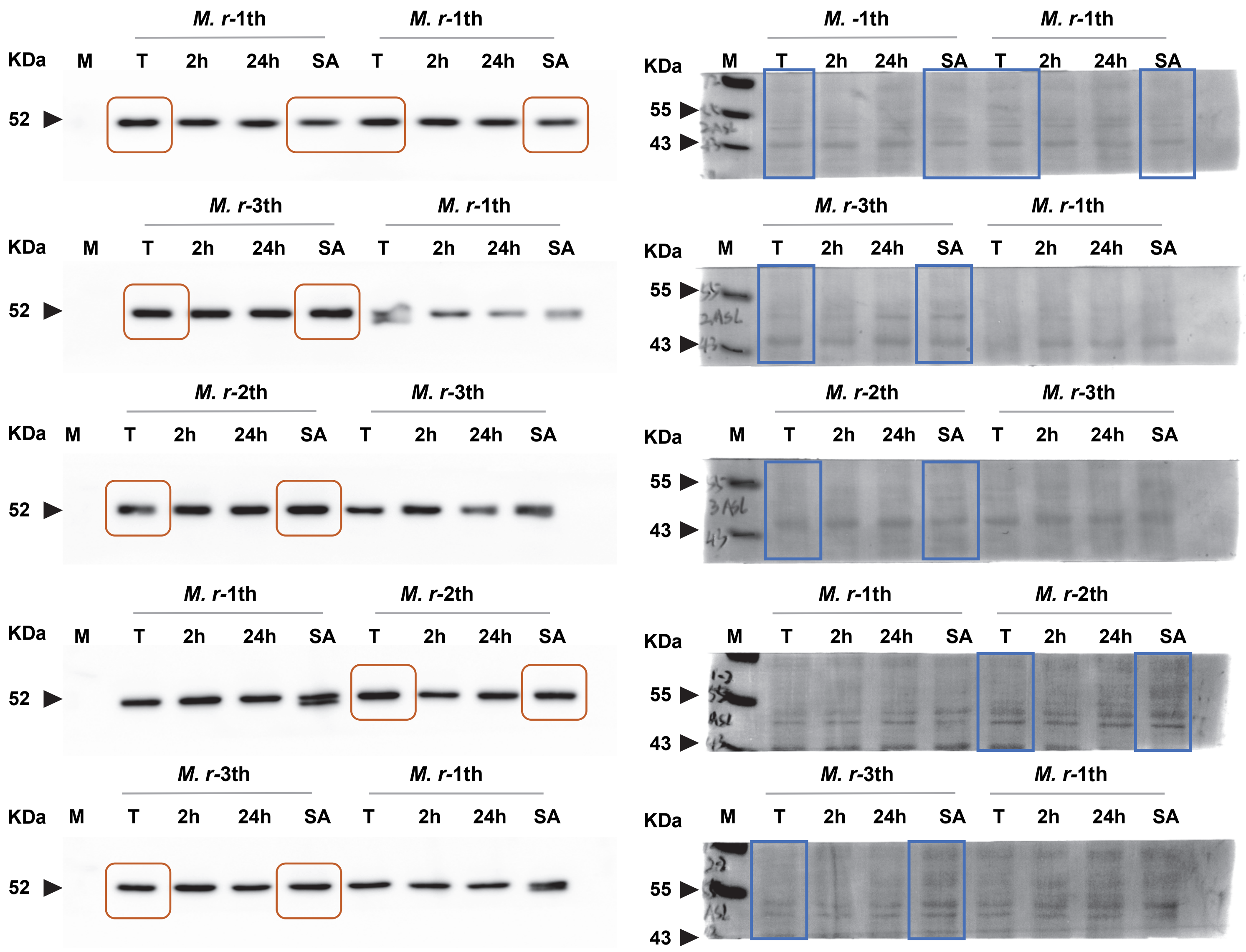

ASL

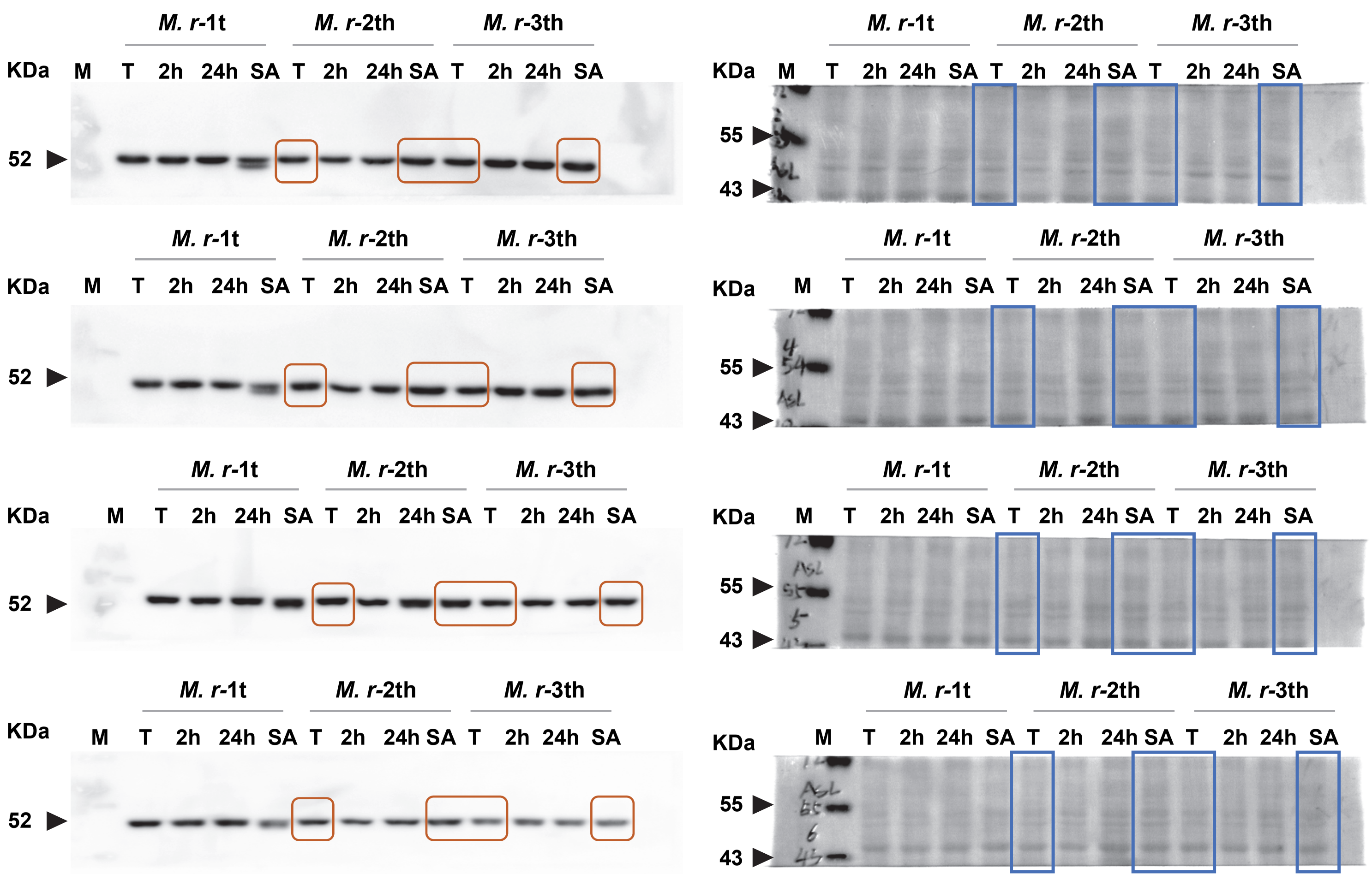

ARG

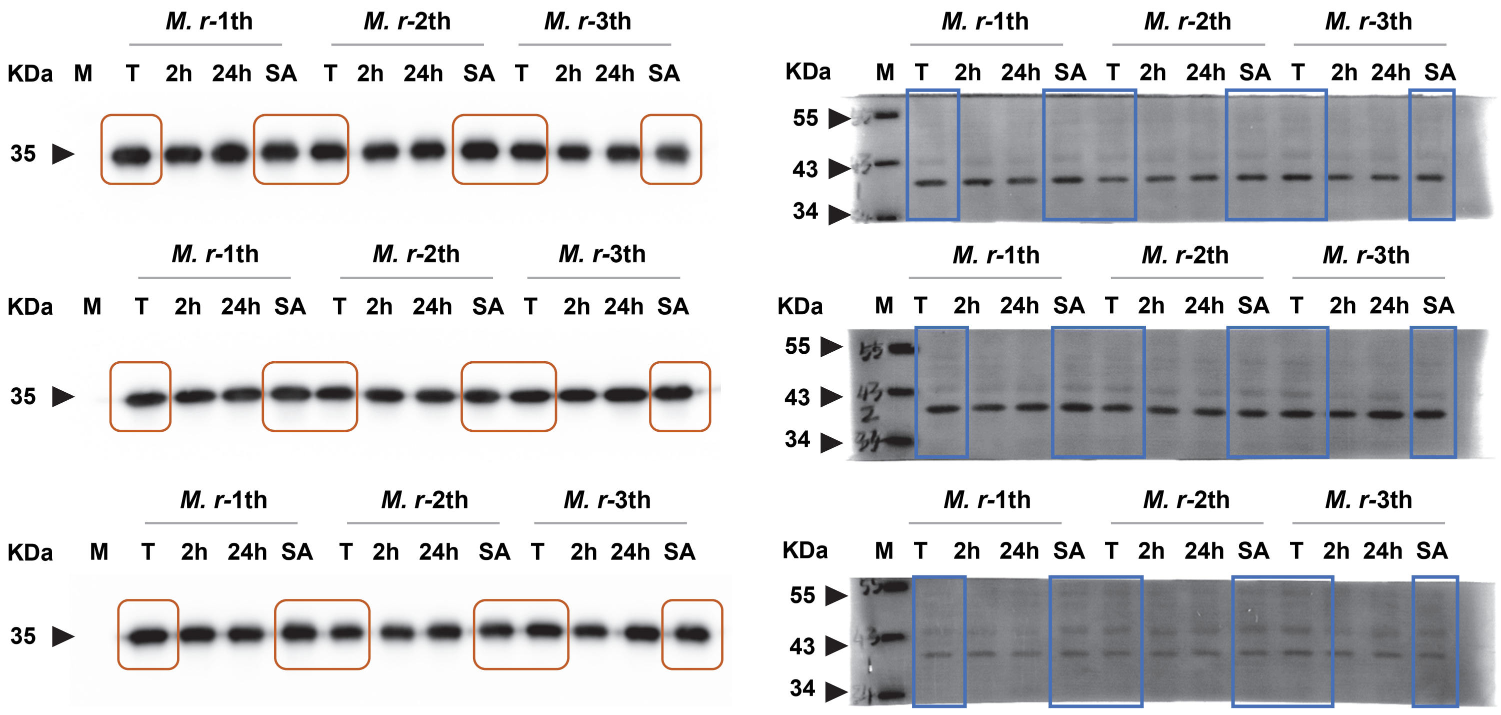

ARG

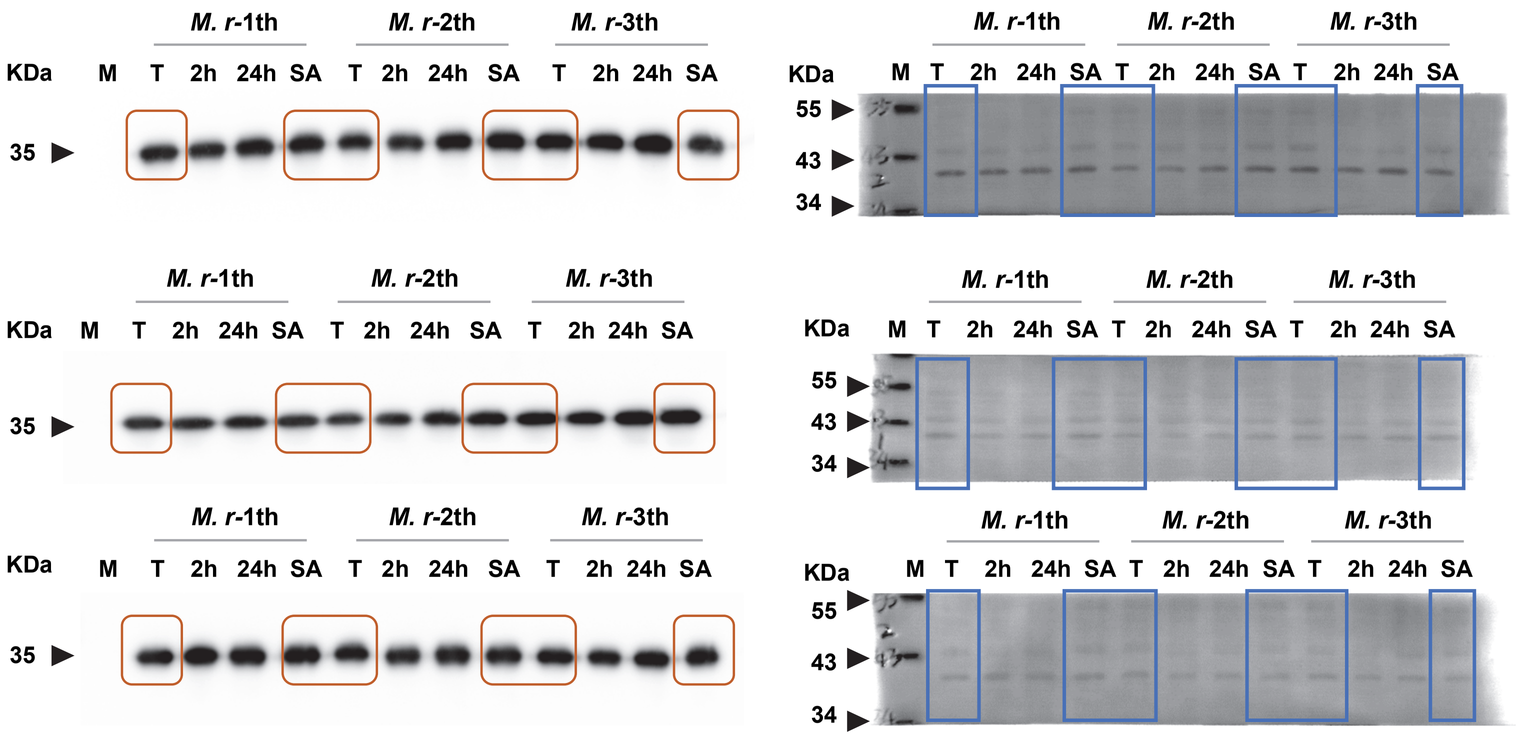

### ODC

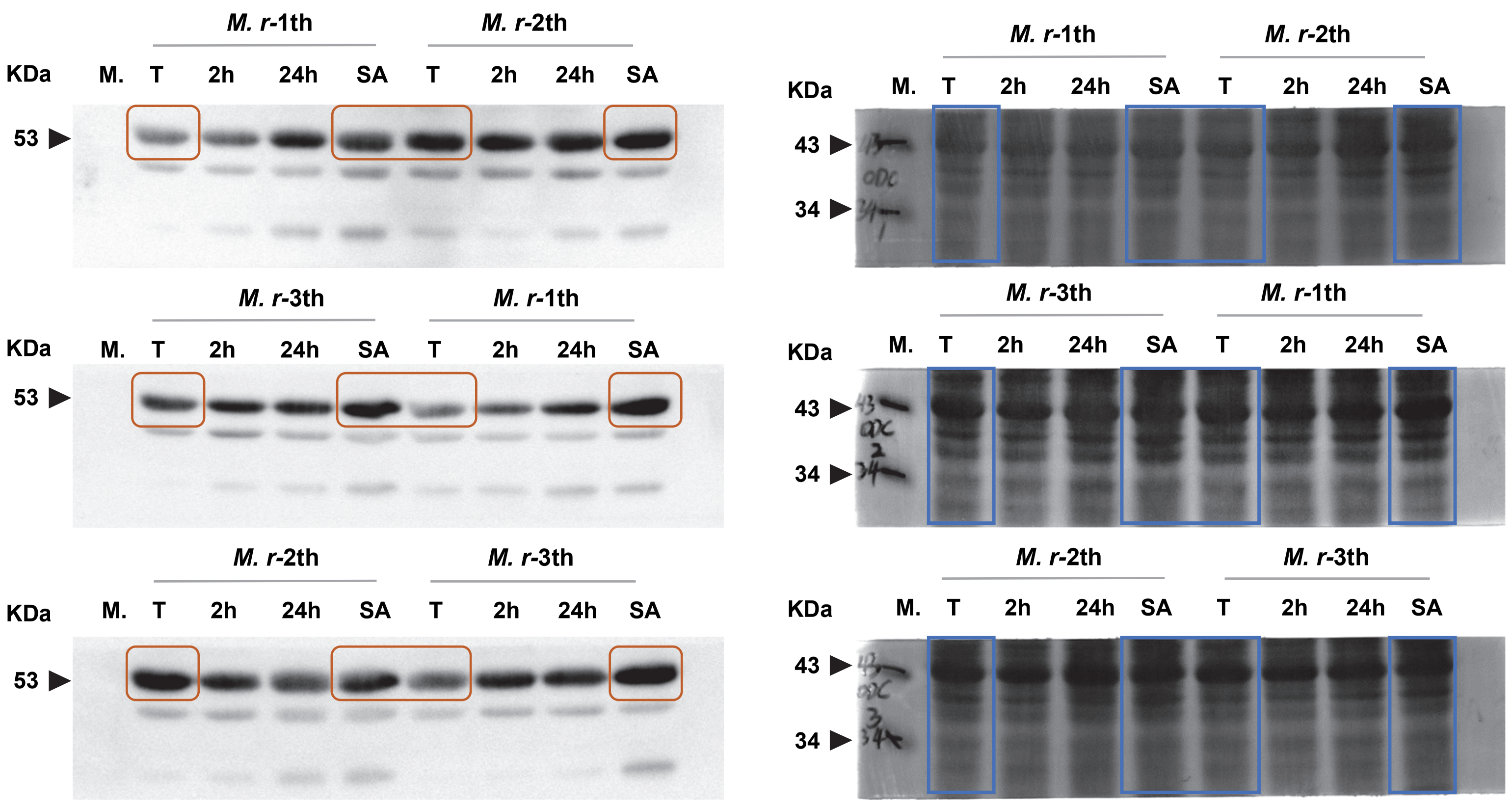

M. r-1th
M. r-2th
KDa
M.
T
2h
24h
SA
T
2h
24h
SA
53
M. r-3th
M. r-1th
53
M. r-2th
M. r-3th
53
43
34

ODC

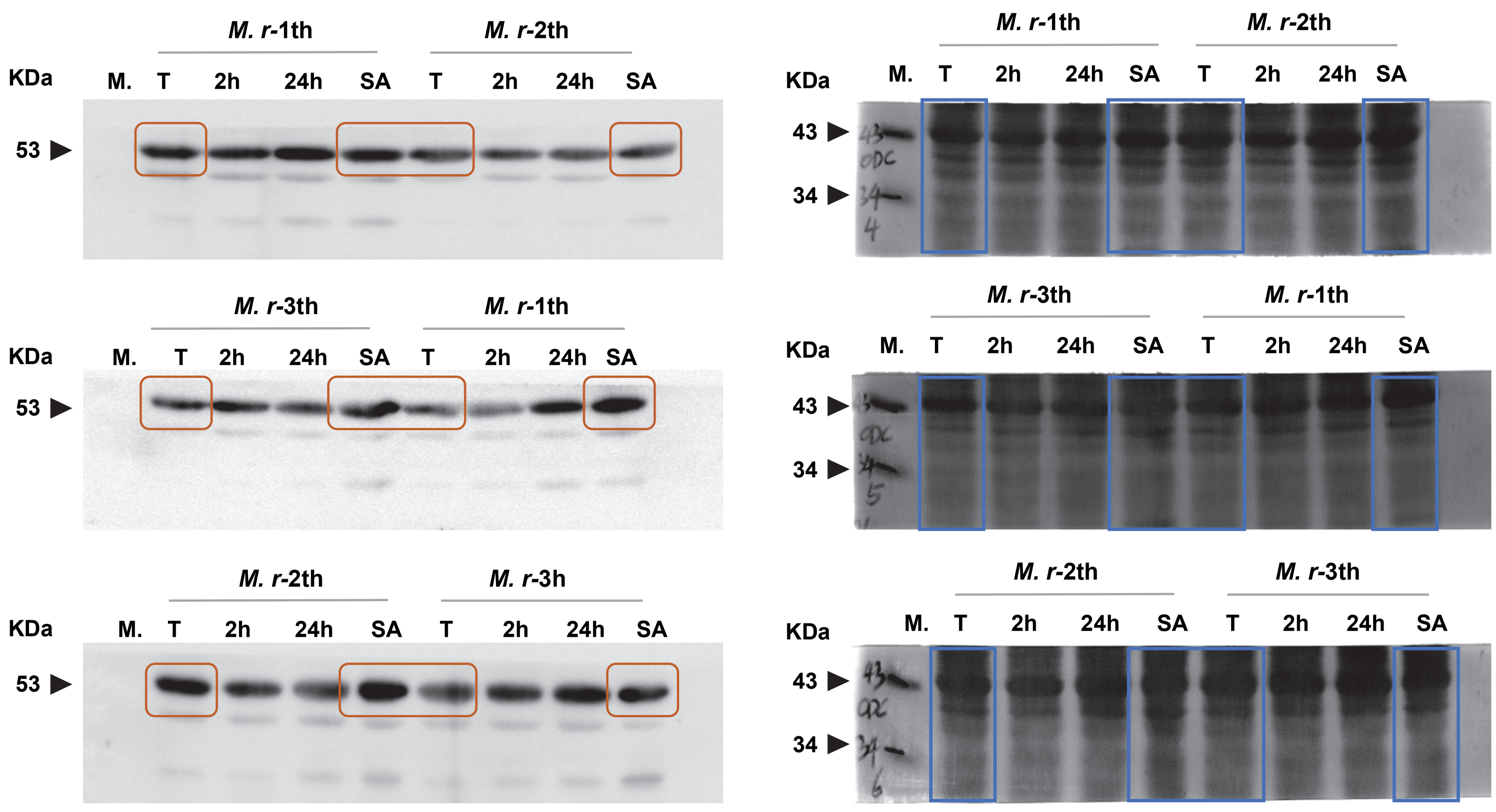
M. r-1th
M. r-2th
KDa
M.
T
2h
24h
SA
53
M. r-3th
M. r-1th
M. r-2th
M. r-3h
M. r-3th
43
34
ODC

### AGMAT

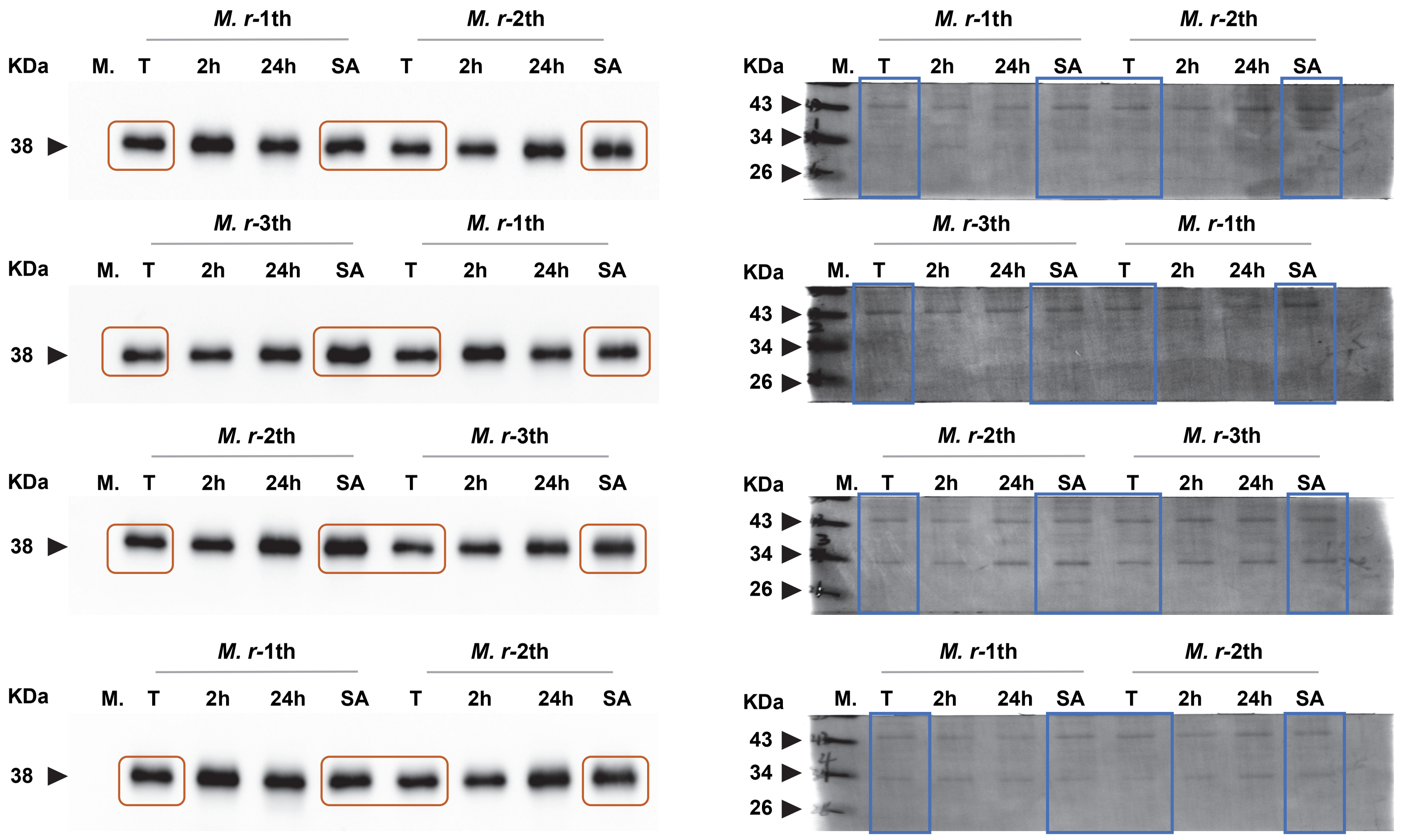

### AGMAT

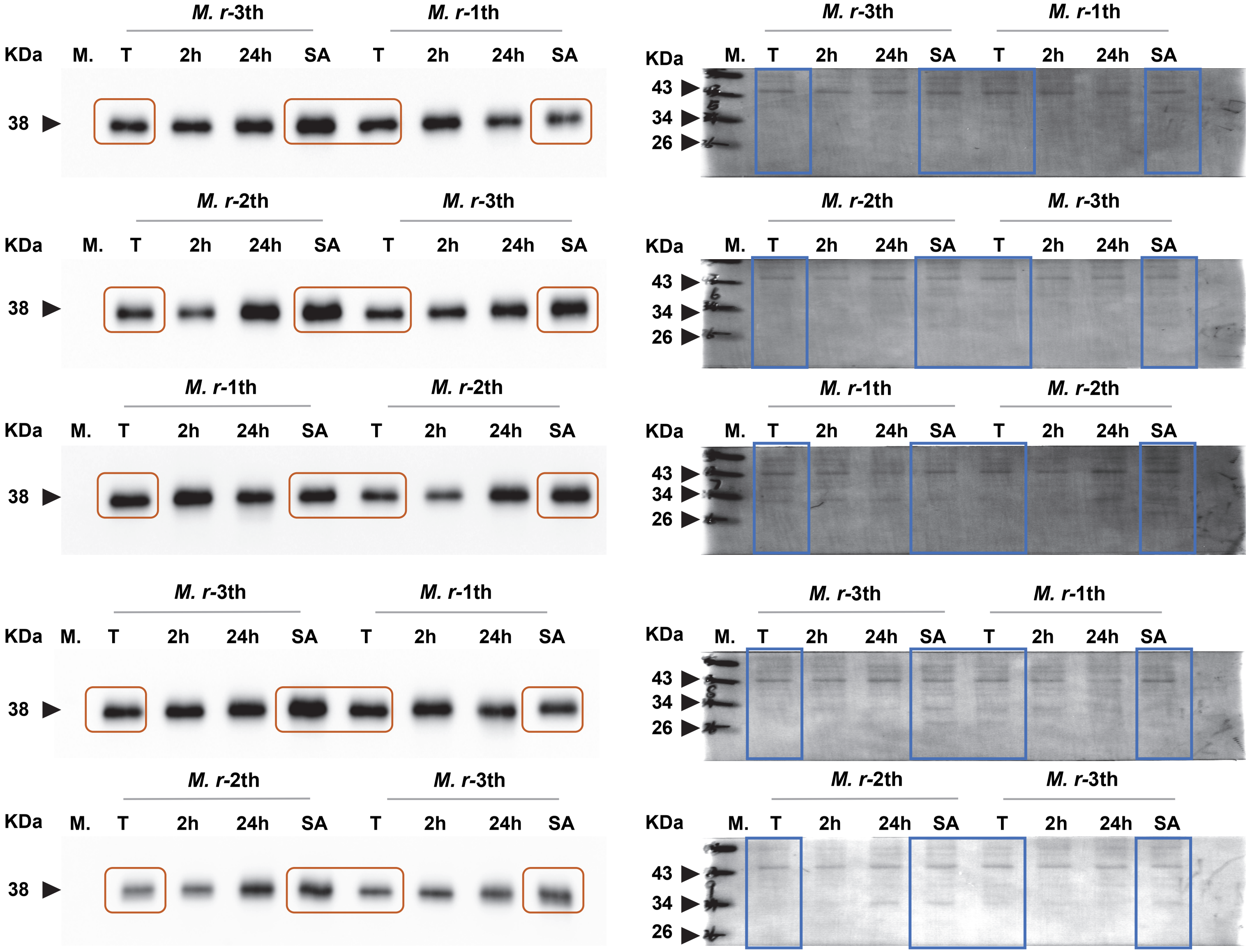

NAGS

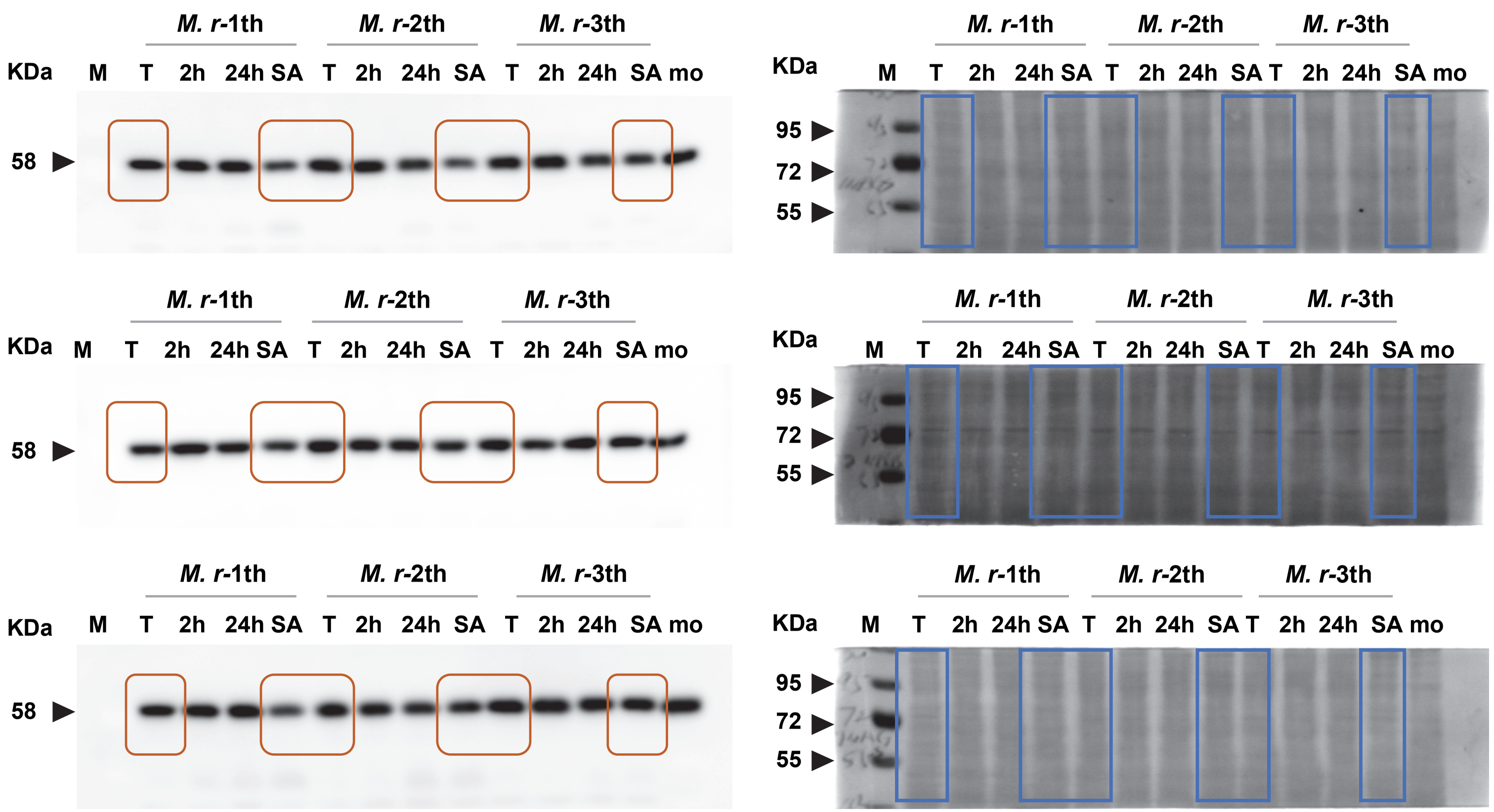
M. r-1th
M. r-2th
M. r-3th
KDa
M T 2h 24h SA T 2h 24h SA T 2h 24h SA mo
58
M. r-1th
M. r-2th
M. r-3th
KDa
M T 2h 24h SA T 2h 24h SA T 2h 24h SA mo
95
72
55

NAGS

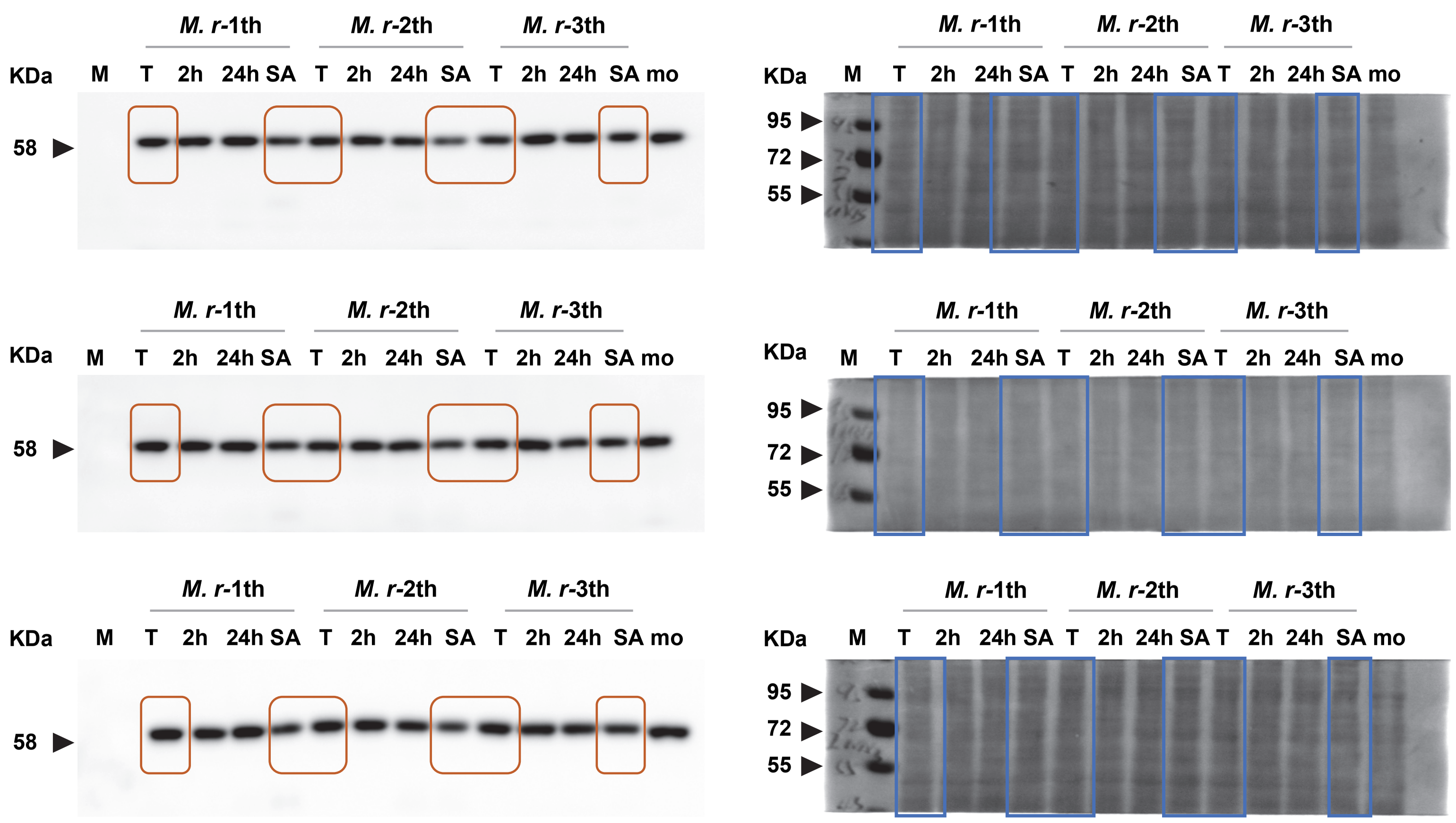
M. r-1th
M. r-2th
M. r-3th
KDa
M T 2h 24h SA T 2h 24h SA T 2h 24h SA mo
58
95
72
55

### SIRT5

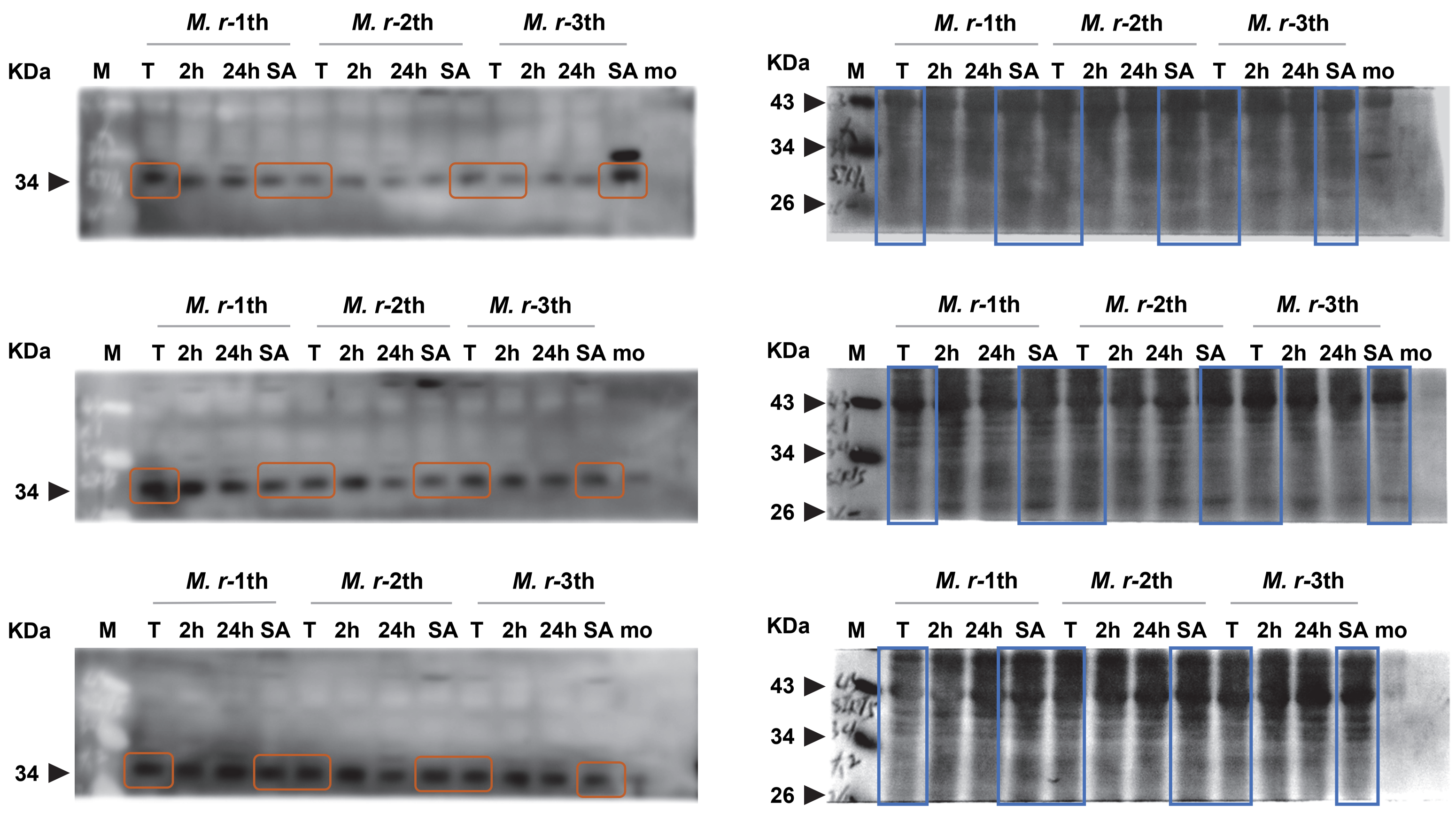
M. r-1th
M. r-2th
M. r-3th
KDa
M
T 2h 24h SA T 2h 24h SA T 2h 24h SA mo
34
43
26

SIRT5

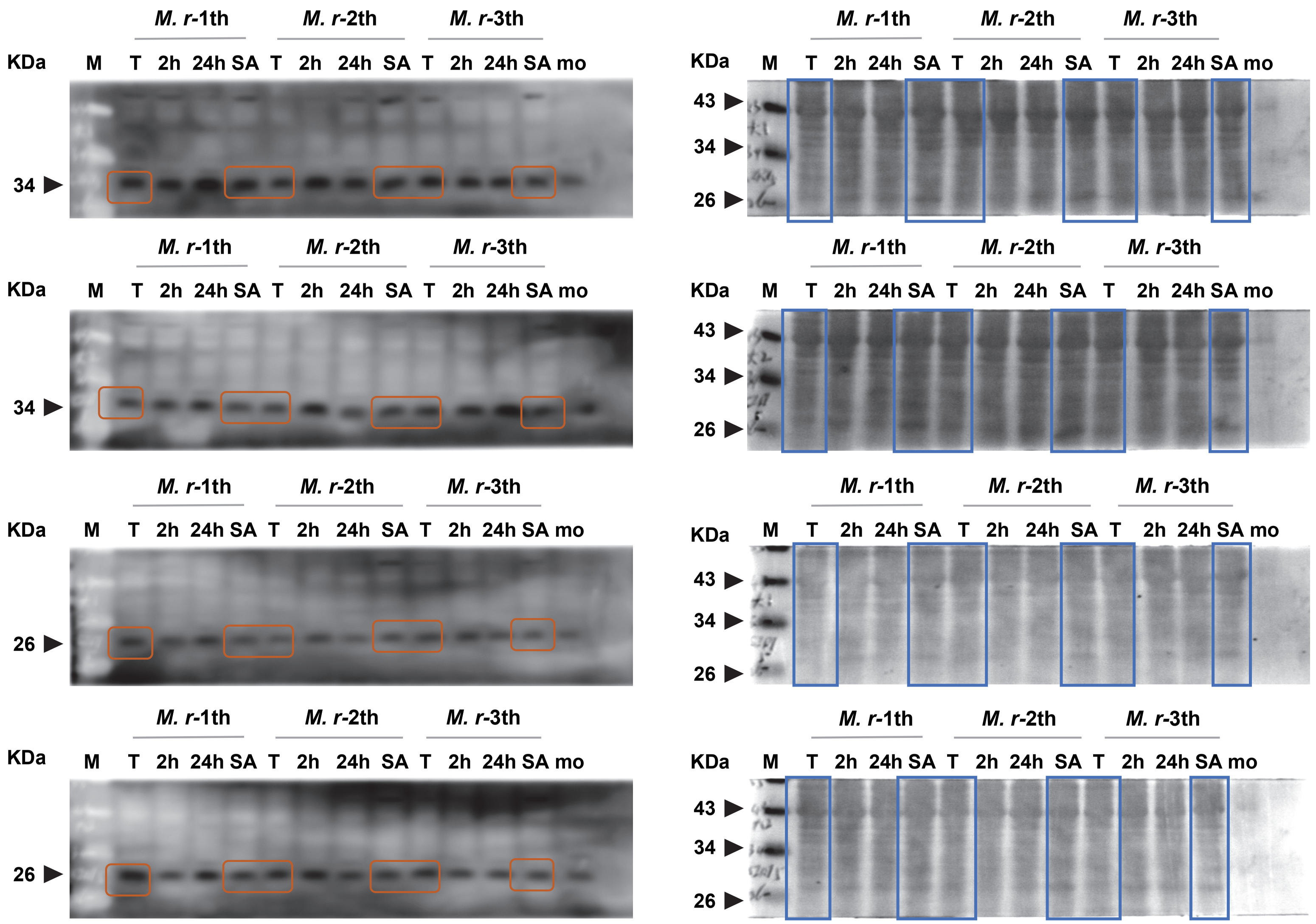
M. r-1th
M. r-2th
M. r-3th
KDa
M T 2h 24h SA T 2h 24h SA T 2h 24h SA mo
34
M. r-1th
M. r-2th
M. r-3th
KDa
M T 2h 24h SA T 2h 24h SA T 2h 24h SA mo
43
34
26
M. r-1th
M. r-2th
M. r-3th
KDa
M T 2h 24h SA T 2h 24h SA T 2h 24h SA mo
34
M. r-1th
M. r-2th
M. r-3th
KDa
M T 2h 24h SA T 2h 24h SA T 2h 24h SA mo
43
34
26
M. r-1th
M. r-2th
M. r-3th
KDa
M T 2h 24h SA T 2h 24h SA T 2h 24h SA mo
26
M. r-1th
M. r-2th
M. r-3th
KDa
M T 2h 24h SA T 2h 24h SA T 2h 24h SA mo
43
34
26
M. r-1th
M. r-2th
M. r-3th
KDa
M T 2h 24h SA T 2h 24h SA T 2h 24h SA mo
26
M. r-1th
M. r-2th
M. r-3th
KDa
M T 2h 24h SA T 2h 24h SA T 2h 24h SA mo
43
34
26

### CPS1

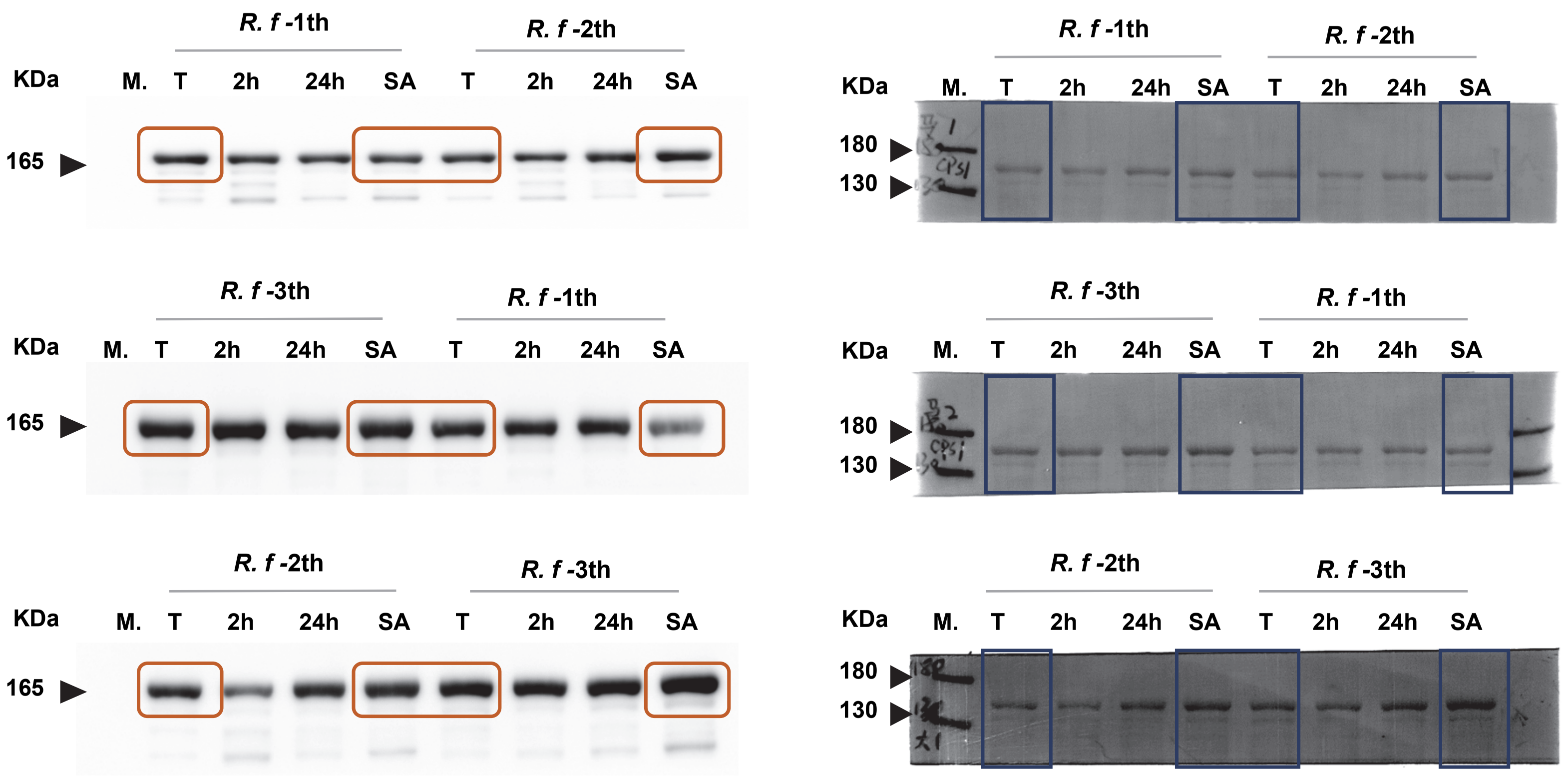
R. f -1th
R. f -2th
KDa
M. T 2h 24h SA T 2h 24h SA
165
R. f -3th
R. f -1th
KDa
M. T 2h 24h SA T 2h 24h SA
165
R. f -2th
R. f -3th
KDa
M. T 2h 24h SA T 2h 24h SA
165
R. f -1th
R. f -2th
KDa
M. T 2h 24h SA T 2h 24h SA
180
130
R. f -3th
R. f -1th
KDa
M. T 2h 24h SA T 2h 24h SA
180
130
R. f -2th
R. f -3th
KDa
M. T 2h 24h SA T 2h 24h SA
180
130

### CPS1

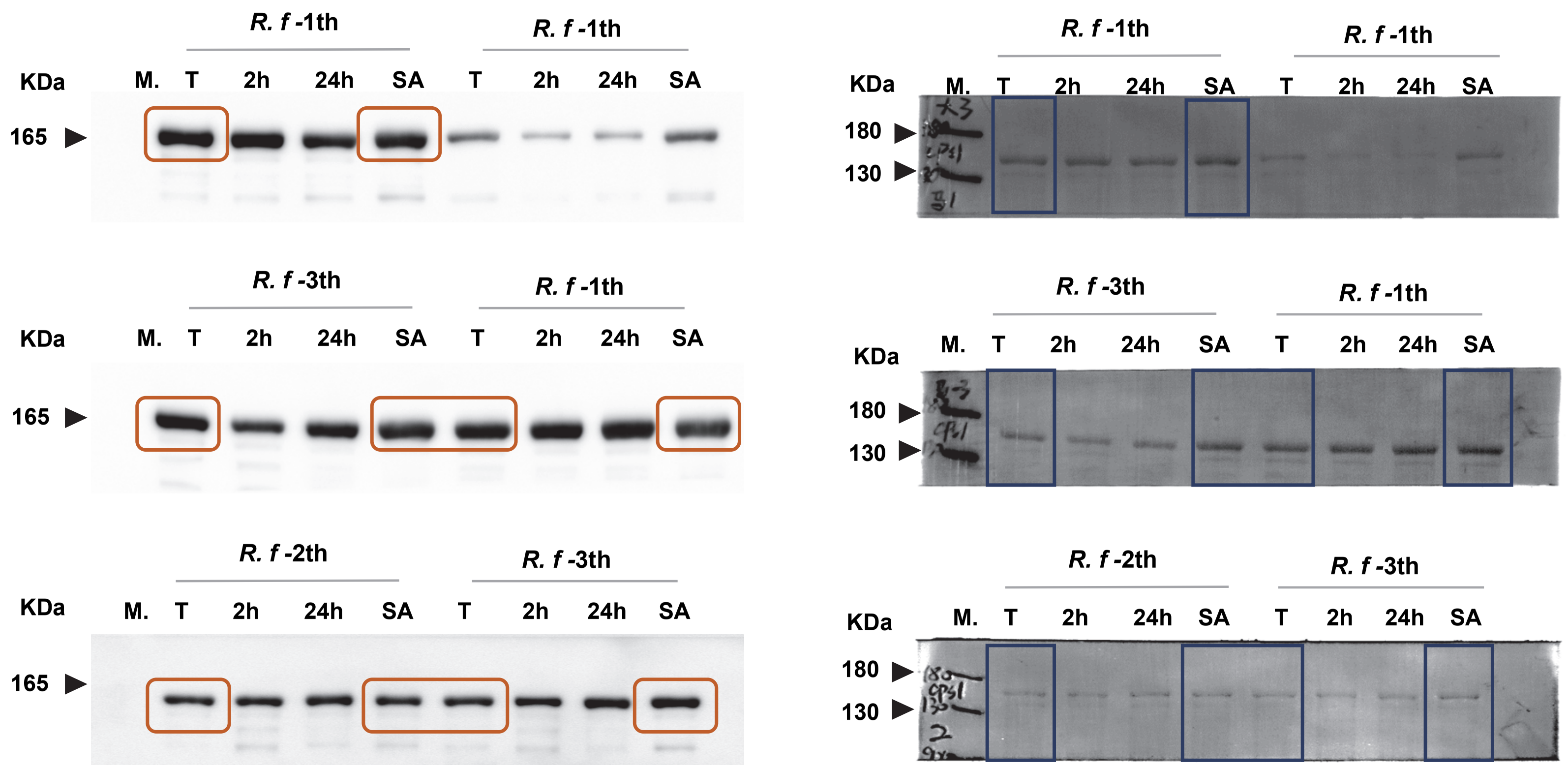
R. f -1th
R. f -1th
KDa
M. T 2h 24h SA T 2h 24h SA
165
R. f -3th
R. f -1th
KDa
M. T 2h 24h SA T 2h 24h SA
165
R. f -2th
R. f -3th
KDa
M. T 2h 24h SA T 2h 24h SA
165
R. f -1th
R. f -1th
KDa
M. T 2h 24h SA T 2h 24h SA
180
130
R. f -3th
R. f -1th
KDa
M. T 2h 24h SA T 2h 24h SA
180
130
R. f -2th
R. f -3th
KDa
M. T 2h 24h SA T 2h 24h SA
180
130

### OTC

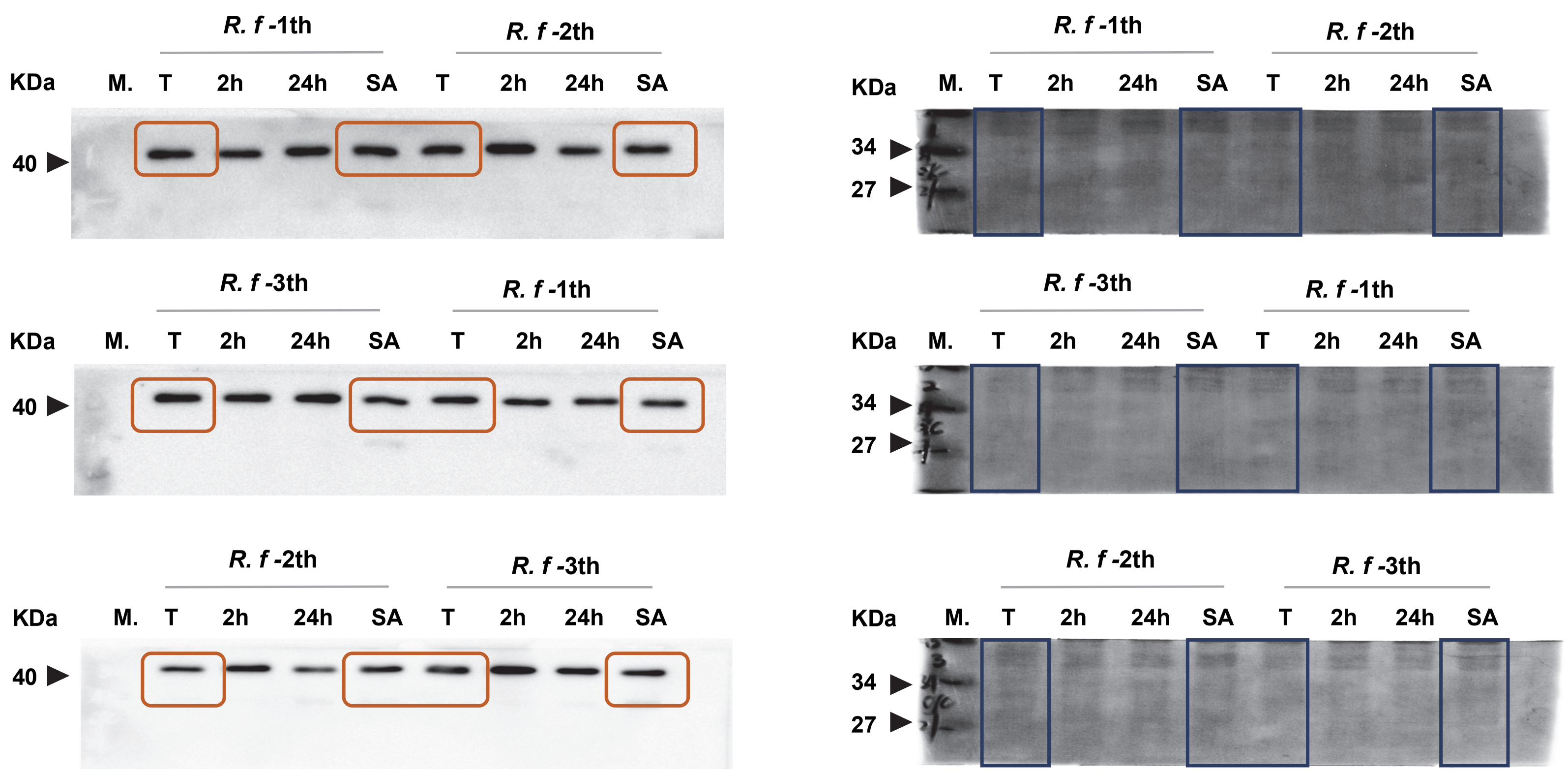

### OTC

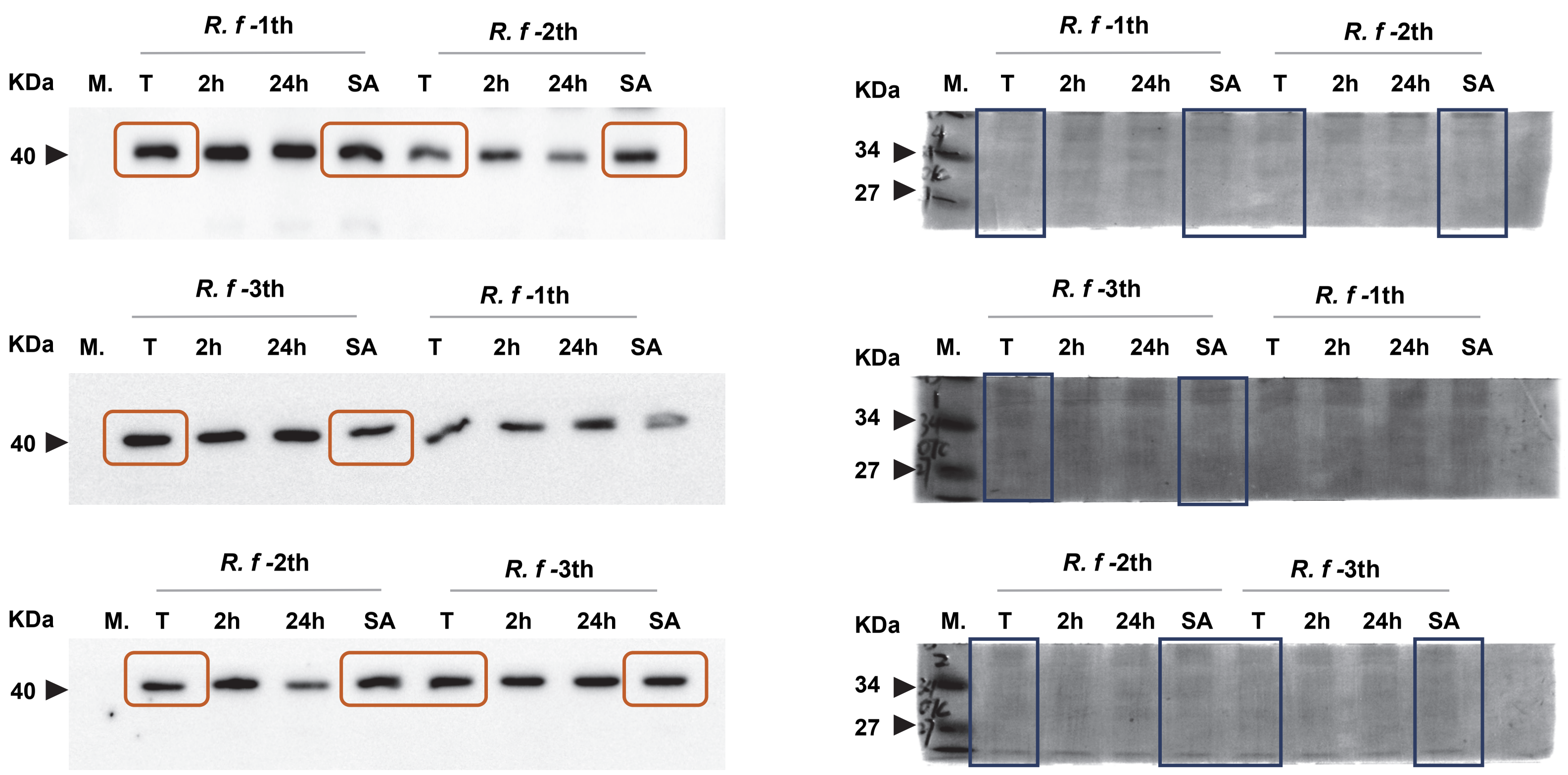

R. f -1th
R. f -2th
KDa M. T 2h 24h SA T 2h 24h SA
40
R. f -3th
R. f -1th
KDa M. T 2h 24h SA T 2h 24h SA
40
R. f -2th
R. f -3th
KDa M. T 2h 24h SA T 2h 24h SA
40
R. f -1th
R. f -2th
KDa M. T 2h 24h SA T 2h 24h SA
34
27
R. f -3th
R. f -1th
KDa M. T 2h 24h SA T 2h 24h SA
34
27
R. f -2th
R. f -3th
KDa M. T 2h 24h SA T 2h 24h SA
34
27

### ASS1

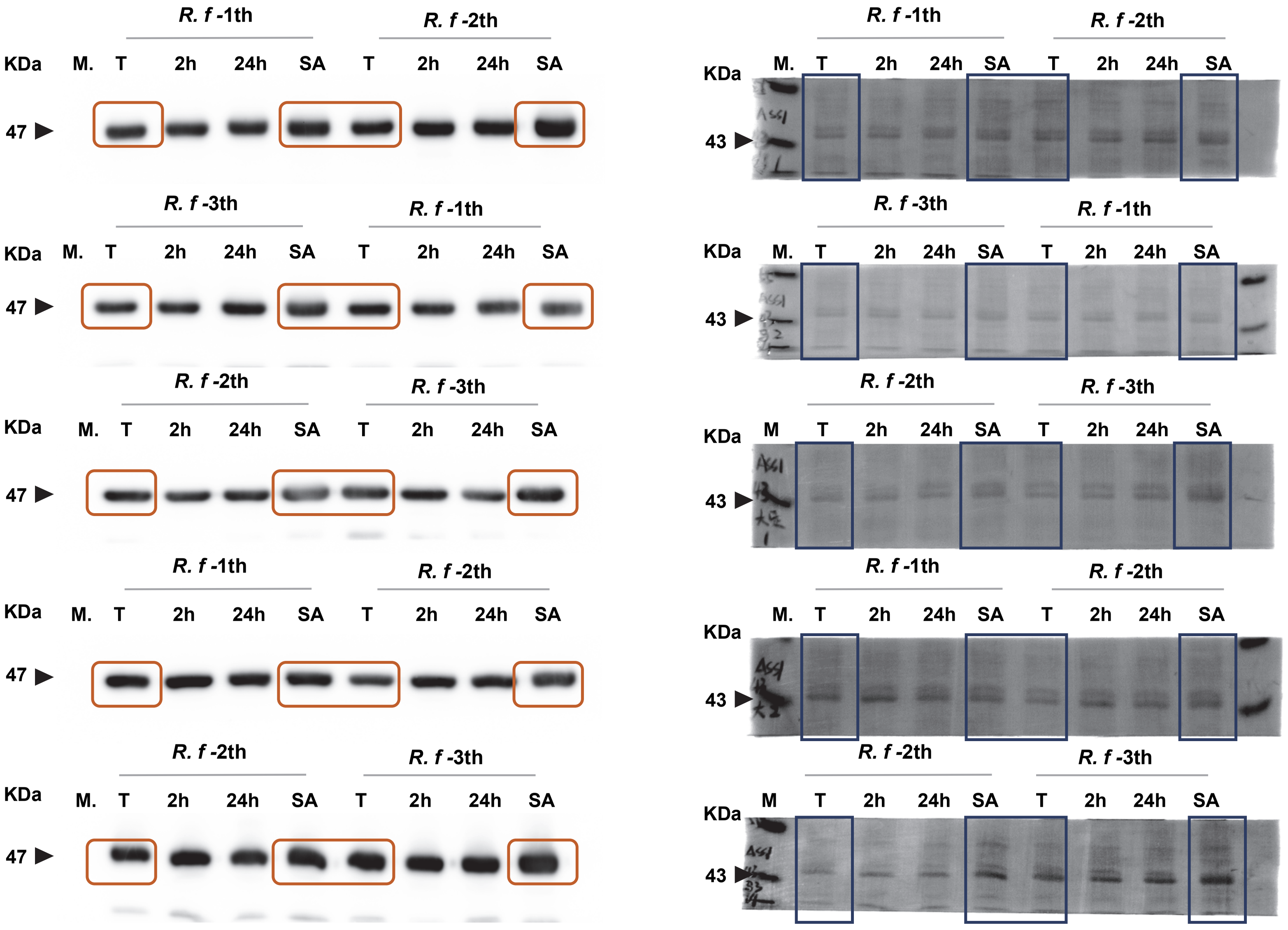

### ASL

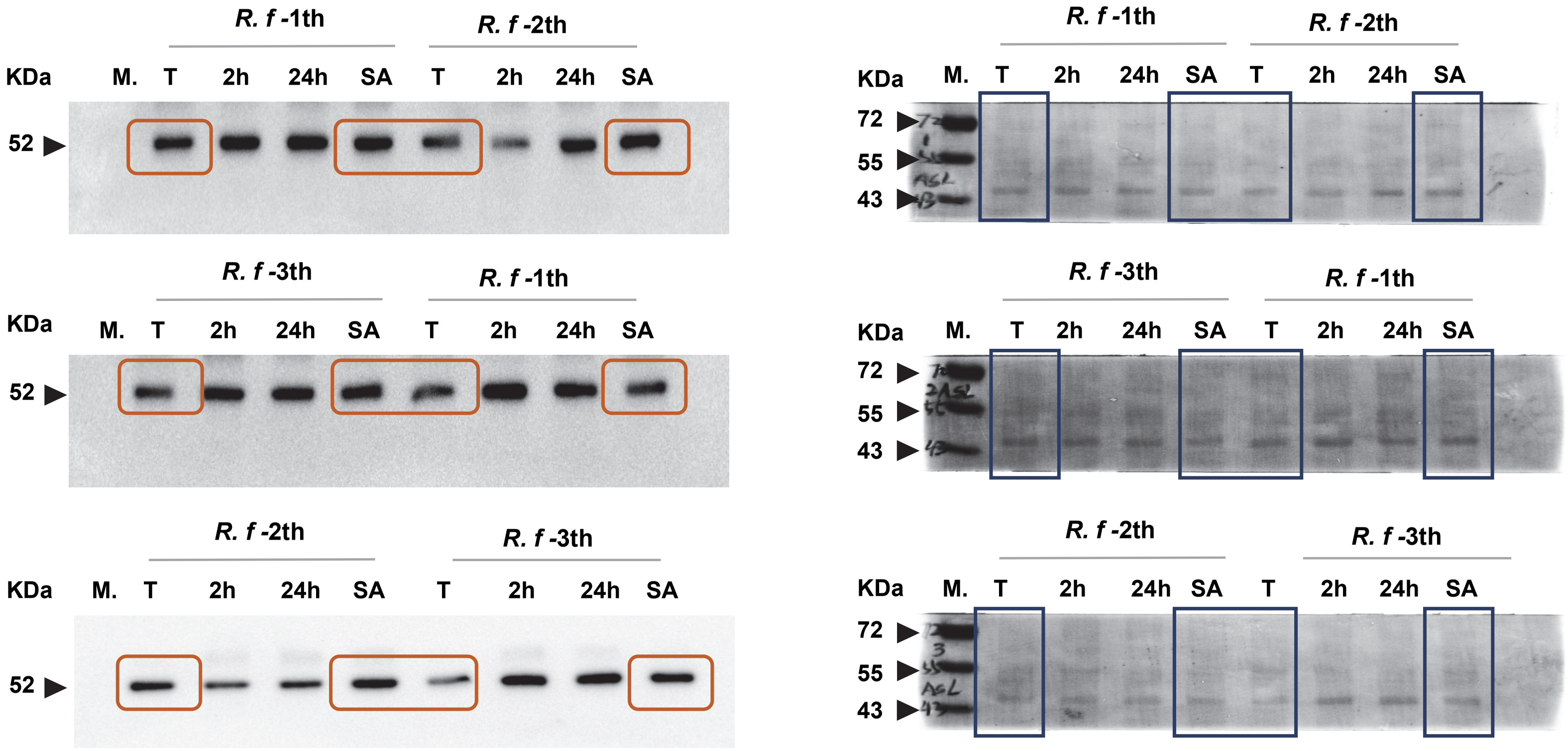

### ASL

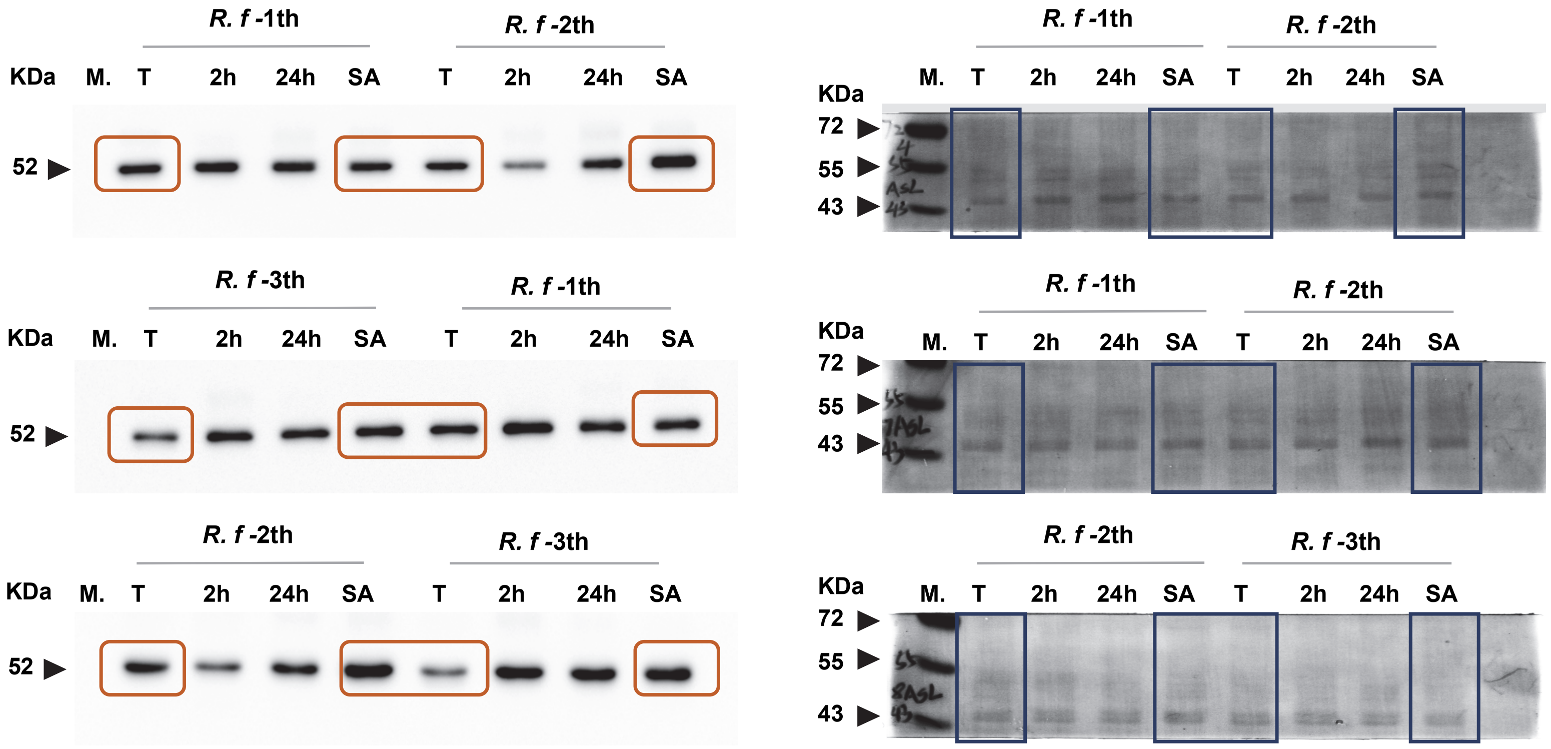

R. f -1th
R. f -2th
KDa
M.
T
2h
24h
SA
52
R. f -3th
R. f -1th
R. f -2th
R. f -3th
72
55
43

### ARG

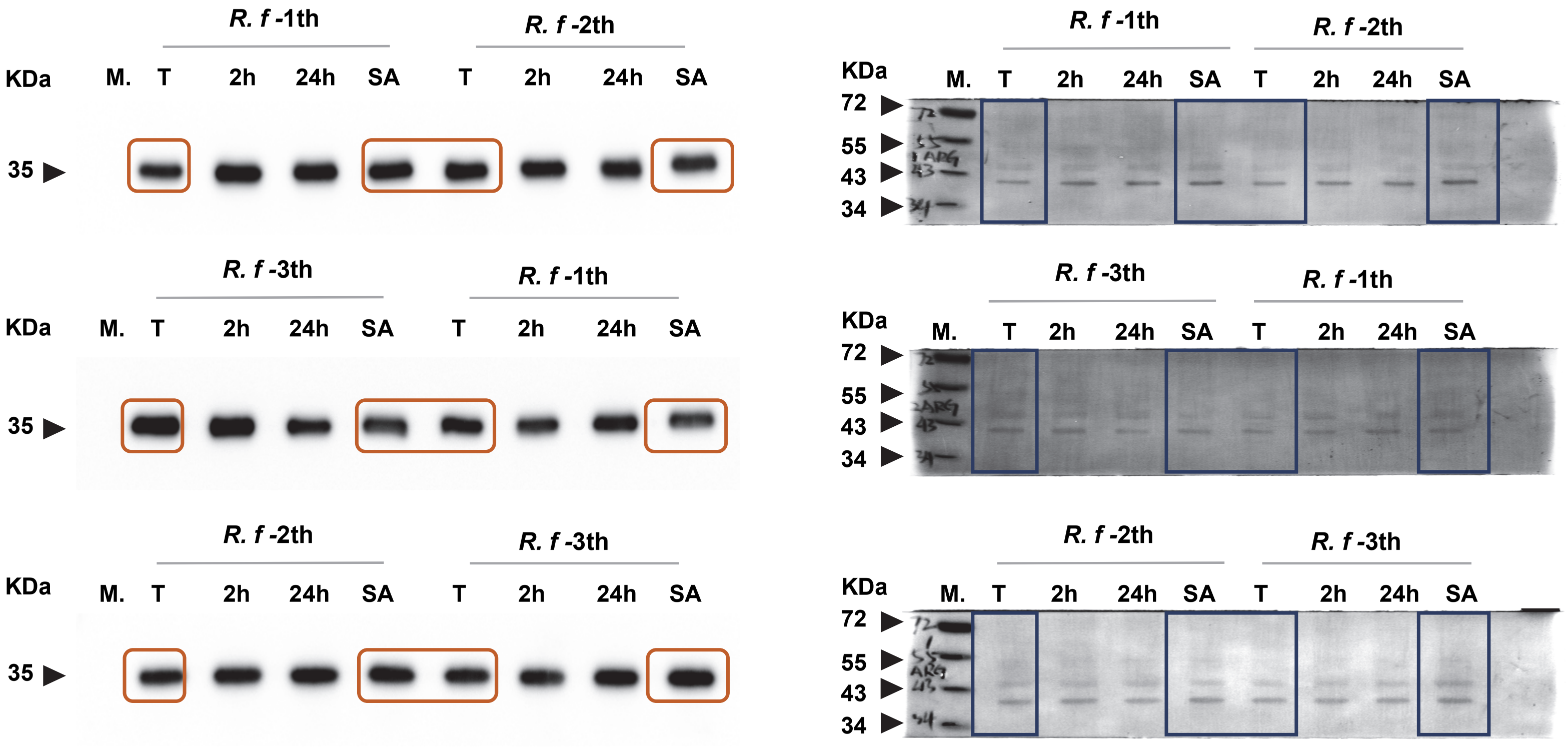
R. f -1th
R. f -2th
KDa
M.
T
2h
24h
SA
35
R. f -3th
R. f -1th
R. f -2th
R. f -3th
72
55
43
34

### ARG

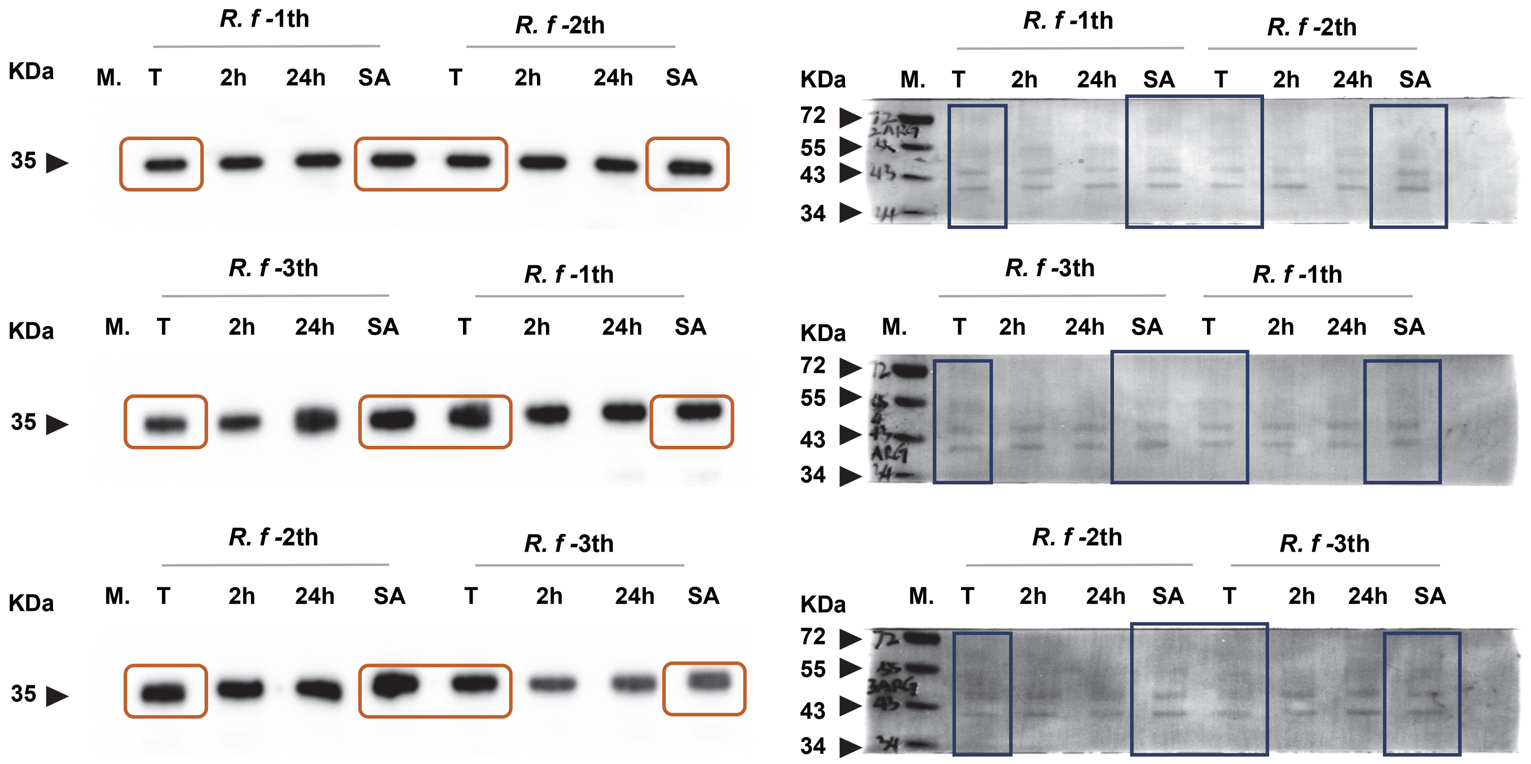

**ODC**

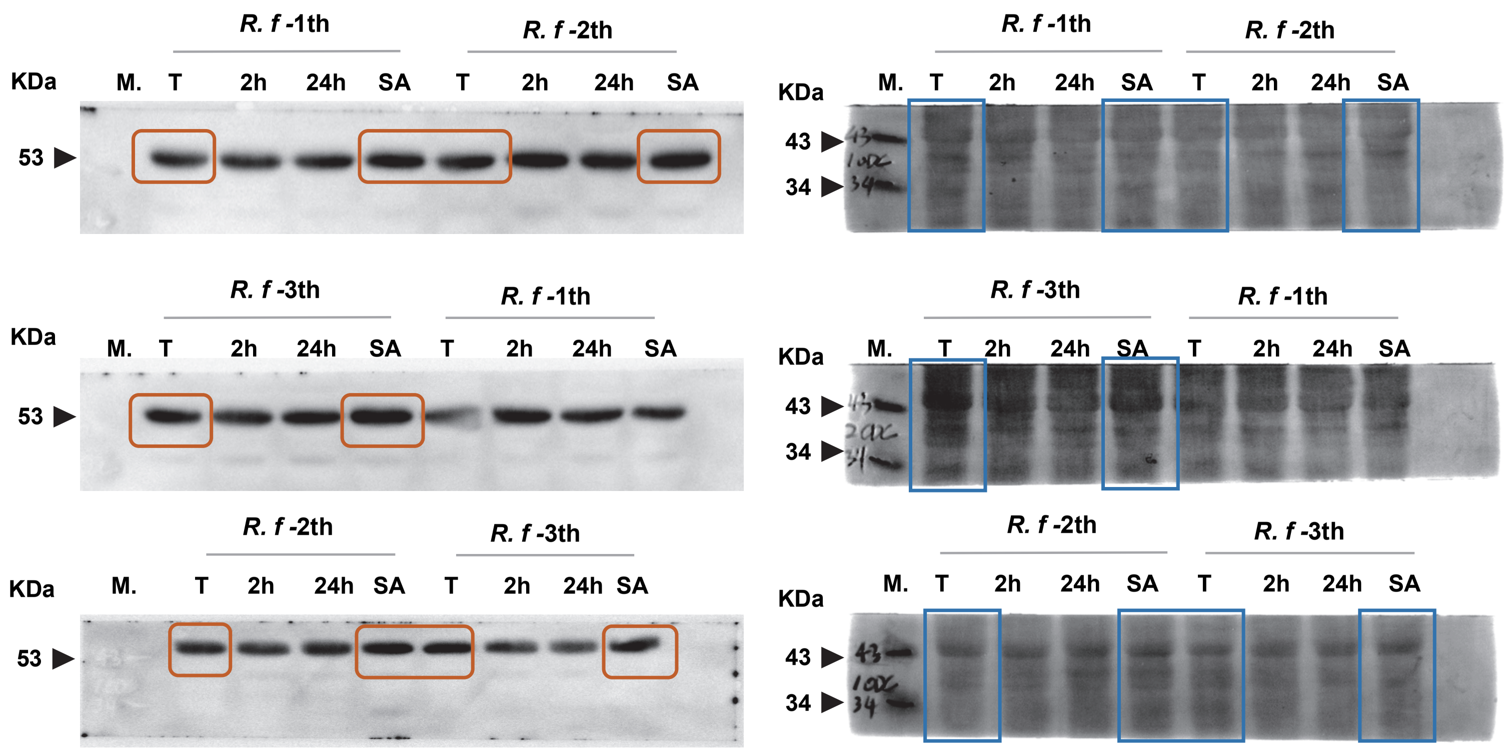

R. f -1th
R. f -2th
KDa
M. T 2h 24h SA T 2h 24h SA
53
43
34
R. f -3th
R. f -1th
R. f -2th
R. f -3th

#### ODC

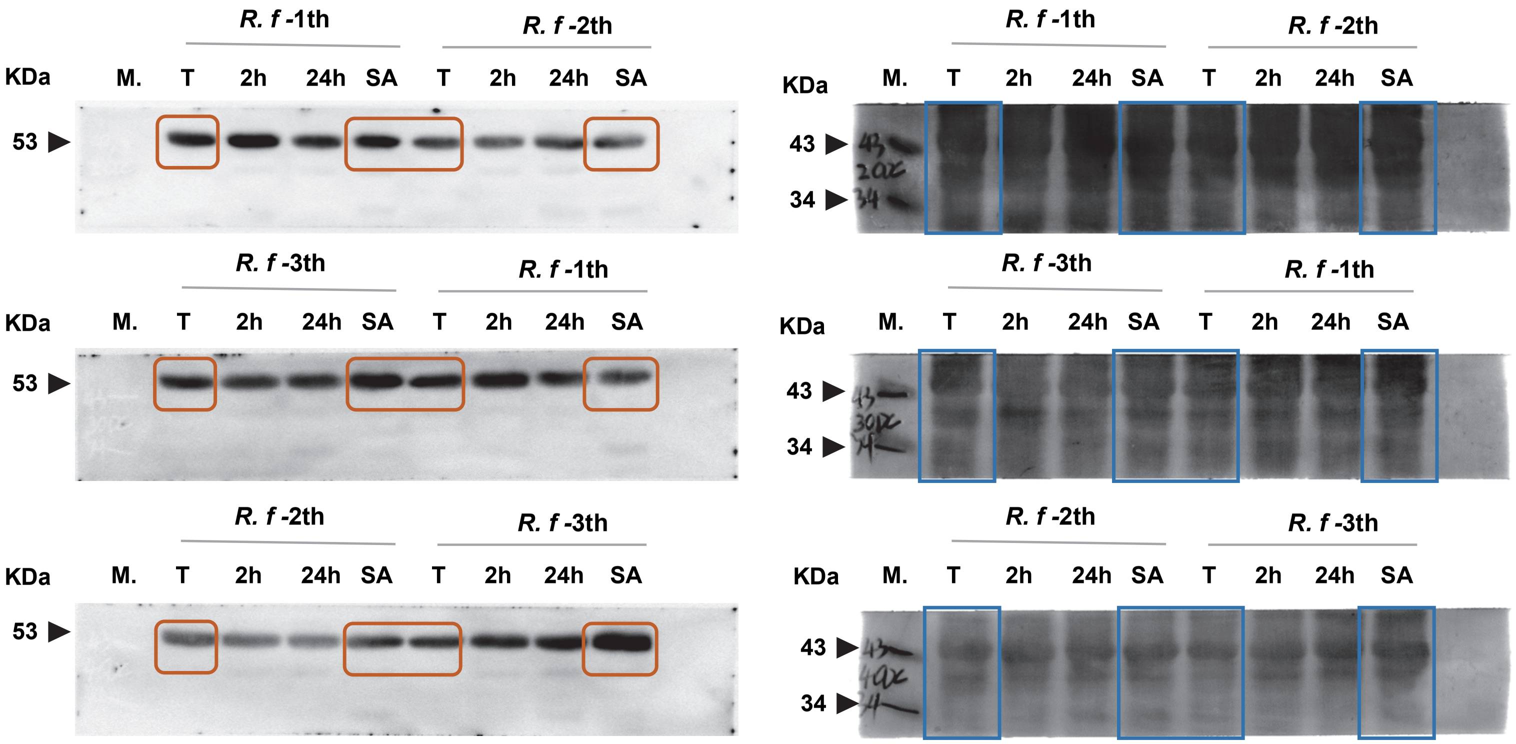

R. f -1th
R. f -2th
KDa
M. T 2h 24h SA T 2h 24h SA
53
R. f -3th
R. f -1th
KDa
M. T 2h 24h SA T 2h 24h SA
53
R. f -2th
R. f -3th
KDa
M. T 2h 24h SA T 2h 24h SA
53
R. f -1th
R. f -2th
KDa
M. T 2h 24h SA T 2h 24h SA
43
34
R. f -3th
R. f -1th
KDa
M. T 2h 24h SA T 2h 24h SA
43
34
R. f -2th
R. f -3th
KDa
M. T 2h 24h SA T 2h 24h SA
43
34

### AGMAT

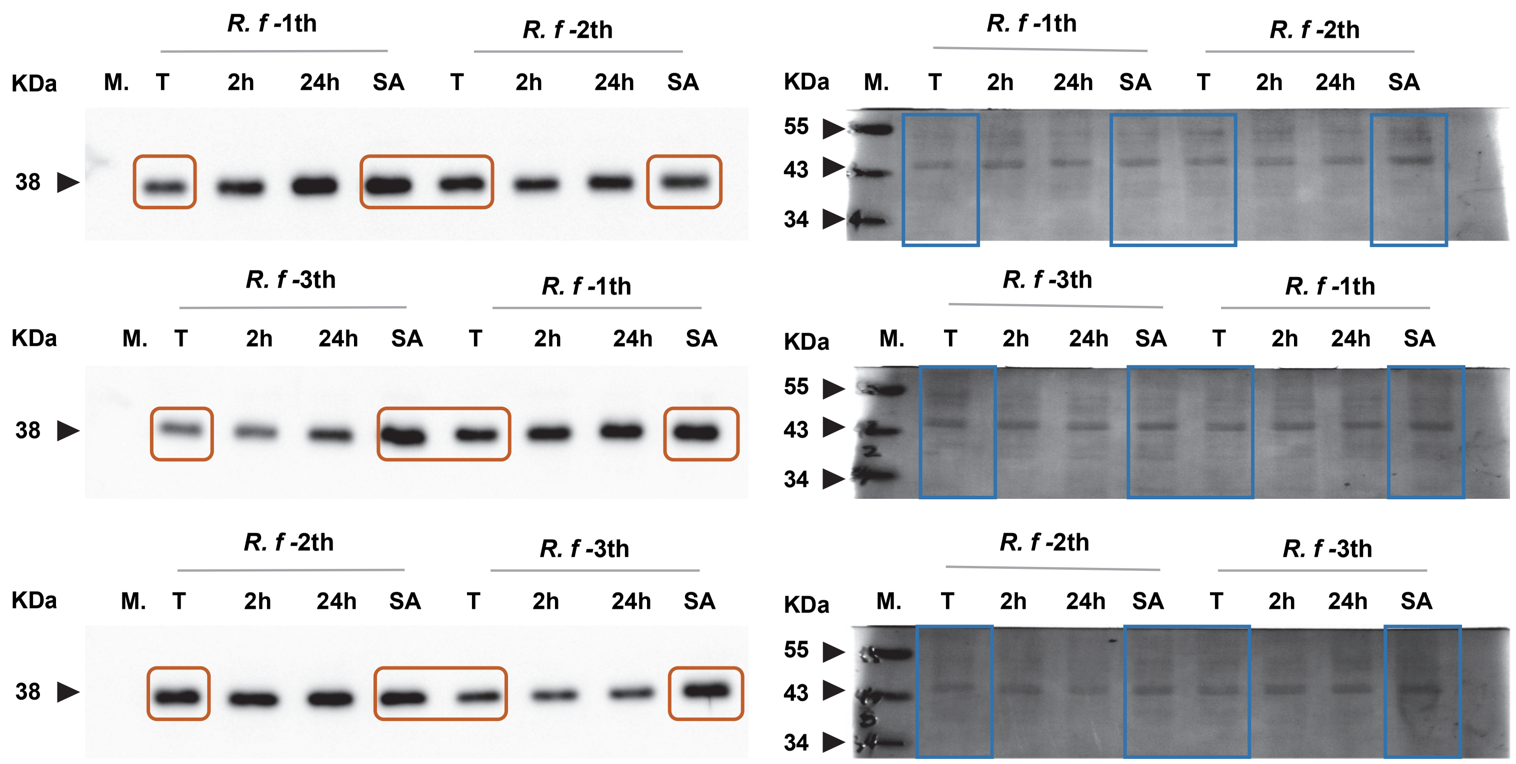
R. f -1th
R. f -2th
KDa
M.
T
2h
24h
SA
38
R. f -3th
R. f -1th
R. f -2th
R. f -3th
55
43
34

### AGMAT

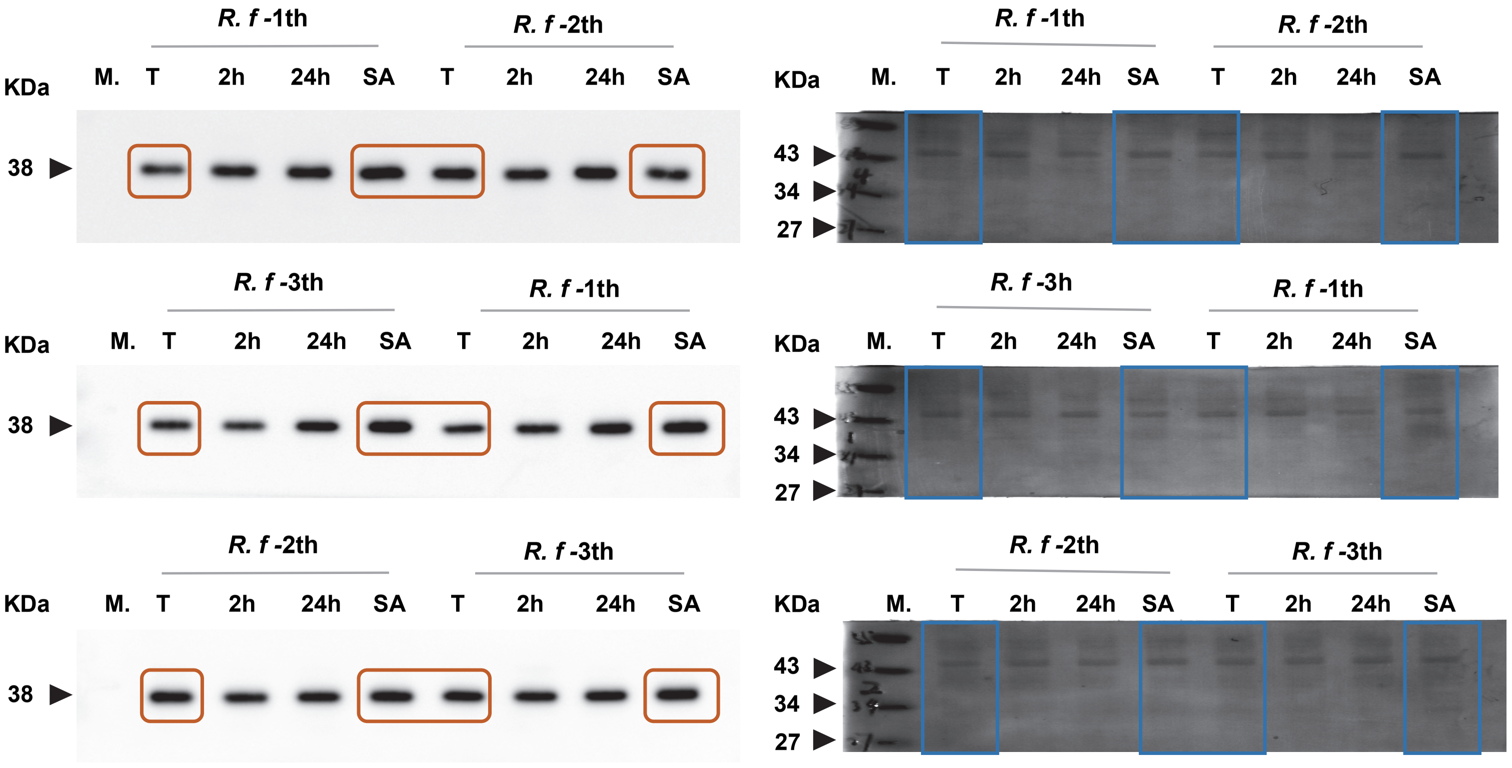

### NAGS

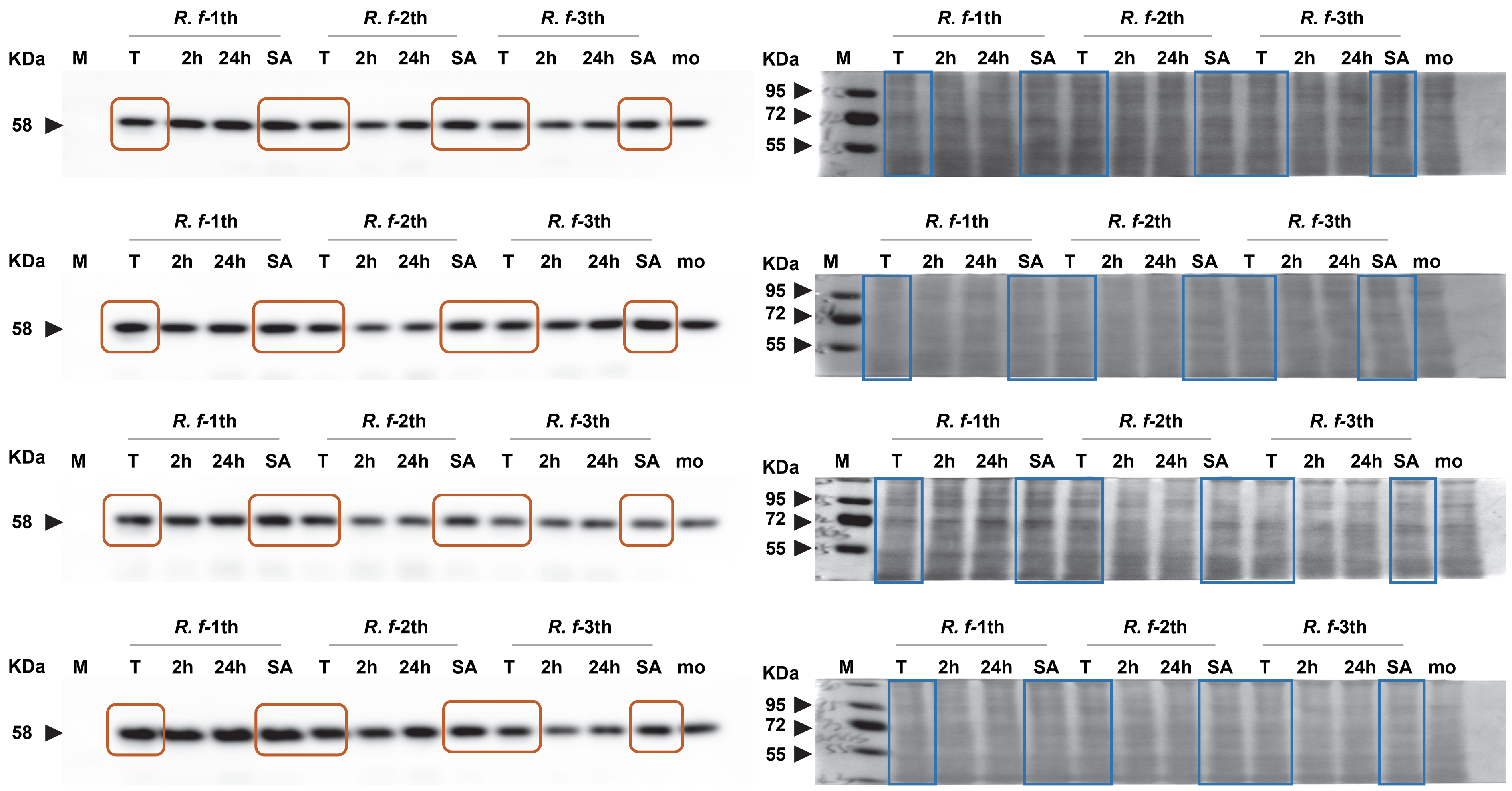

### NAGS

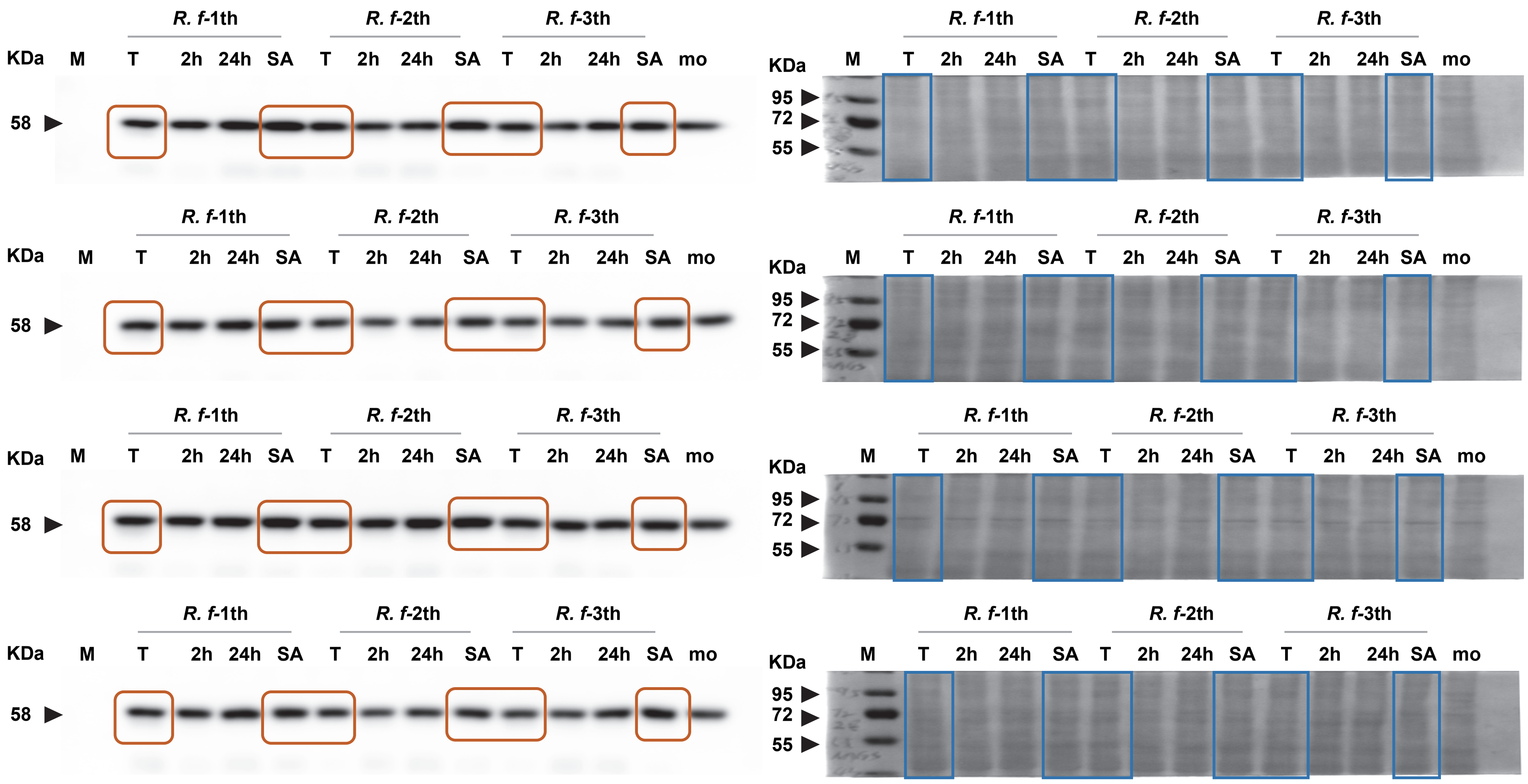

#### SIRT5

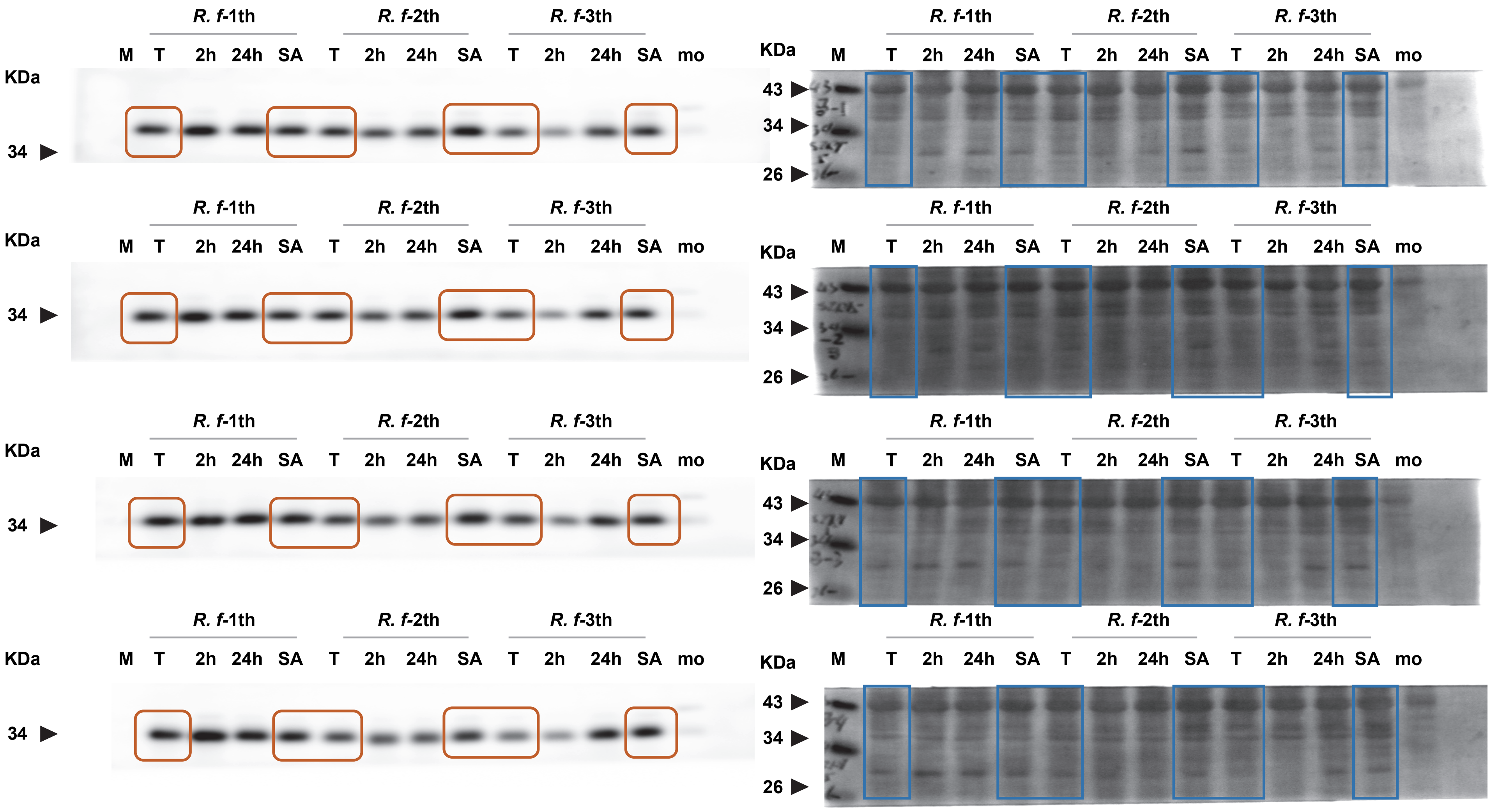
R. f-1th
R. f-2th
R. f-3th
M T 2h 24h SA T 2h 24h SA T 2h 24h SA mo
KDa
34
43
26

### SIRT5

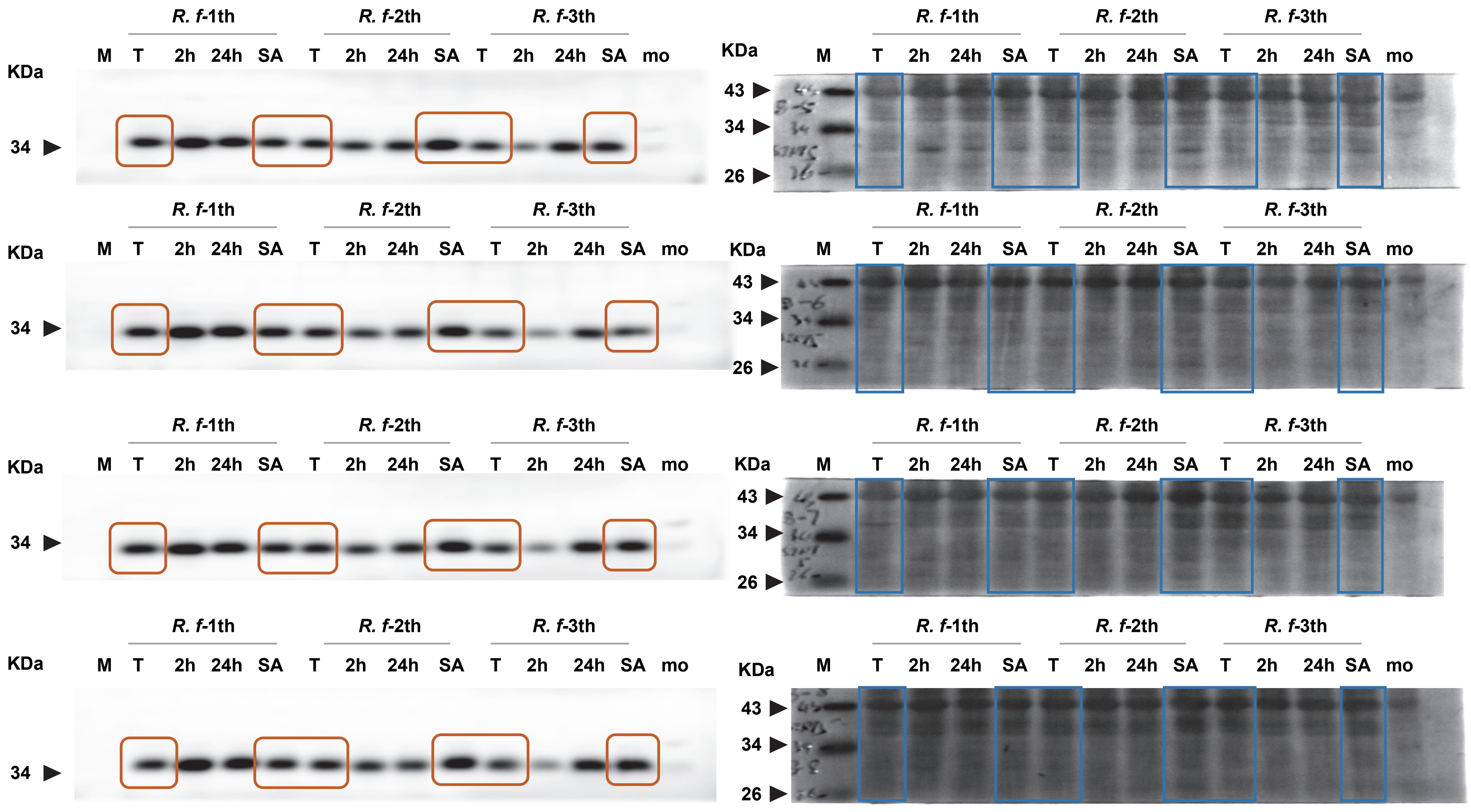
R. f-1th
R. f-2th
R. f-3th
M T 2h 24h SA T 2h 24h SA T 2h 24h SA mo
KDa
43
34
26
