## Supplementary Information 2 for "Interaction of Carbamoyl-Phosphate Synthase 1 with Agmatinase in the Liver of Torpid Bats"

### *Myotis ricketti*-Torpor

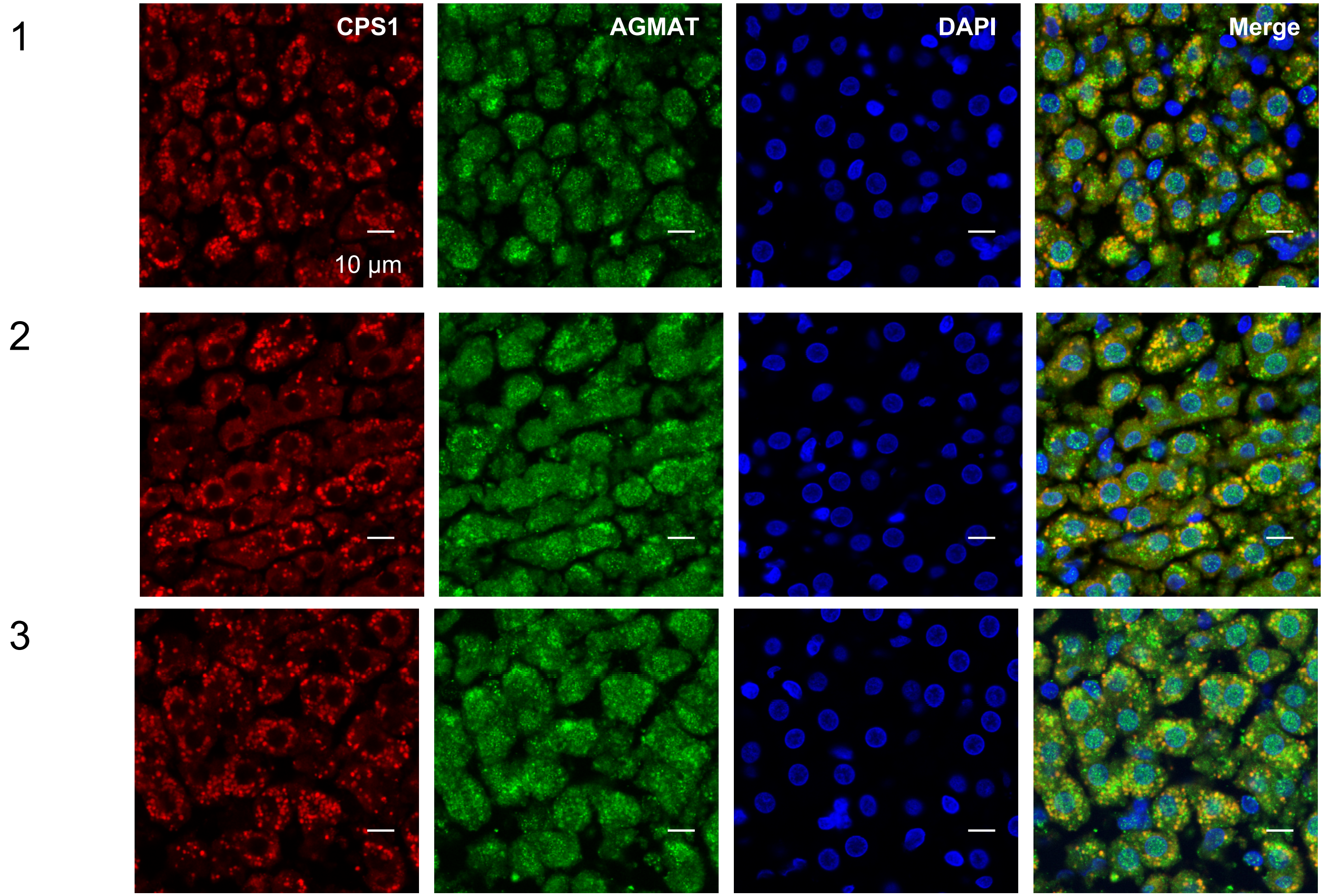


| | PCC | M1 | M2 |
|---|---|---|---|
| **1** | 0.654 | 0.477 | 0.914 |
| **2** | 0.654 | 0.456 | 0.907 |
| **3** | 0.622 | 0.39 | 0.938 |

### *Myotis ricketti*-summer active

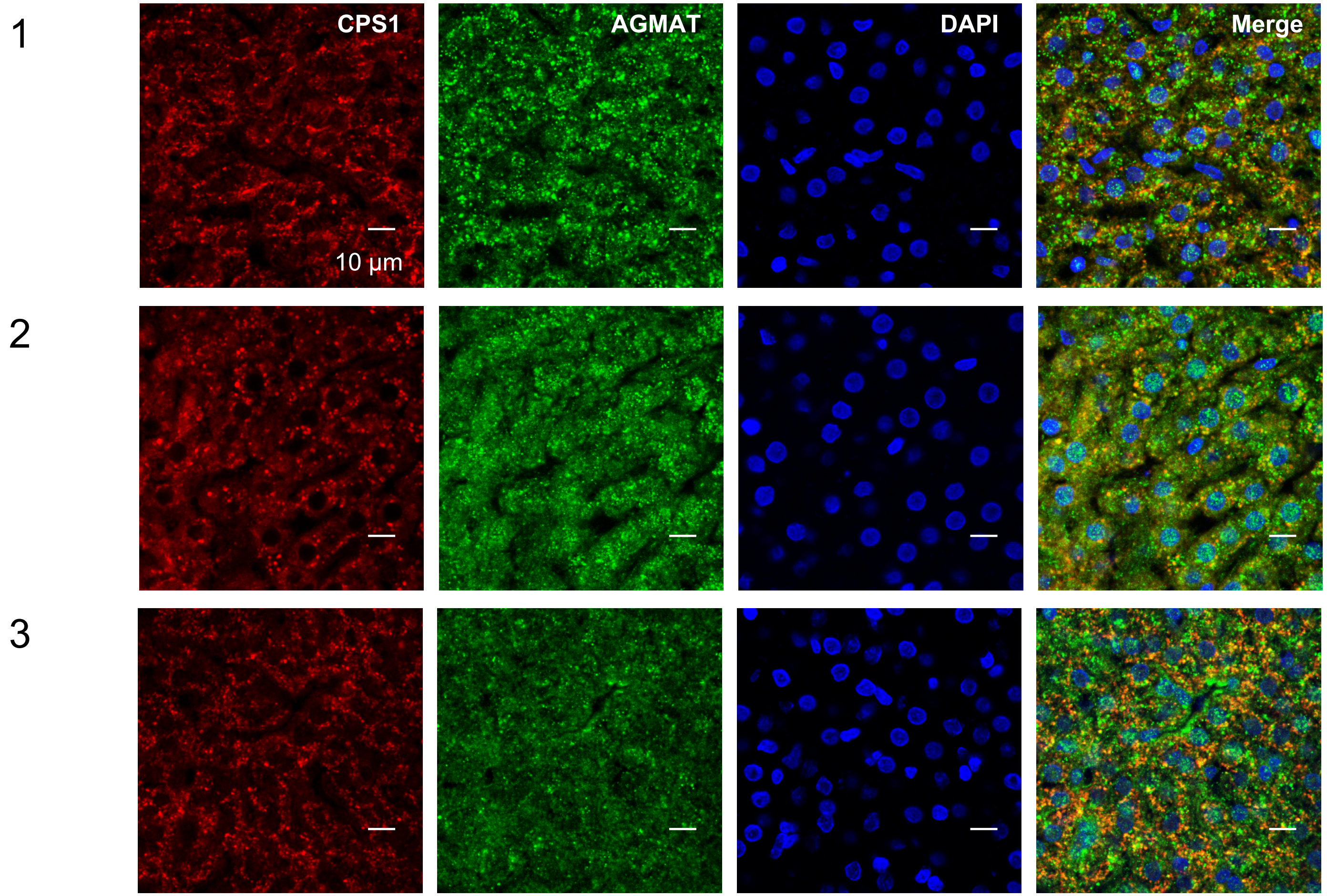


| | PCC | M1 | M2 |
|---|---|---|---|
| **1** | 0.414 | 0.51 | 0.585 |
| **2** | 0.399 | 0.505 | 0.738 |
| **3** | 0.353 | 0.335 | 0.706 |

### *Rhinolophus ferrumequinum* – Torpor

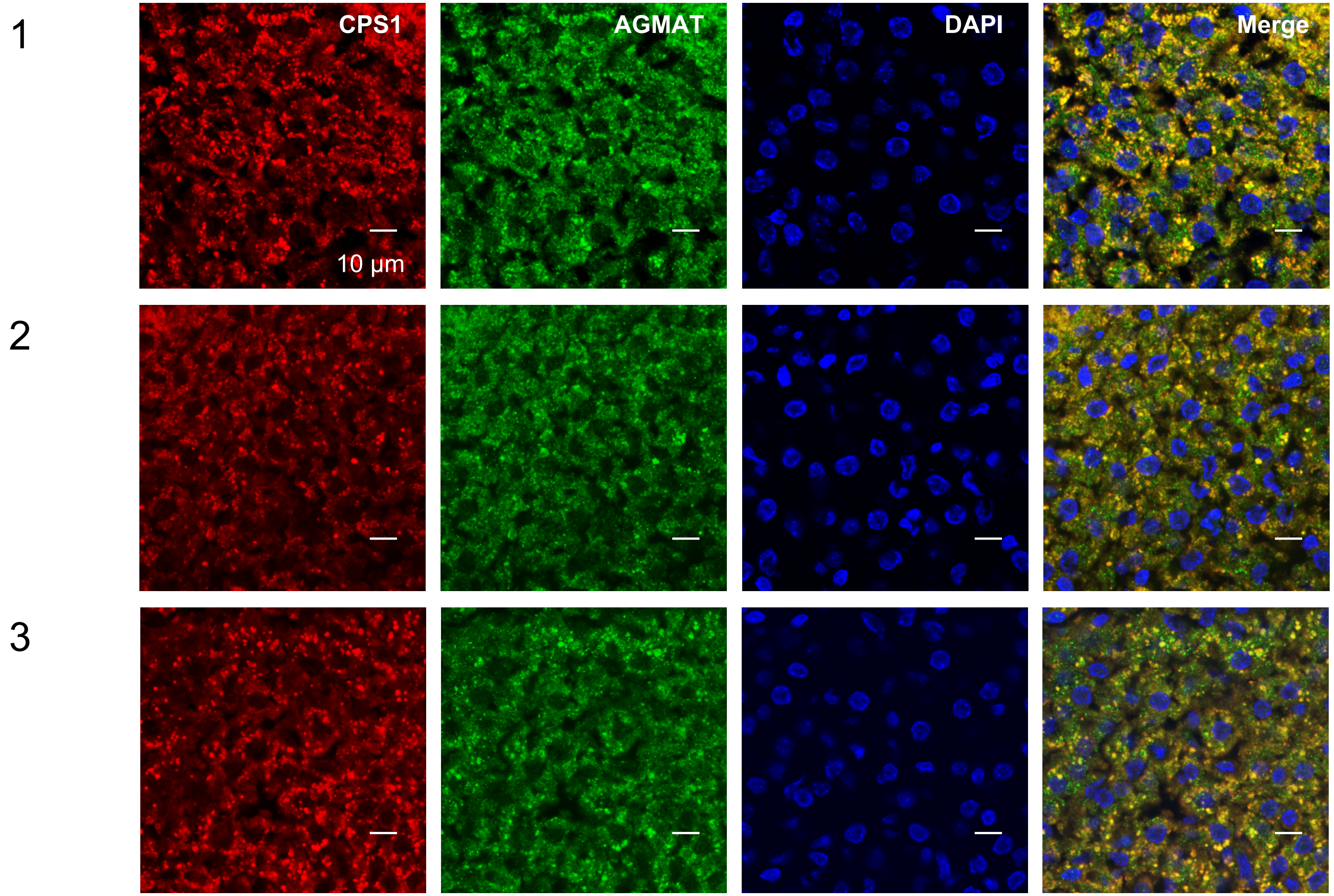


| | PCC | M1 | M2 |
|---|---|---|---|
| **1** | 0.805 | 0.577 | 0.942 |
| **2** | 0.73 | 0.516 | 0.922 |
| **3** | 0.767 | 0.499 | 0.929 |

### *Rhinolophus ferrumequinum* – summer active

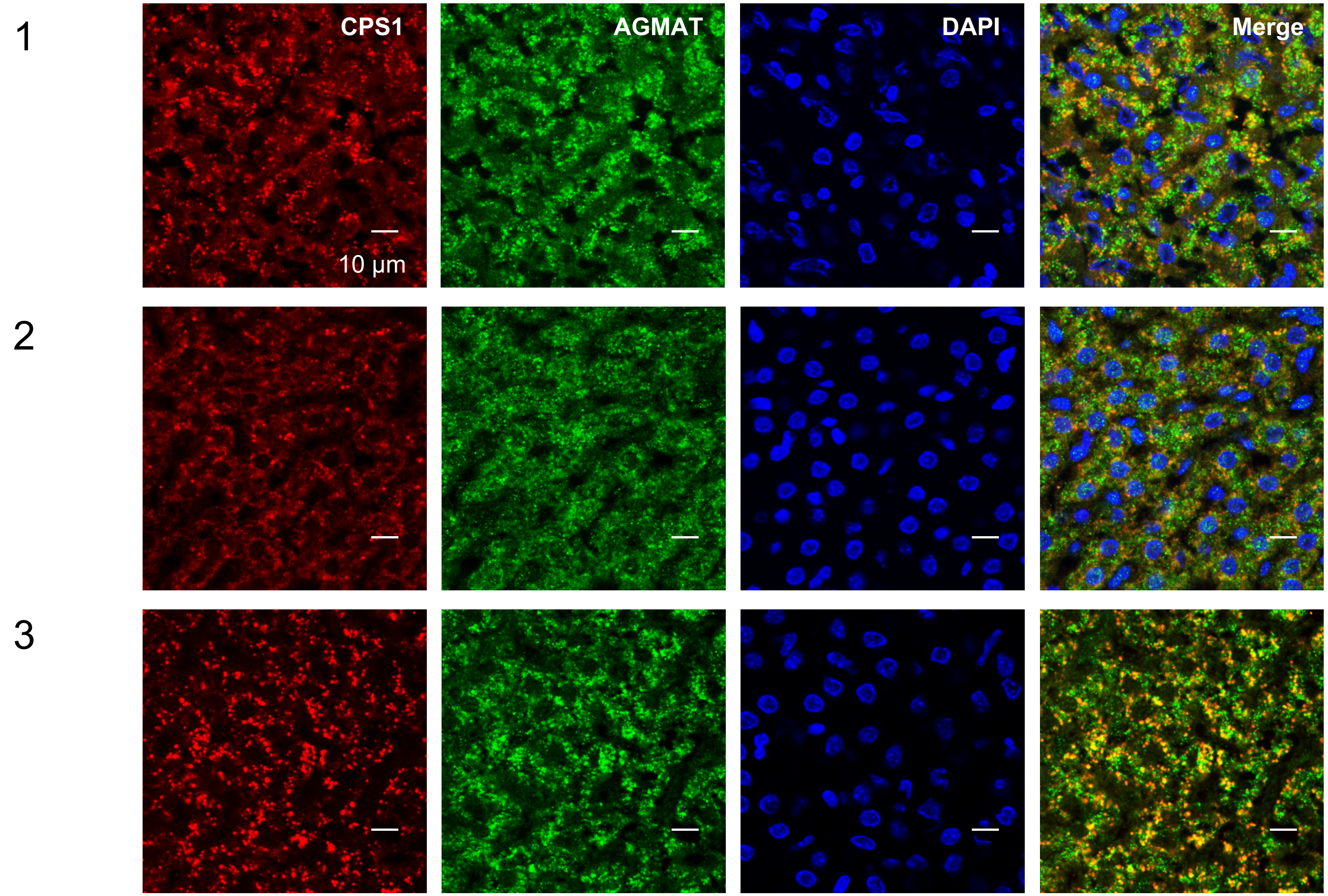


| | PCC | M1 | M2 |
|---|---|---|---|
| **1** | 0.493 | 0.516 | 0.707 |
| **2** | 0.501 | 0.561 | 0.744 |
| **3** | 0.619 | 0.421 | 0.816 |

### Mouse

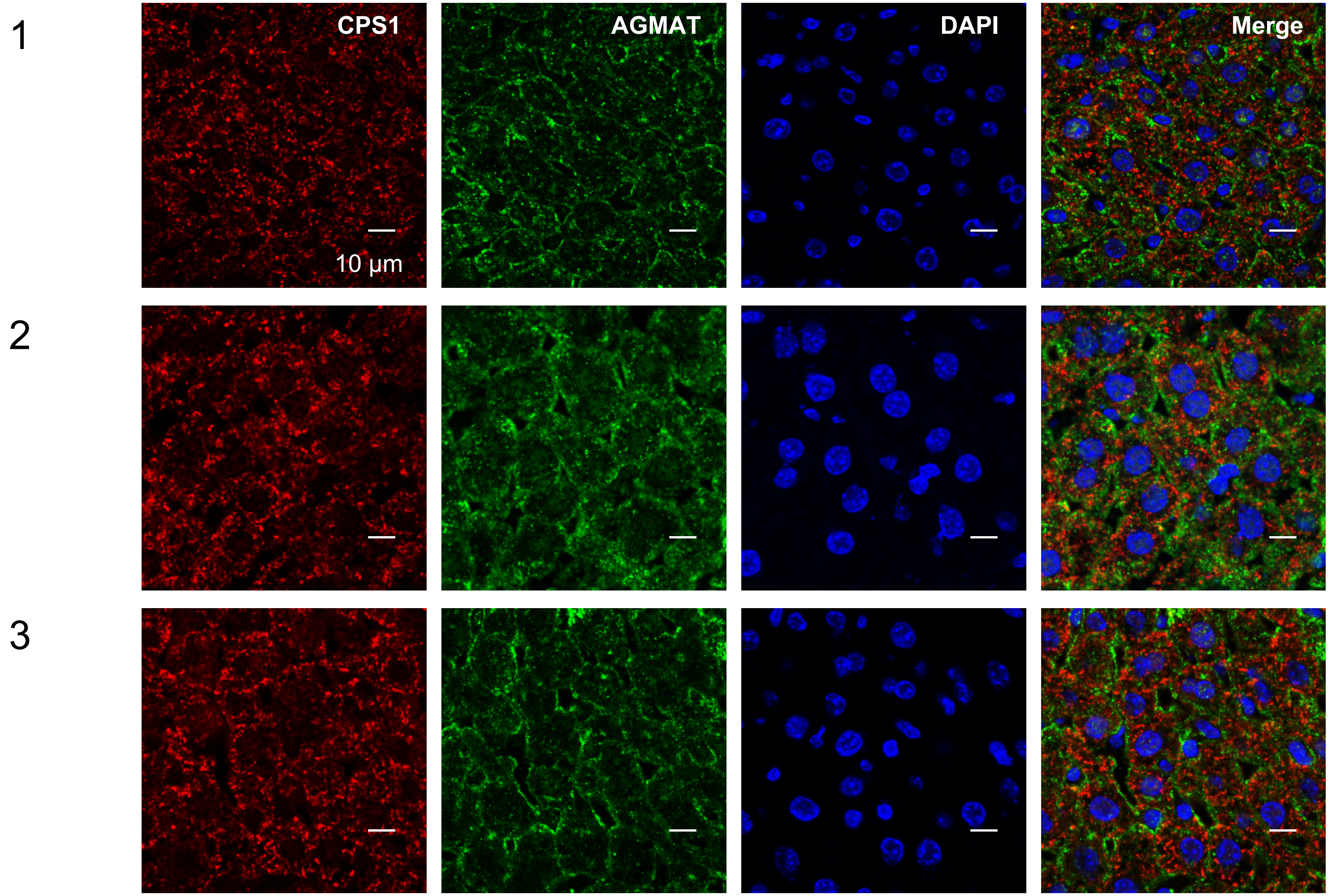


| | PCC | M1 | M2 |
|---|---|---|---|
| **1-1** | 0.134 | 0.27 | 0.363 |
| **2-1** | 0.164 | 0.305 | 0.264 |
| **3-1** | 0.178 | 0.268 | 0.398 |
