## Supplementary Information 3 for "Interaction of Carbamoyl-Phosphate Synthase 1 with Agmatinase in the Liver of Torpid Bats"

2021

### *Myotis ricketti*

#### Torpor – Individual 1

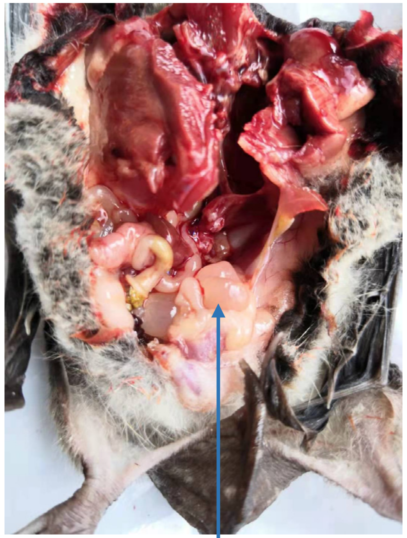

Bladder

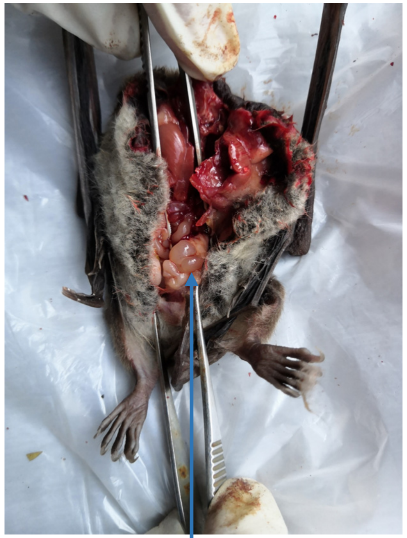

Bladder

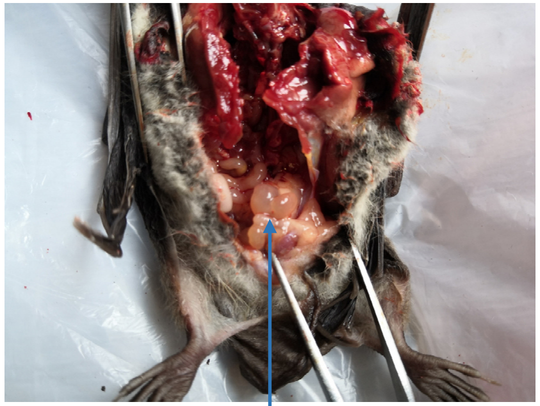

Bladder

#### Torpor – Individual 2

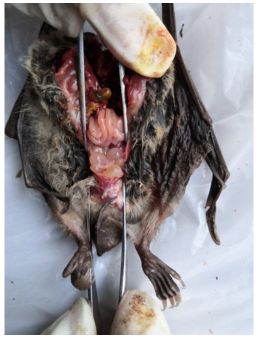
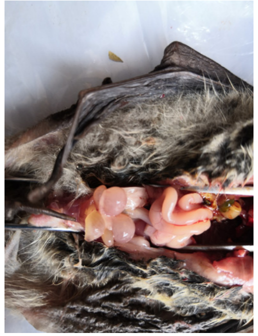
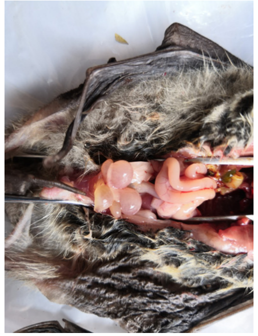
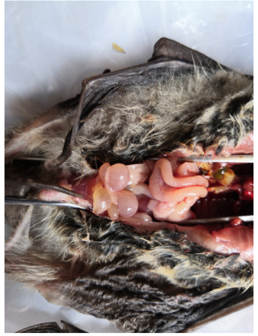
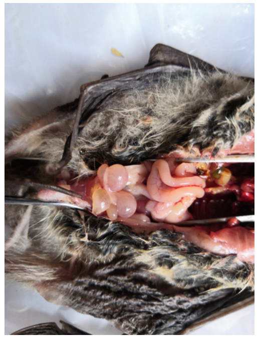

#### Torpor – Individual 3

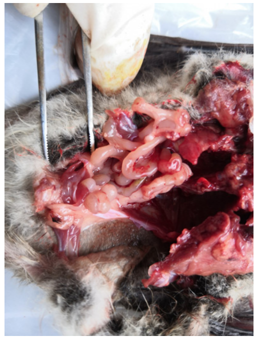
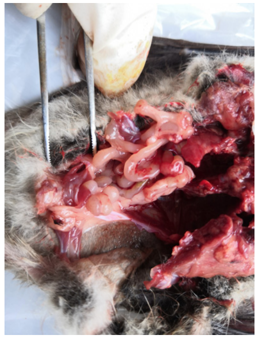
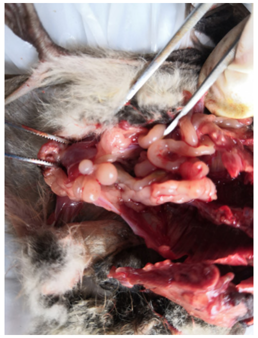

### *Myotis ricketti*

#### 2 h after arousal – Individual 1

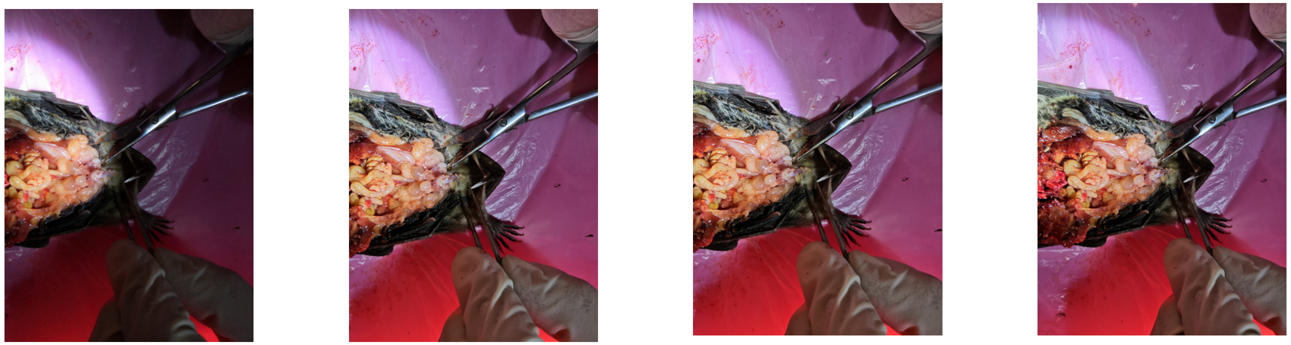

#### 2 h after arousal – Individual 2

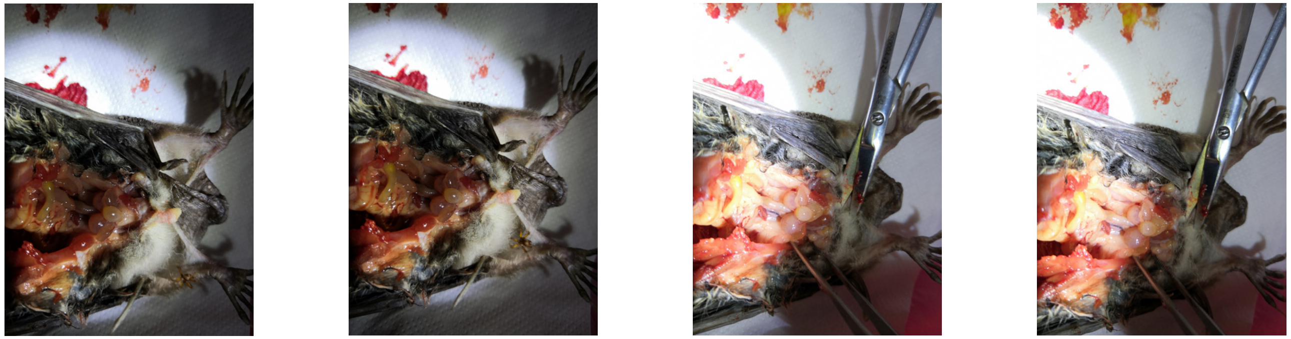

#### 2 h after arousal – Individual 3

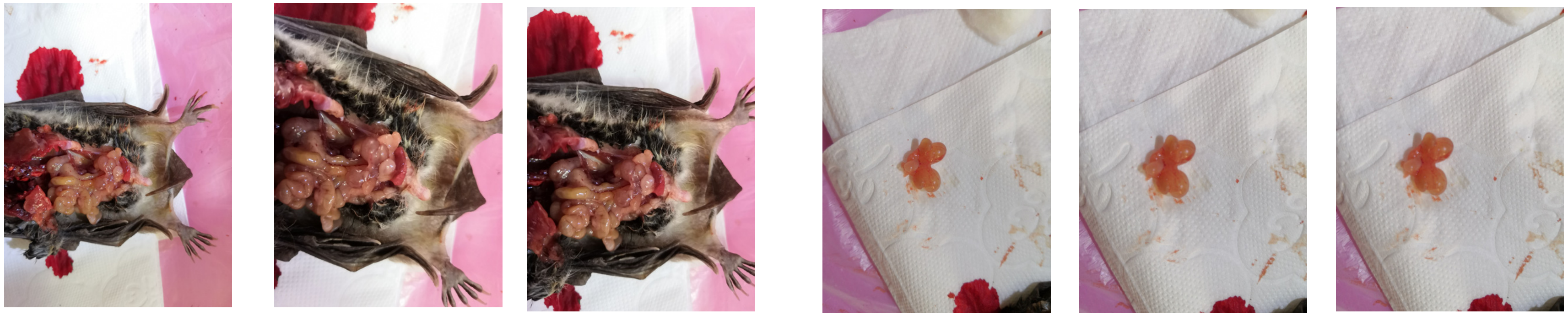

#### 24 h after arousal – Individual 1

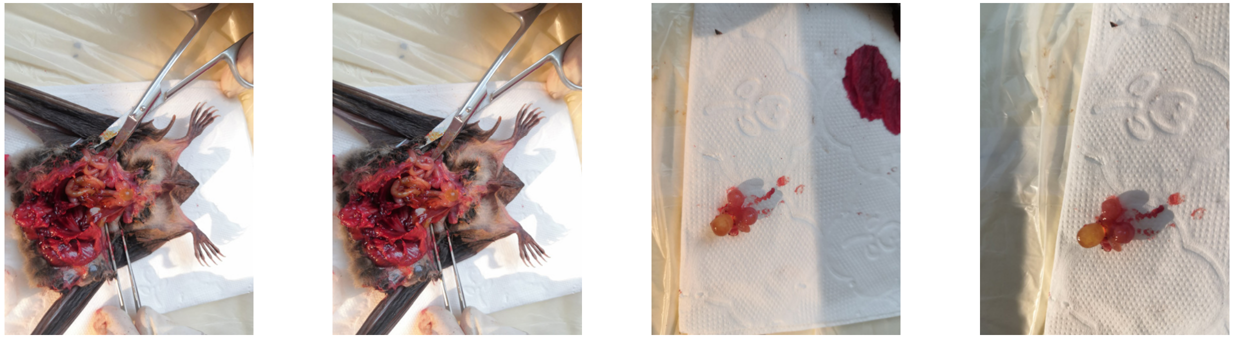

#### 24 h after arousal – Individual 2

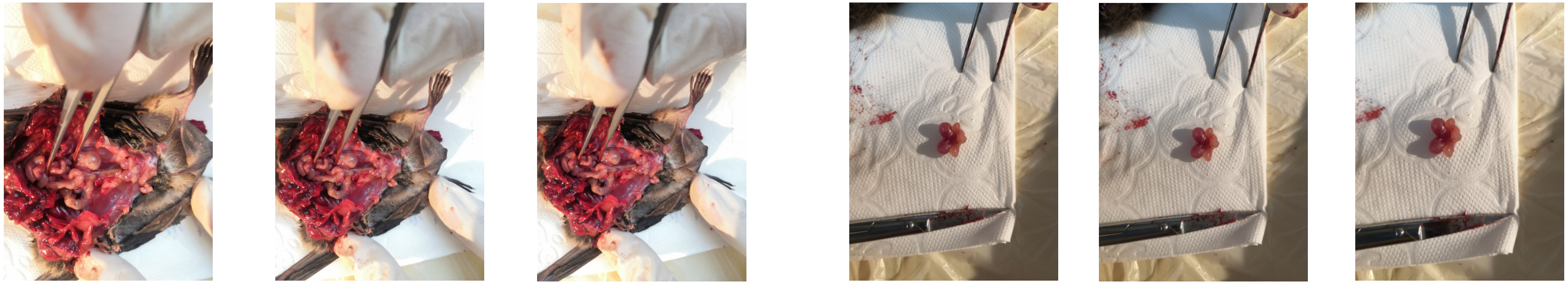

### *Myotis ricketti*

#### 24 h after arousal – Individual 3

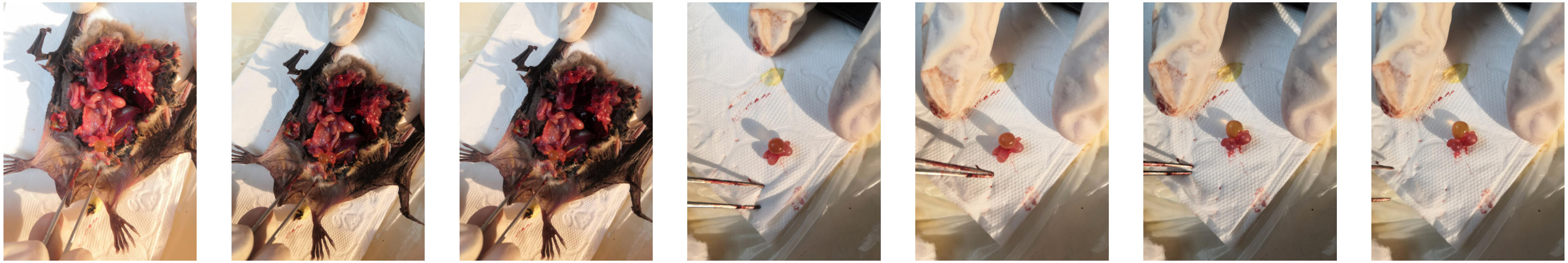

#### 24 h after arousal – Individual 4

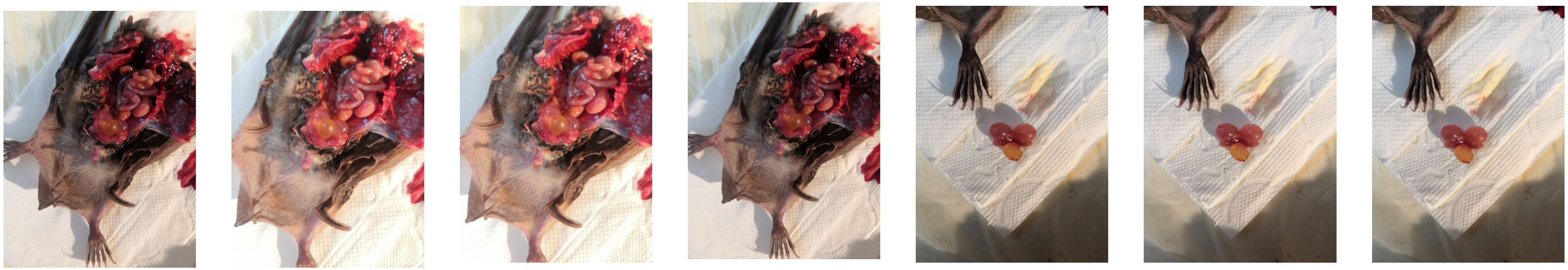

### *Myotis ricketti*

#### 24 h after arousal – Individual 5

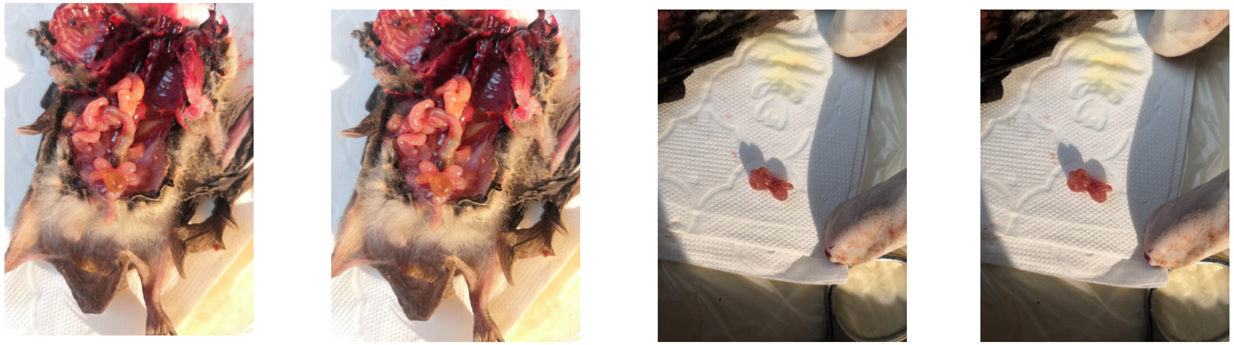

#### 24 h after arousal – Individual 6

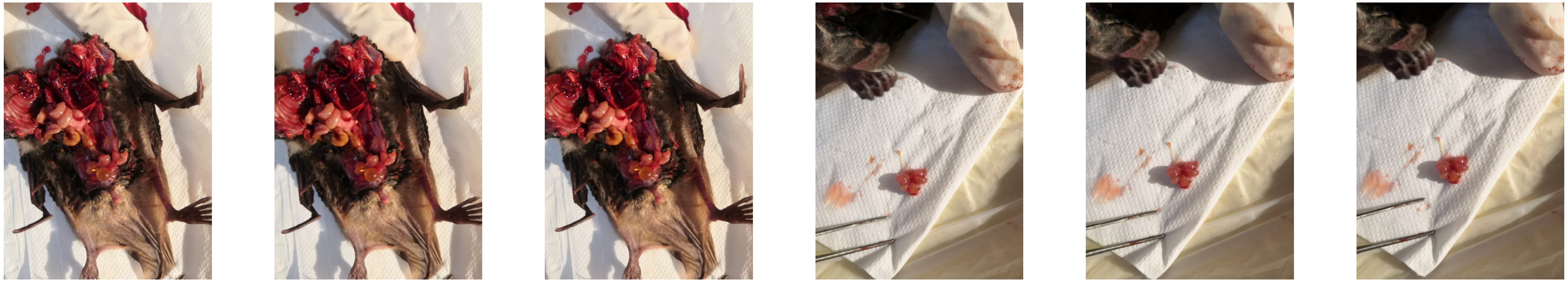

2021

### *Myotis ricketti*

#### Summer active – Individual 1

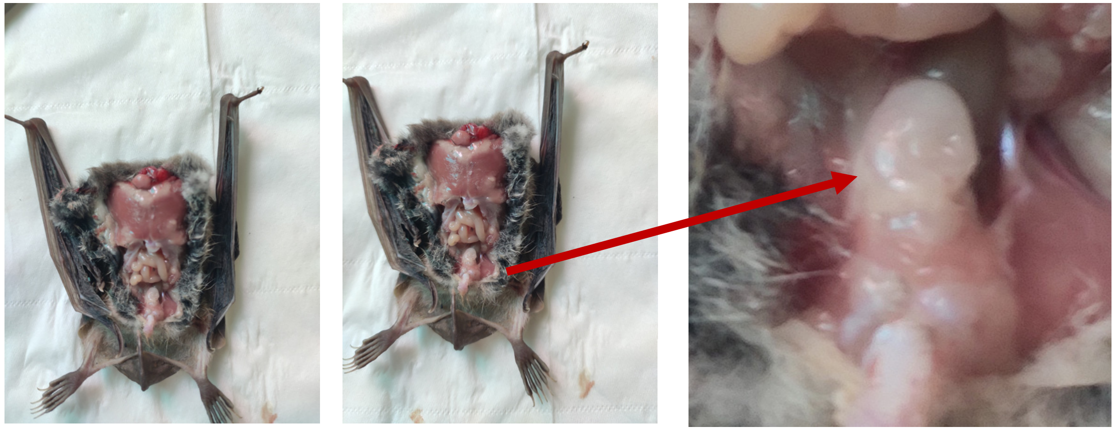

#### Summer active – Individual 2

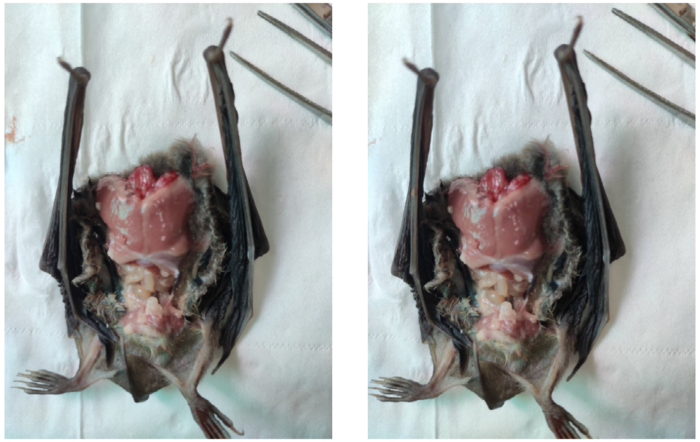

#### Summer active – Individual 3

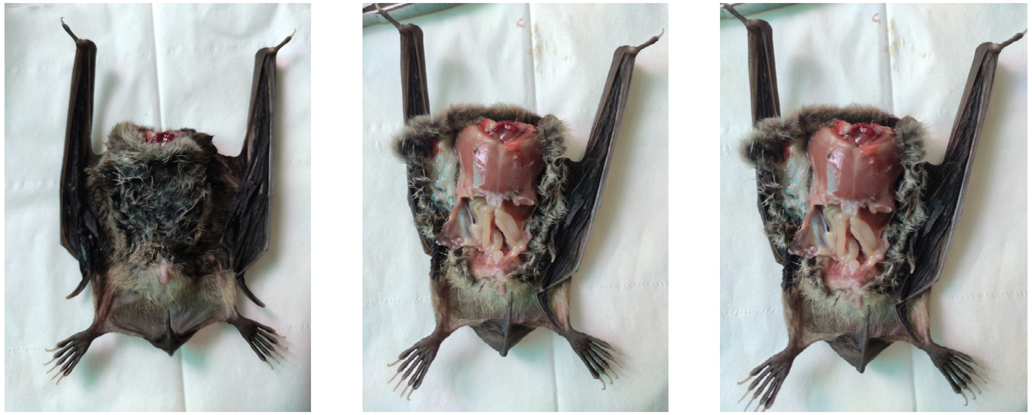

#### Summer active – Individual 4

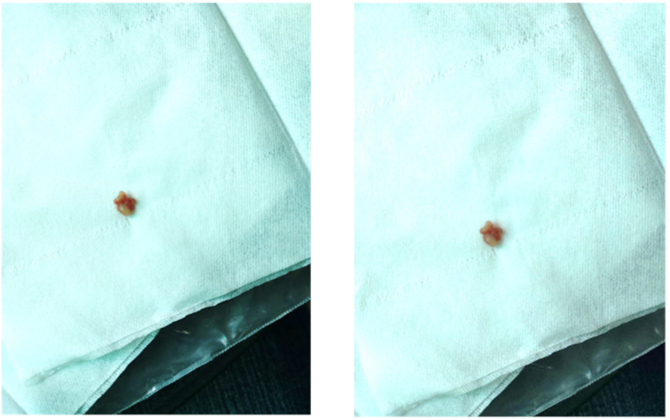

### *Myotis ricketti*

#### Torpor – Individuals

1

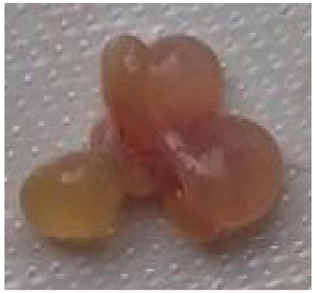

2

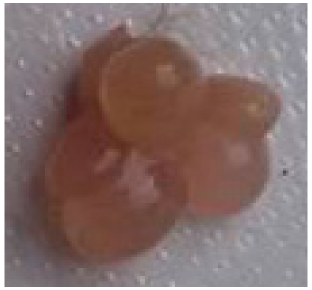

3

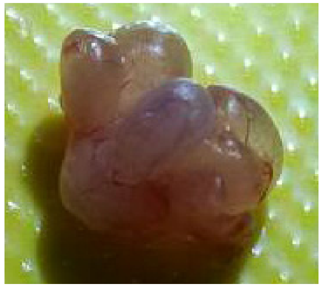

4

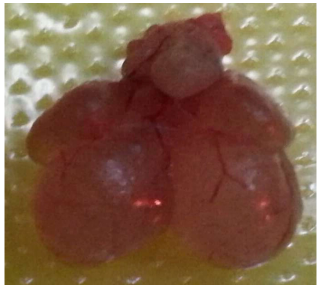

2021

### *Rhinolophus ferrumequinum*

#### Torpor – Individual 1

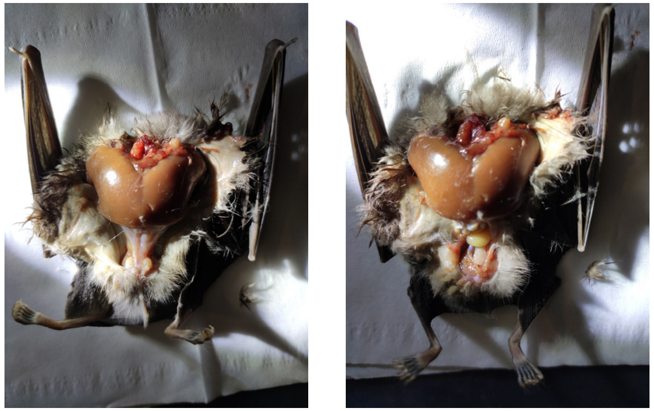

#### Torpor – Individual 2

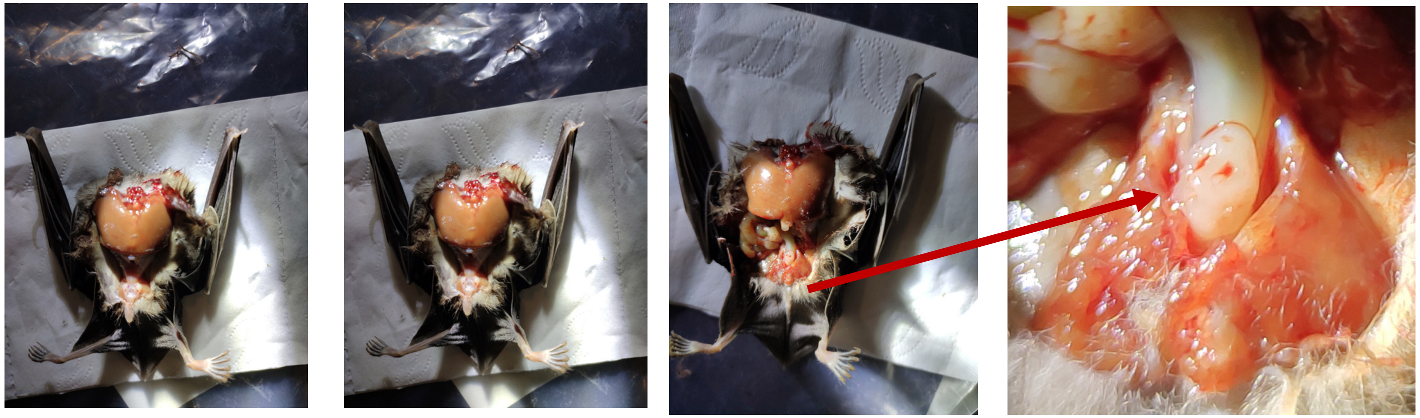

### *Rhinolophus ferrumequinum*

#### 2 h – Individual 1

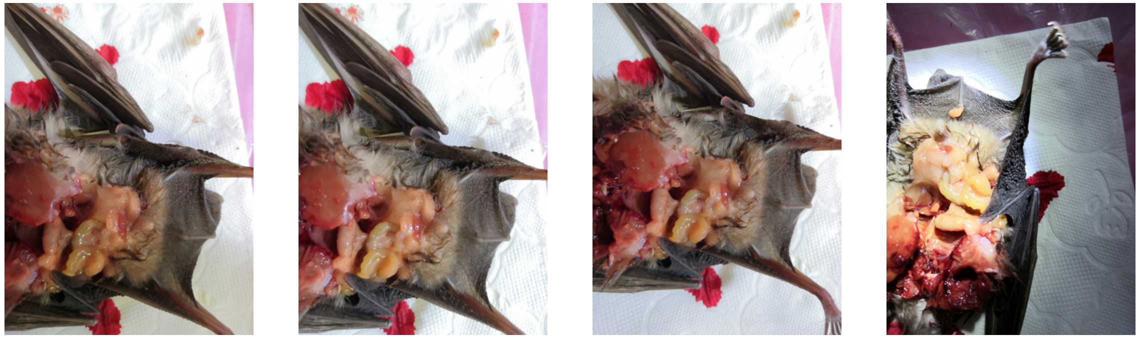

#### 2 h – Individual 2

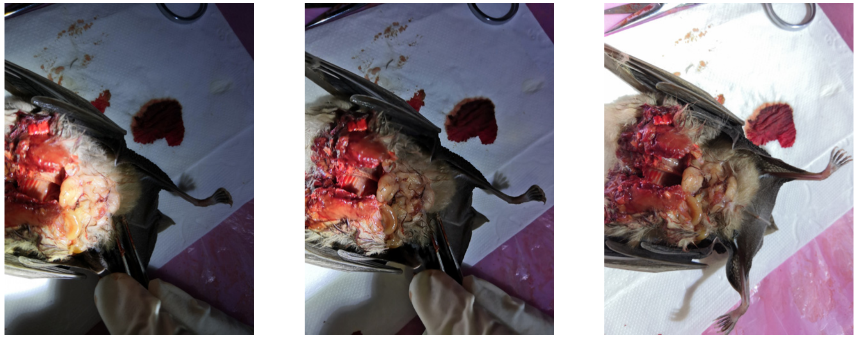

#### 2 h – Individual 3

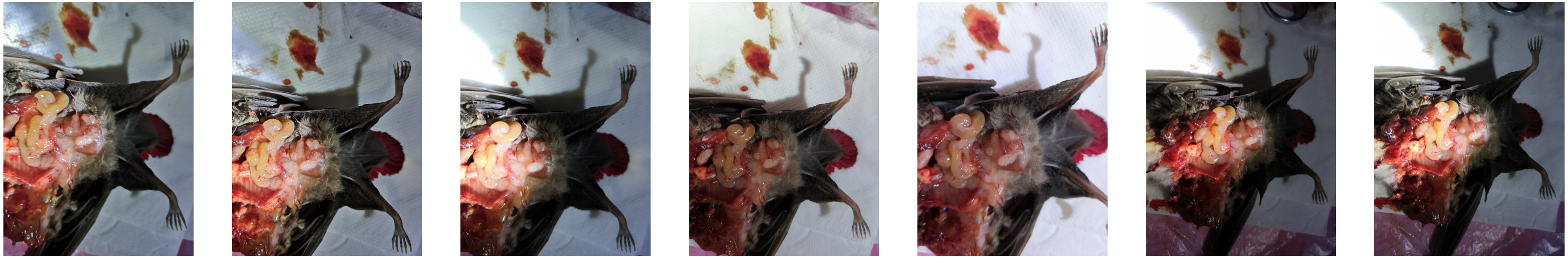

2021

### *Rhinolophus ferrumequinum*

**24 h – Individual 1**

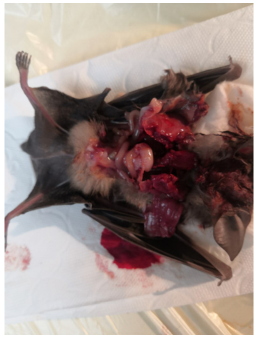

**24 h – Individual 2**

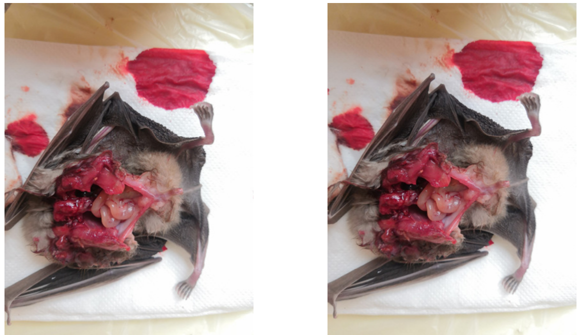

**Summer active – Individual 1**

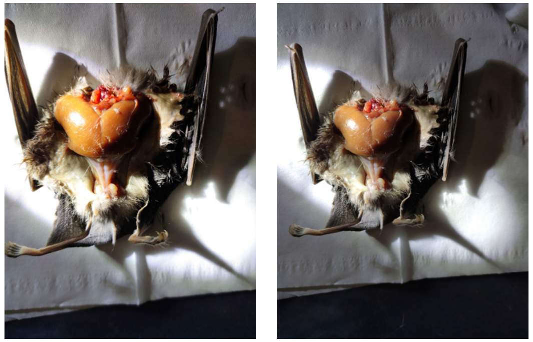

**Summer active – Individual 2**

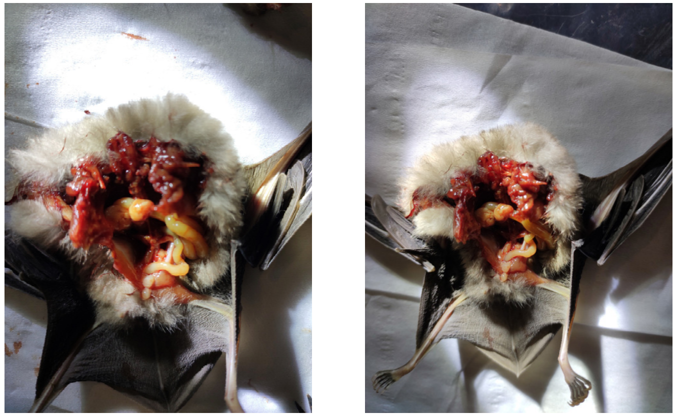

**Summer active – Individual 3**

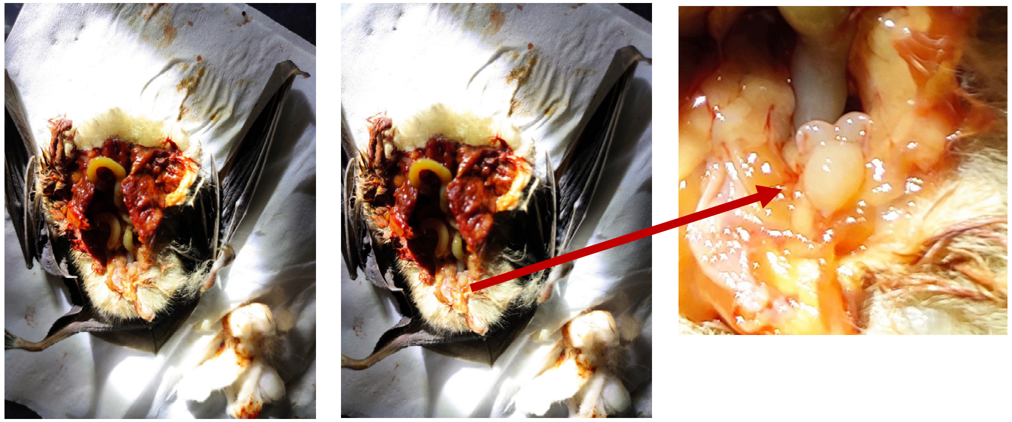

### *Mus musculus*

#### Mouse – Individual 1

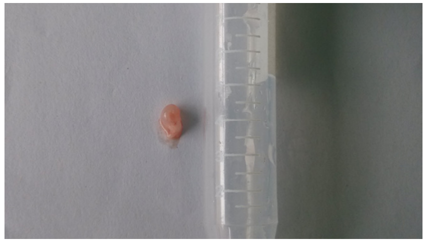

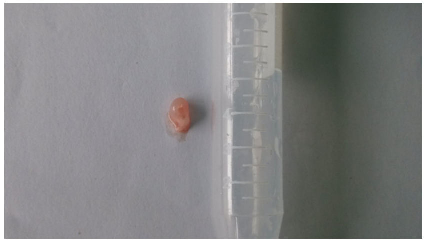

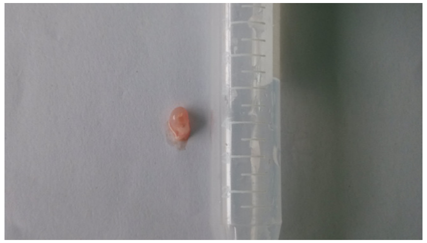

#### Mouse – Individual 2

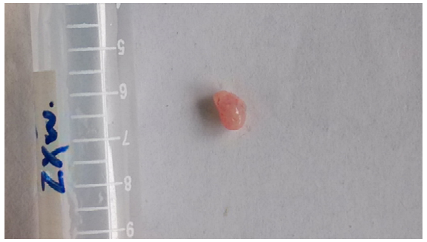

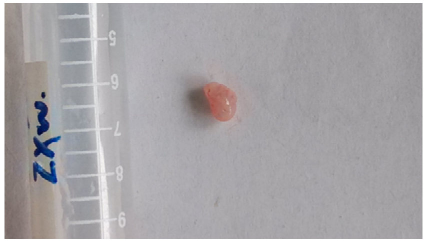

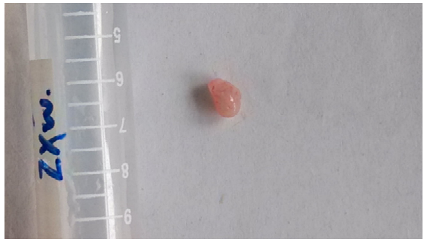

#### Mouse – Individual 3

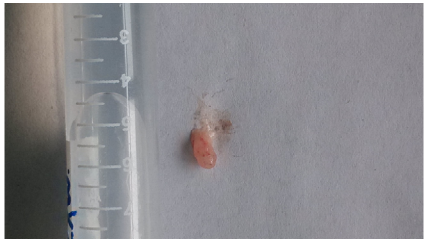

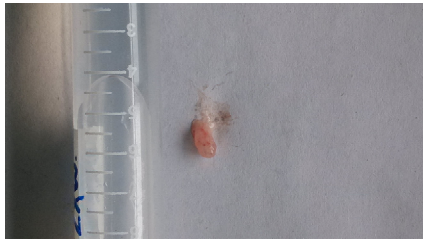

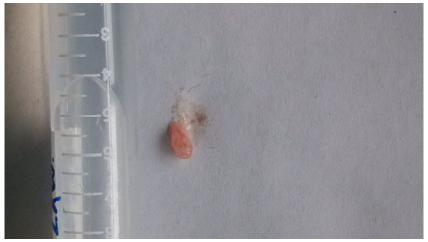

### *Mus musculus* (*db/db* mice)

**Mouse – Individual 1**

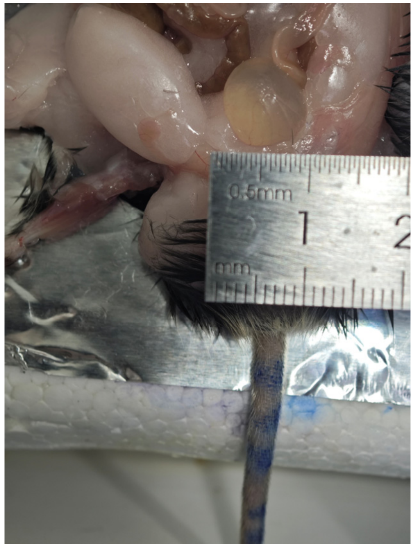


**Mouse – Individual 2**

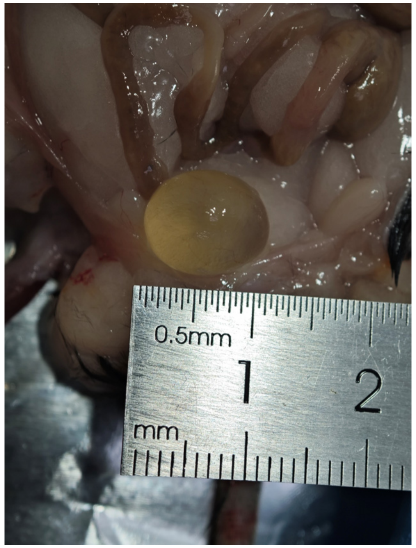


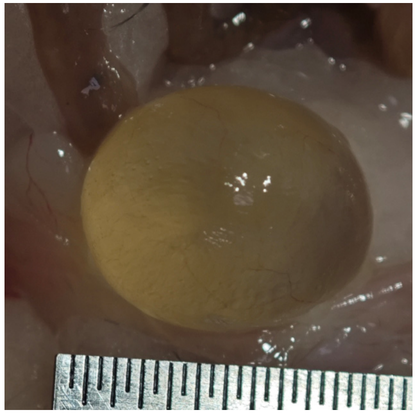

**Mouse – Individual 3**

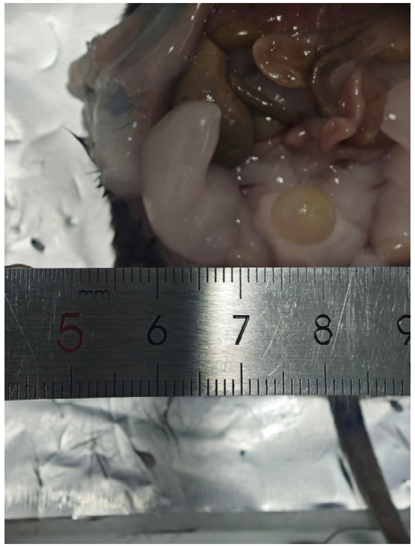
